# The shape of fitness functions and the distribution of mutational effect sizes jointly limit adaptation by regulatory mutations

**DOI:** 10.1101/2025.09.09.675213

**Authors:** Simon Aubé, Alexandre K. Dubé, Christian R. Landry

## Abstract

Mutations in gene regulatory regions have been shown to play a role in rapid adaptation but the factors determining their contribution are largely unknown. Using the yeast metabolic enzyme cytosine deaminase, we examine if adaptation to 5-fluorocytosine (5-FC), which requires reduced cytosine deamination and can readily arise from amino acid substitutions, may be reached by single promoter mutations. We generated all single-nucleotide substitutions and indels in the *FCY1* promoter and assayed the resulting mutants in presence of 5-FC. This revealed that no promoter mutation is sufficient for adaptation to occur. We next investigated how this inaccessibility of adaptation arises by combining large-scale expression measurements with the experimental characterization of the corresponding expression-fitness function. These experiments showed that the shape of this function precludes single promoter mutations from being adaptive. Although 24% of mutations significantly affect expression, the fitness curve is flat around wild-type level. As such, adaptation can only emerge from a severe reduction of expression, which cannot occur from a single mutation in the promoter. Our results show that the contribution of regulatory mutations to rapid adaptation not only depends on the distribution of mutational effect sizes on expression level but also on the shape of the function linking fitness to expression levels.

## Introduction

For rapid adaptation to occur, advantageous phenotypes must be immediately available through mutation. Whether a given mutation is adaptive depends on how the corresponding molecular change translates into a fitness effect. For expression changes, this is defined by the affected gene’s fitness function (**Fig. 1a**) – describing the relationship linking expression level to organismal fitness –, of which a diversity of shapes have been reported^1^. As such, the corresponding distribution of mutational effects (DME) on expression level and the shape of the relevant fitness function jointly dictate if adaptive changes can arise within a given cis-regulatory region. These two factors thus have a major impact on the available pathways to adaptation, as well as on the rate and mode of regulatory evolution. Adaptation would for instance be unlikely to arise from the combination of a narrow DME, such that mutations have small effects on expression, with a step-like fitness function where only large expression changes are beneficial. Conversely, single-nucleotide mutations sampled from the same narrow DME could be adaptive if the fitness function instead displayed more gradual variation, allowing small expression changes to provide a benefit (**Fig. 1b**).

**Fig. 1:**
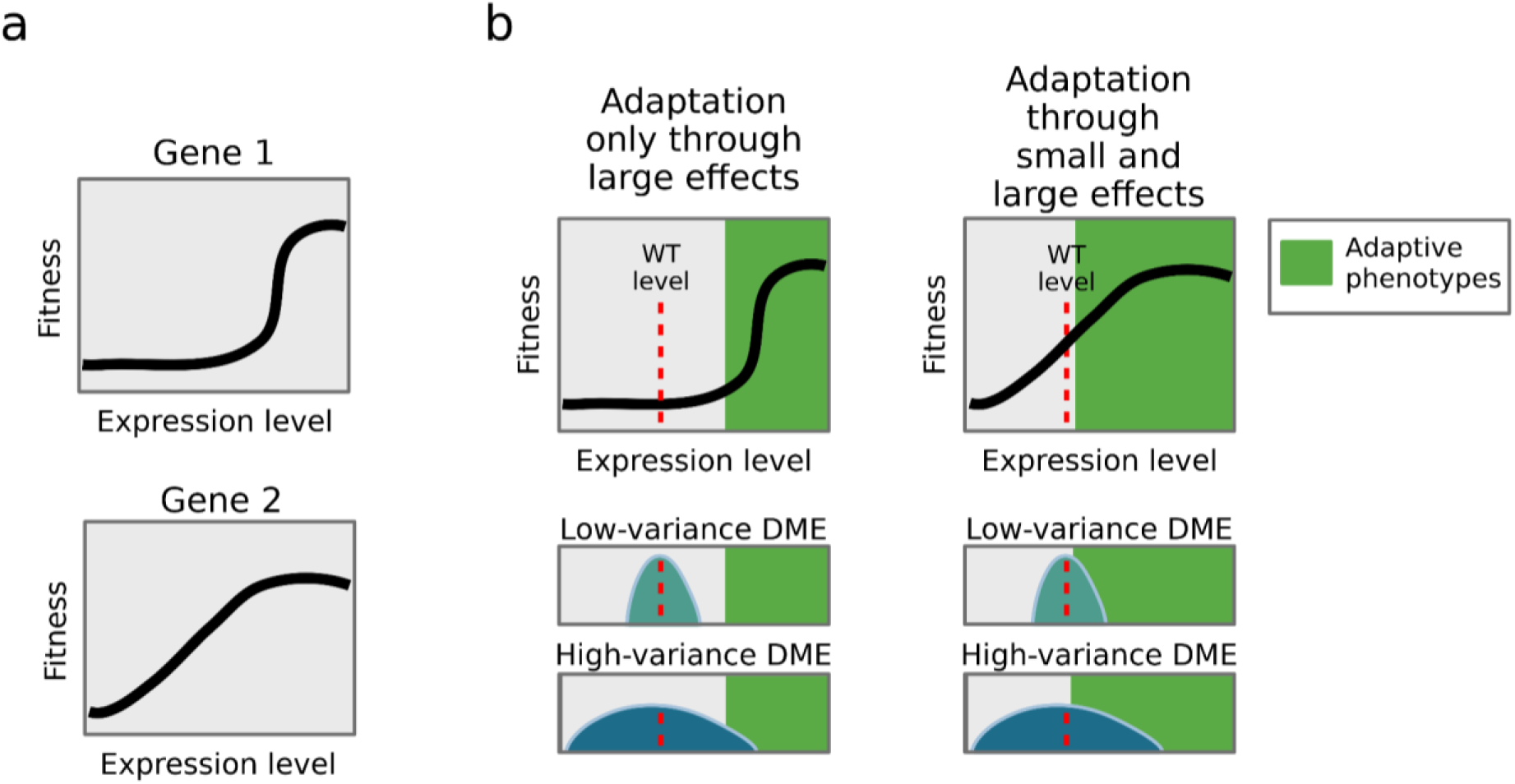
The DME and the shape of the fitness function jointly define the accessibility of adaptation. **a,** Fitness functions describe the relationship between the expression level of a gene and organismal fitness. Hypothetical fitness functions are shown for two genes. In each case, the shape of the curve displays how fitness varies with increasing expression level of the corresponding gene. **b,** The accessibility of adaptation through expression level changes depends on mutational effect sizes and fitness function shape. For the two fitness functions, adaptation from the same low-fitness wild-type (WT) expression level is shown. When the function is flat around WT expression (left), only large expression increases from the high-variance DME can be adaptive. In contrast, when the fitness function is curved near WT expression (right), small expression increases are adaptive, such that adaptation is accessible from the low-variance DME as well as the high-variance one. Here, WT expression levels are shown to lie far from the high-fitness region for visual clarity. While such a situation is likely rare^1^, the existence of beneficial gene duplications shows that it is possible^29–34^. Created in BioRender. Aubé, S. (2025) https://BioRender.com/rw8qljy.

Fully understanding the evolution of cis-regulatory sequences and its contribution to adaptation hence requires assaying mutational effects on expression level and characterizing their conversion through fitness functions^2^. Previous large-scale studies have investigated the effect of mutations within promoters and cis-regulatory regions^3–9^ generally focused on expression level without directly measuring the corresponding fitness consequences. Other studies collectively assayed the effects of cis-regulatory mutations on fitness and mapped the corresponding fitness function, for a limited number of yeast genes: *TDH3* ^10–12^ and *SUL1* ^13,14^. As such, the interplay between the DME on expression level and the shape of the fitness function remains unknown for most genes, in spite of its prime importance for understanding the evolution of gene regulation ^2^

Across organisms, a major engine of rapid adaptation is loss-of-function (LOF) mutations, both in coding and regulatory sequences. In humans, the best known mechanisms of resistance to malaria are the loss of expression of specific genes in red blood cells^15,16^ and the partial loss of a key metabolic enzyme activity^17^. Among the best documented adaptive morphological changes in vertebrates in natural populations arose through the repeated loss of expression of a developmental gene^18^. LOF mutations in coding sequences are also a major pathway to antimicrobial resistance, for instance in fungal pathogens evolving in response to antifungal drugs^19^. Expression-decreasing epigenetic changes are involved in the adaptive evolution of cancer cells, by reducing expression of tumor suppressor genes^20^. Finally, laboratory microbial evolution has shown that adaptation occurs through LOF at least as frequently as through gain of function(s)^21–28^. LOF therefore provides a powerful model to investigate questions regarding the contributions of mutations to rapid adaptation.

We focused on the adaptive loss of Fcy1 activity in yeast exposed to prodrug 5-fluorocytosine (5-FC). While this enzyme’s canonical activity is cytosine deamination, it also converts 5-FC into 5-fluorouracil, which leads to growth arrest and cell death^35^. Accordingly, complete or partial LOF mutations in the *FCY1* coding sequence confer resistance and are adaptive^36^. To assess whether regulatory mutations can also contribute to rapid adaptation to this condition, we performed saturation mutagenesis of the *FCY1* promoter. We measured the effect of all possible single nucleotide substitutions and indels in this region on fitness and expression level. These experiments revealed no measurable fitness gain in medium with 5-FC, although a quarter of single-nucleotide changes significantly affect gene expression level. Through the characterization of the underlying fitness function, we show that this discrepancy emerges jointly from the molecular effect of mutations and from the shape of this relationship. Because the fitness function is flat around the endogenous expression level, adaptation can only occur through large expression decreases, which cannot arise from a single-nucleotide change in the promoter. These findings highlight how the availability of adaptative phenotypes through mutation can be restricted by the combined action of the DME and of the relevant fitness function. In *FCY1*, this results in rapid adaptation being only accessible through mutations in the coding sequence. This observation may be generalizable to other enzymes, especially those which exert low control over metabolic fluxes.

## Results

### Measuring the effects of mutations within the *FCY1* promoter

Because the growth of yeast *Saccharomyces cerevisiae* has been shown to decrease monotonically with increasing 5-FC concentrations^36^ (**Fig 2a**, **Extended Data Fig. 1**), we hypothesized that at least some mutations in the *FCY1* promoter would be adaptive, by acting similarly as a decrease in the concentration of 5-FC (**Fig 2a**). To identify promoter regions to mutate, we performed deletions of successive 77 bp fragments of the intergenic region upstream of the *FCY1* gene. Among four tested deletions, only that of the most proximal fragment alleviated the toxicity of 5-FC **(Extended Data Fig. 2**). For this reason, and because most regulatory elements in yeast promoters are found within the 150 bp closest to the transcribed gene^37,38^, we focused on the 154 bp immediately upstream of *FCY1*. Throughout, we defined nucleotide positions within this region relative to the start codon of *FCY1*, with position 0 assigned to the nucleotide immediately upstream of the gene and position −153 representing the nucleotide most distal from *FCY1*.

**Fig. 2:**
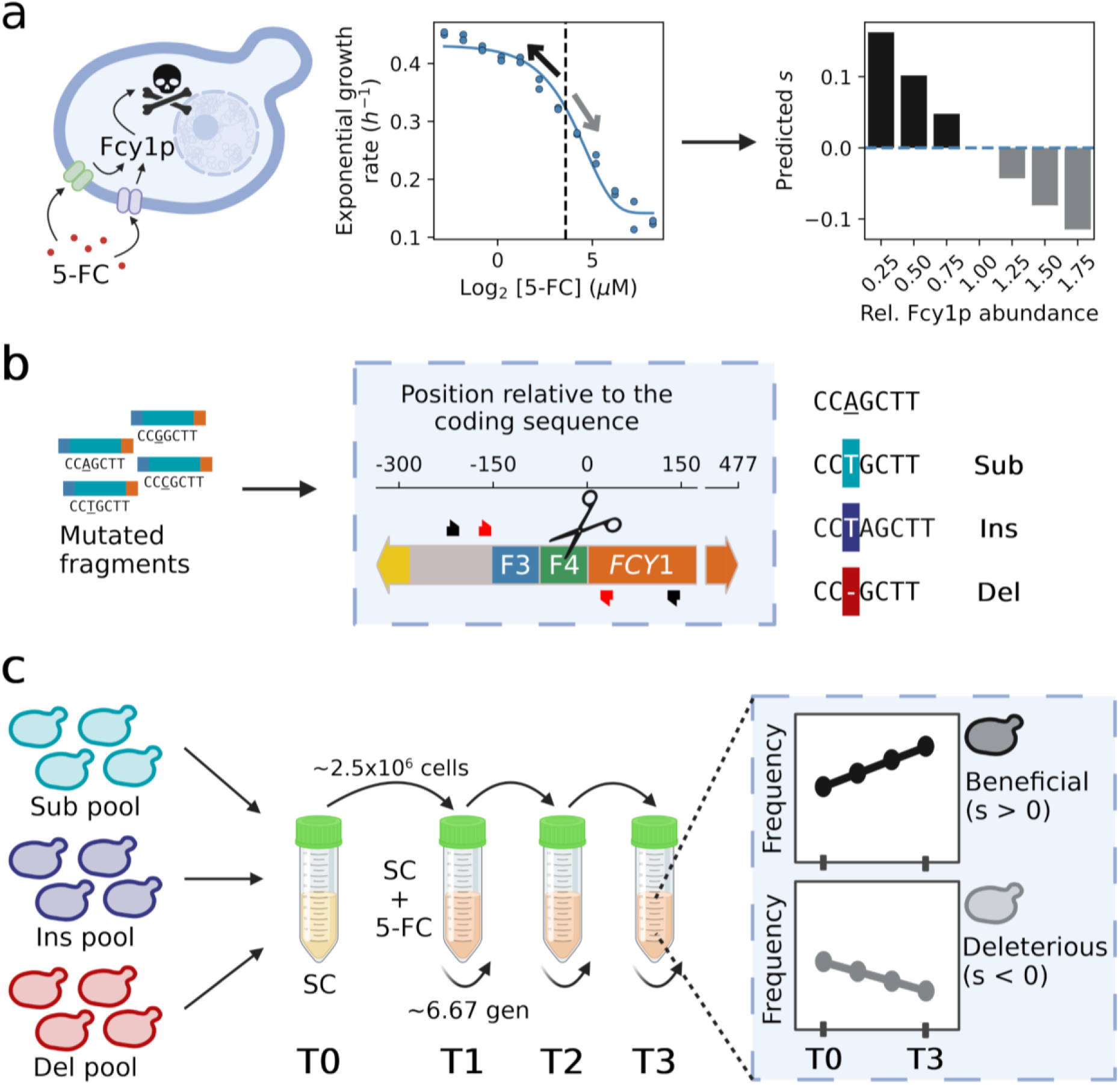
Assaying the accessibility of adaptation through single mutations in the *FCY1* promoter. **a,** In presence of 5-FC, the activity of cytosine deaminase Fcy1 is toxic for yeast, by catalyzing the conversion of 5-FC into 5-fluorouracil. Accordingly, the growth rate of *S. cerevisiae* decreases with increasing 5-FC concentrations, as shown by the corresponding dose-response curve (**Extended Data Fig. 1a**). Under the naive assumption that changing the abundance of the Fcy1 protein is equivalent to varying the concentration of 5-FC, promoter mutations impacting *FCY1* expression could affect fitness. **b,** To generate mutated *FCY1* promoters, regions F3 and F4 (154 bp closest to the coding sequence) were replaced with libraries of synthetic sequences corresponding to all possible single-nucleotide substitutions (Sub), insertions (Ins) and deletions (Del). This was performed at the native genomic locus, using a CRISPR-Cas9-based method (*Methods*^39,40^). In downstream experiments, the various single mutants were genotyped by sequencing of the F3-F4 region, using oligonucleotide pairs shown in red (short fragment) and in black (extended fragment). **c,** The three types of mutant pools (Sub, Ins, Del) obtained from the genomic insertion of F3 and F4 variants were combined and grown in synthetic complete media (SC) and then transferred to media with 5-FC to allow for competition based on the abundance of Fcy1p. Loss of function of *FCY1* has no detectable effect in SC media without 5-FC. At every timepoint, DNA was extracted and the edited locus was sequenced to follow the frequency of each mutant in the culture, allowing to estimate selection coefficient for each mutation from their enrichment or depletion through time. Schematics created in BioRender. Aubé, S. (2026) https://BioRender.com/k3mujqk.

The mutagenized region corresponds to two fragments hereafter named F3 and F4 (**Fig. 2b**). Within each of them, we generated all single-nucleotide mutations – substitutions, insertions and deletions. The corresponding variants of F3 and F4 were inserted at the endogenous *FCY1* locus (YPR062W) using a CRISPR-Cas9-based pooled approach (**Fig. 2b**, *Methods*^39,40^). This resulted in 1021 unique mutated sequences (462 substitutions, 463 insertions and 96 deletions), of which 1020 were successfully recovered by sequencing of the mutant pools (**Supplementary Fig. 2**). This process was repeated four times for each of the three classes of mutations. The corresponding pools of substitutions, insertions and deletions were then combined (**Fig. 2c**), resulting in four independent transformation replicates, each containing all mutations. Because the set of substitutions was obtained by successively degenerating each nucleotide position, the WT *FCY1* promoter was also (re)constructed along with the desired mutants, at a frequency of ∼17%.

From each of the four transformation replicates, two culture replicates were made, which were both grown in liquid media with [12 μM] 5-FC and without, resulting in 16 cultures. This concentration inhibits the growth of *S. cerevisiae* by ∼50% and was previously used to assay all single-codon substitutions in the *FCY1* coding sequence^36^ (**Fig 2a**, **Extended Data Fig. 1**). Additional replicates of all 16 cultures were also done by combining the promoter mutants with 960 representative coding sequence variants^36^, to act as positive controls (**Extended Data Fig. 3**).

All 32 cultures were grown for ∼20 generations in liquid media, with dilution every ∼7 generations to maintain exponential growth (*Methods*). Samples were taken at each dilution step, as well as before the start of the experiment, resulting in a series of timepoints from T0 to T3 (**Fig. 2c**). The relative frequencies of all mutations were monitored by sequencing of the F3-F4 region (**Fig. 2b**) and the selection coefficient (𝑠) associated with each promoter sequence was estimated from the slope of its log-scaled frequencies relative to the WT^41^ (**Fig. 2c**).

Mutant pools sampled from the competition experiment were first genotyped by sequencing a 174 bp fragment, which covered the full F3-F4 region and the first 20 bp of *FCY1* (red oligonucleotide pair; **Fig. 2b**). There were however major reproducibility issues. While estimated selection coefficients were consistent between culture replicates of the same transformation, they diverged substantially across transformation replicates (**Supplementary Fig. 1a**). In addition, comparisons of the distributions of selection coefficients across the different time intervals revealed a gradual shift of the mean. From ∼0 for the T0-T1 interval, it decreased below −0.15 for the full competition experiment (**Supplementary Fig. 1b**). Through further experiments, we found that these effects were caused by secondary mutations introduced during library construction (**Supplementary Fig. 1c-d**). We modified our sequencing protocol to genotype a longer region (black oligonucleotide pair; **Fig. 2b**) and repeated the analyses to eliminate these effects. We explain the observations in detail in the supplementary material (**Supplementary Note**) as we suspect these could be common in genome editing studies and thus be of broad interest.

### Fitness is robust to *FCY1* promoter mutations

After correcting for the impact of secondary mutations, the inferred selection coefficients for the F3-F4 mutants reveal only small fitness effects, which vary mostly uniformly along the mutagenized region (**Fig. 3a**). Across all assayed promoter mutations, median *s* estimates obtained from replicate experiments range from ∼-0.037 to ∼0.035 𝑔𝑒𝑛^−1^, for a grand median of ∼-0.002 𝑔𝑒𝑛^−1^. Experimental noise moreover appears of a similar magnitude as these deviations from 0, as shown by an average difference of ∼0.028 𝑔𝑒𝑛^−1^ between the highest and lowest *s* among replicate estimates for the same genotype, up to ∼0.110 𝑔𝑒𝑛^−1^ (**Extended Data Fig. 4a**). A small number of secondary mutations remain undetected (not found within the longer sequencing fragment), resulting in outlier selection coefficients for one pair of culture replicates within some F3-F4 genotypes. Following stringent removal of these rare outliers (**Supplementary Fig. 8**), none of the 1020 *s* estimates for *FCY1* promoter mutations appear significantly different from 0 (𝑝 > 0.05; one-sample t-test with Benjamini-Hochberg correction at FDR=5%). Identifying significant effects using |𝑠| thresholds on the most extreme replicates instead results in small numbers of significant changes (12 at |𝑠| > 0.01), which are consistent with the null expectation from random permutations of the data (**Extended Data Fig. 4**). Overall, we thus conclude that there were no measurable fitness effects among the assayed single-nucleotide mutations.

**Fig. 3:**
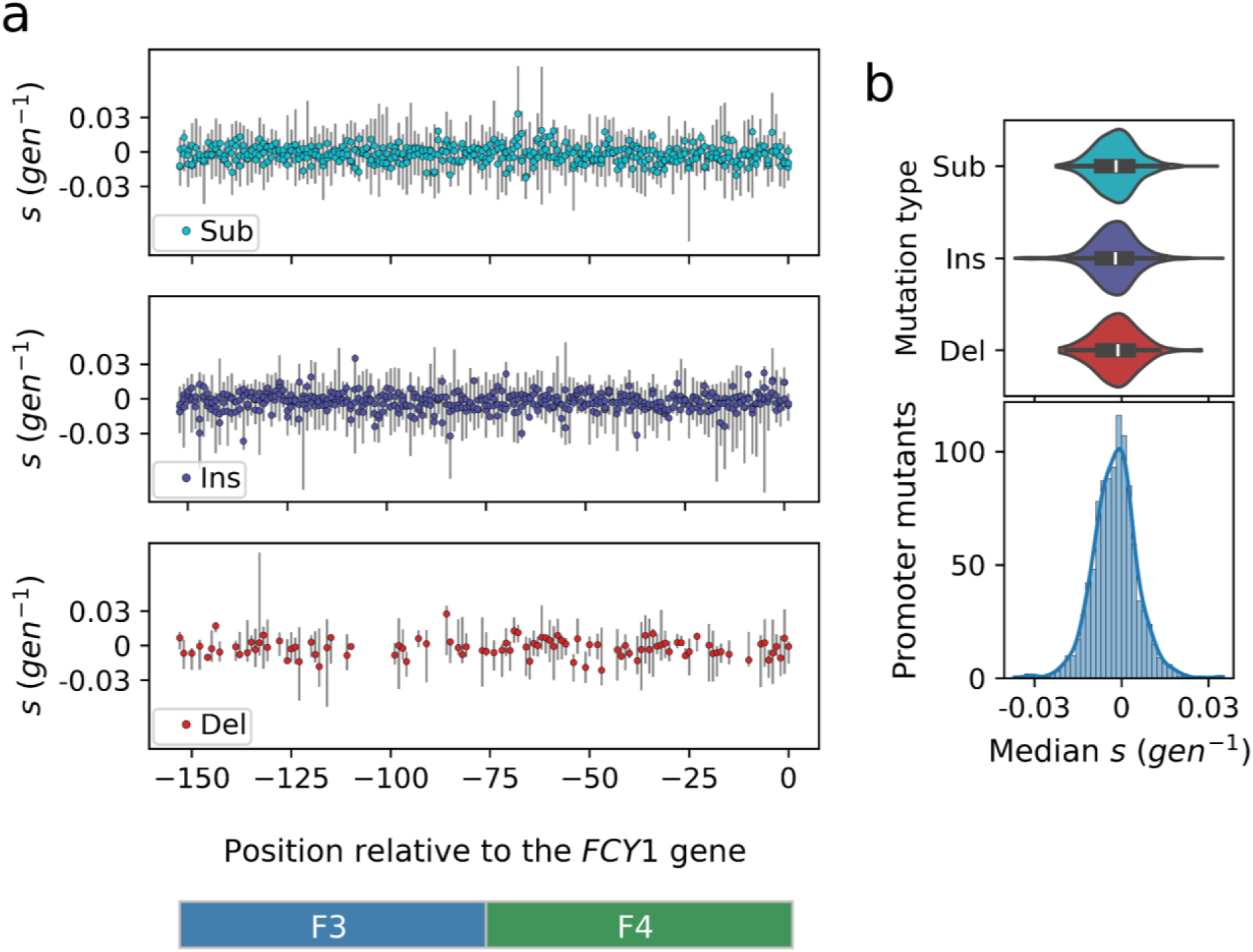
Mutations in the *FCY1* promoter do not have a measurable impact on fitness in 5-FC conditions. **a,** Fitness effects of single-nucleotide mutations in the *FCY1* promoter. All substitutions (Sub; n=462), insertions (Ins; n=462) and deletions (Del; n=96) are shown. Points display the median selection coefficient for each mutant genotype, while error bars show the minimal and maximal values observed at each position along the mutated region. Nucleotide positions which appear to be missing are due to mutations which result in the same genotype (**Supplementary** Fig. 2), for instance deletion of any single base in a homopolymer region. Only data from two unbiased transformation replicates (each duplicated into two culture replicates; four samples in total) are shown (**Supplementary Note**, **Supplementary** Fig. 6). **b,** Distribution of mutational effects on fitness for single-nucleotide mutations in the *FCY1* promoter. The median selection coefficient obtained for each single mutant is shown, either as separate distributions for each of three classes of mutation (top) or as the full distribution (bottom; n=1020 observations, one per mutant).

This absence of mutational effects is not due to a failure of the competition experiment, as confirmed by two layers of controls. First, when considering all mutations (F3-F4 and secondary), the variance of estimated selection coefficients is indeed much narrower for cultures without 5-FC (**Supplementary Fig. 9**). Second, selection coefficients inferred for the added coding sequence mutants correlate well with previously published variant effects^36^ (𝜌 = 0.780 with 𝑝 < 10^−64^; **Extended Data Fig. 5a**). There is moreover almost perfect agreement between the two experiments regarding the number and identity of resistance-conferring mutations (**Extended Data Fig. 5b**). Together, these results confirm that adaptive changes would have been observed in our assay, if they could occur through single-nucleotide promoter mutations. We note that, whereas samples which contained coding sequence variants also included the complete pool of promoter mutations (**Extended Data Fig. 3**), the resulting cultures were not used to precisely estimate F3-F4 mutational effects. Although noisier, the corresponding selection coefficients were nonetheless qualitatively similar to those reported in **Fig. 3a** (**Supplementary Fig. 10**).

This absence of adaptive mutations within the *FCY1* promoter contrasts markedly with the high accessibility of adaptation through changes in the corresponding coding sequence. Within the *FCY1* gene, ∼29% (408/1411) of single-nucleotide substitutions indeed result in codon changes which confer resistance to 5-FC in the same conditions (**Extended Data Fig. 5c**)^36^. Among these, only 67 introduce a premature stop codon. As such, although adaptation to 5-FC is readily available through *FCY1* mutations, it cannot be conferred by single-nucleotide changes in the promoter. In order to understand how this inaccessibility of adaptation arises, we first measured the effect of promoter mutations on *FCY1* expression.

### Many mutations in the *FCY1* promoter substantially affect expression level

To obtain expression level measurements for all promoter mutants, we used a sort-seq approach^42,43^. The complete pool of F3-F4 single mutants was reconstructed in a genetic background where the distal part of the *FCY1* gene – starting from its 43rd base – was replaced with a mEGFP sequence (*Methods*, **Extended Data Fig. 6**). As such, the translation initiation context could be maintained while adding a fluorescent reporter to measure promoter activity. Yeast cells of the corresponding mutant library were sorted into five bins (numbered 0 to 4) according to their log-scaled fluorescence intensity (**Fig. 4a, Supplementary Fig. 11**). All the resulting samples were sequenced and read counts were used to compute a weighted average of bins for each genotype (*Methods*). The effect of each mutation was quantified using a 𝛥_𝑅𝑒𝑓_ score equal to the difference between the corresponding mean bin and the average obtained for the WT. This was performed separately within each of six replicates (three transformation replicates, each subdivided in two culture replicates).

**Fig. 4:**
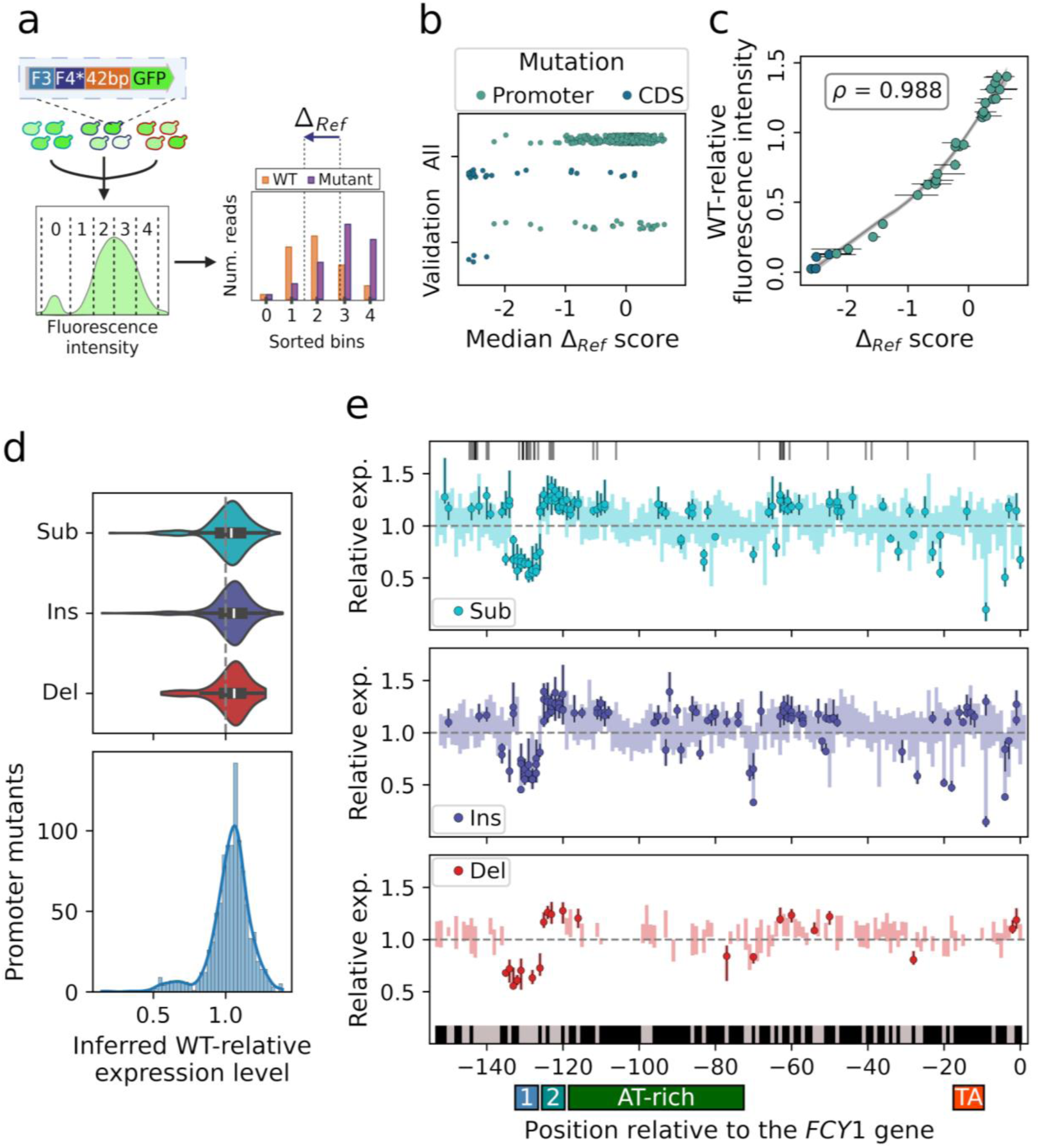
Mutations in the *FYC1* promoter impact expression level. **a,** Schematics of the sort-seq experiment. Yeast cells from pools of promoter mutants were sorted into five bins according to their fluorescence intensity (representative distribution shown). From the sequencing of each bin, WT-relative fluorescence intensity could be estimated as a 𝛥_𝑅𝑒𝑓_ score. Created in BioRender. Aubé,S. (2025) https://BioRender.com/uzzv5xy. **b,** Median sort-seq scores obtained for all variants detected across six replicate experiments. In addition to 1019 F3-F4 variants (“Promoter”), 28 single mutations in the first 42 bp of *FCY1* were identified (“CDS”), of which the vast majority are indels. The full distribution of scores (“All”) as well as the 30 mutants selected for reconstruction (“Validation”) are shown, split between the two categories in each case. **c,** Calibration curve of sort-seq scores. Points display the medians of sort-seq scores (n=6) and WT-relative fluorescence intensities (n=3) for all 30 validation genotypes. Horizontal and vertical error bars show the minimal and maximal observations associated with each median. The high correlation between medians is shown with Spearman’s rho (𝑝 < 10^−23^). The line and the corresponding shaded area display a LOWESS fit with its 95% confidence interval (*Methods*). **d,** Distributions of WT-relative expression levels for all single-nucleotide promoter mutations, as inferred from the calibration of sort-seq scores. At the top, distributions are shown separately for substitutions (Sub; n=461), insertions (Ins; n=462) and deletions (Del; n=96), with a dashed line indicating WT expression level (1.0). At the bottom, the full distribution, combining all three types of single mutants, is shown (n=1019). **e,** Distribution of promoter mutations with a significant effect on expression (n=245), along the mutagenized region. Separately for Sub (n=113), Ins (n=111) and Del (n=21) variants, the estimated WT-relative expression (point) and its corresponding 95% prediction interval (error bar) are shown for each mutant affecting FCY1 expression by 5% or more (minimum or maximum of the corresponding 95% prediction interval; *Methods*). Shaded rectangles display the minima and maxima across all prediction intervals for each position along the sequence, including variants which are not shown. In the upper plot, vertical lines show the midpoint of hits (n=41) to known TF recognition motifs, identified using YeTFaSCo^46^. In the lower plot, the black and white rectangles respectively display A/T and G/C positions. Promoter positions corresponding with the two main clusters (sites “1” and “2”), the approximate AT-rich region and the TATA-like element (“TA”)^47,48^ are also identified.

A narrow range of scores was obtained, with the median 𝛥_𝑅𝑒𝑓_ of most promoter mutations falling between −1 and 0.5 (**Fig. 4b**). Of the 1020 F3-F4 mutants, 1019 were covered in at least two replicates after abundance-based filtering (*Methods*). Many secondary mutations were also recovered (**Fig. 4b**), including indels in the first 42 bp of *FCY1*. Interestingly, while most such mutations were associated with a low fluorescence intensity and thus a negative 𝛥_𝑅𝑒𝑓_, some did not completely abolish expression. We specifically found out that indels which occurred before the second methionine codon of *FCY1* maintained ∼10% of the WT fluorescence level (**Extended Data Fig. 7 a,b**). This is consistent with a previous report of a minor translation initiation site at this codon^44^.

To validate the sort-seq assays, we selected 30 mutants for independent reconstruction (**Fig. 4b**). For each newly constructed strain as well as for the WT reference, fluorescence intensity was measured by flow cytometry (*Methods*). The resulting WT-relative intensities are in strong agreement with the corresponding median 𝛥_𝑅𝑒𝑓_ scores (**Fig. 4c, Supplementary Fig. 12**). We additionally used this validation to draw up a calibration curve, enabling the conversion of sort-seq scores into more easily interpretable relative fluorescence intensities. To this end, we fit a LOWESS regression (**Fig. 4c**), to avoid making specific assumptions about the relationship between the two variables. This was performed within a bootstrapping routine which involved repeated resampling of 𝛥_𝑅𝑒𝑓_ values as well as validation fluorescence measurements, thus generating 95% prediction intervals for the curve itself and for the relative fluorescence intensities of all genotypes (*Methods*, **Supplementary Table 9**). We hereafter refer to the medians of the latter intervals as WT-relative expression levels for the corresponding mutations, where a value of 1 denotes the WT level.

Examining the inferred relative expression levels for all F3-F4 mutants reveals a slight bias towards expression increases, with medians lying between 1.04 and 1.06 (**Fig. 4d**). That mutations are more likely to increase expression than to decrease it may appear surprising, since Fcy1p is already highly expressed and is among the 10% most abundant proteins in *S. cerevisiae*^45^. While this bias could be artefactual, the y-intercept of the calibration curve suggests otherwise (**Fig. 4c**). It is very close to 1 (∼1.00, with 95% prediction interval from 0.98 to 1.02) – as expected for a 𝛥_𝑅𝑒𝑓_ of 0 –, although no WT measurements were used in the LOWESS fitting.

The overall distribution of effects highlights that most single mutations in the *FCY1* promoter impact expression levels by less than 15%, with 75% of variants lying between values of 0.98 and 1.11 (**Fig. 4d**). Yet, we also observe a small number of mutations with larger effects, decreasing expression to ∼50% of its WT level, or even lower. This tail appears more pronounced than previously reported in yeast^3,49^, potentially because we specifically mutated the most proximal part of the *FCY1* promoter, where mutations are more likely to disrupt processes such as transcription initiation. Among the 1019 F3-F4 mutations, 245 significantly affect *FCY1* expression, by 5% or more (95% prediction intervals). In agreement with a previous study of a yeast promoter^13^, these significant mutations are not distributed uniformly along the mutagenized promoter region (**Fig. 4e**). This distribution reveals important determinants of *FCY1* expression level.

The fact that significant mutations form two main clusters, from positions −132 to −127 (site 1) and from −125 to −120 (site 2), first highlights the importance of transcription factor (TF) binding. Within these two sites, 92.4 % (73/79) of all variants change expression by 5% or more, in a consistent direction within each cluster. Both sites are within a 80 bp segment known to be critical for TF binding in yeast promoters^4,42,50^, while the whole −132 to −120 region is a hotspot of predicted TF binding sites^46^ (**Fig. 4e top; Supplementary Table 10**). As such, the clusters likely reveal TF binding sites. Interestingly, the deletion of either of the two sites is not more impactful than the strongest mutations within the corresponding cluster (**Extended Data Fig. 8**), suggesting that single mutations are sufficient to disrupt them.

The position of the two most impactful mutations additionally highlights how alterations of transcription initiation may severely reduce *FCY1* expression. These variants, a substitution and an insertion which respectively decrease expression to ∼16% and ∼13% of its WT level (cytometry; **Fig. 4c**), both occur at position −9 (**Fig. 4e**). This position is immediately downstream of the previously identified TATA-like element of the *FCY1* promoter^47,48^ – from −17 to −10 –, where the transcription pre-initiation complex assembles. Additionally, the transcription of *FCY1* has been shown to initiate at position −7^38,51^. The corresponding mutational effects thus likely reflect the disruption of transcription initiation. The conflicting report of a most abundant transcript isoform beginning further upstream, at −29^52^, however adds uncertainty.

Finally, the complete set of significant mutations underscores that most single-nucleotide changes within the poly(dA:dT) region of the *FCY1* promoter weakly affect its transcriptional activity. The corresponding AT-rich sequence, approximately between positions −118 and −73 (17.39% GC; **Fig. 4e**, bottom), is indeed comparatively depleted in strong mutational effects. Only 17.6 % (52/295) of variants within it are significant, compared to 26.7 % (193/724) in the remainder of the mutagenized region. This may seem contradictory, considering that such regions are important for the establishment of nucleosome-depleted regions within promoters^53,54^, which favor transcription and can be sufficient to induce expression from a downstream core promoter^55^. It is however consistent with previous work on the *URA3* promoter, which showed that deletion of more than half of the corresponding 50 bp poly(dA:dT) was necessary to abolish expression^56^.

Overall, we show that 24% (245/1019) of single-nucleotide mutations in the *FCY1* promoter have a significant effect on expression level, many of which appear to disrupt the binding of TFs at two putative sites. As such, *FCY1* expression is sensitive to mutations, meaning that the DME on expression level is not sufficient to explain the absence of fitness changes in 5-FC conditions.

### The expression-fitness relationship for *FCY1* is mostly flat

To investigate the mechanism underlying the absence of measurable fitness effects, we set out to characterize experimentally the expression-fitness relationship for *FCY1* (**Fig. 5a**). Six yeast promoters covering a wide range of expression levels were used to construct two sets of strains, in which each promoter was inserted upstream of either the native *FCY1* sequence or the previously described 42bp-mEGFP construct (**Extended Data Fig. 6c**). The *FCY1* strains were grown in media with 5-FC (12 μM; like in the competition experiment) – to obtain growth rate measurements –, while mEGFP strains were used to measure expression levels (*Methods*). For each promoter, the measurements from the two strains were combined to draw the corresponding expression-fitness curve (**Fig. 5a**).

**Fig. 5:**
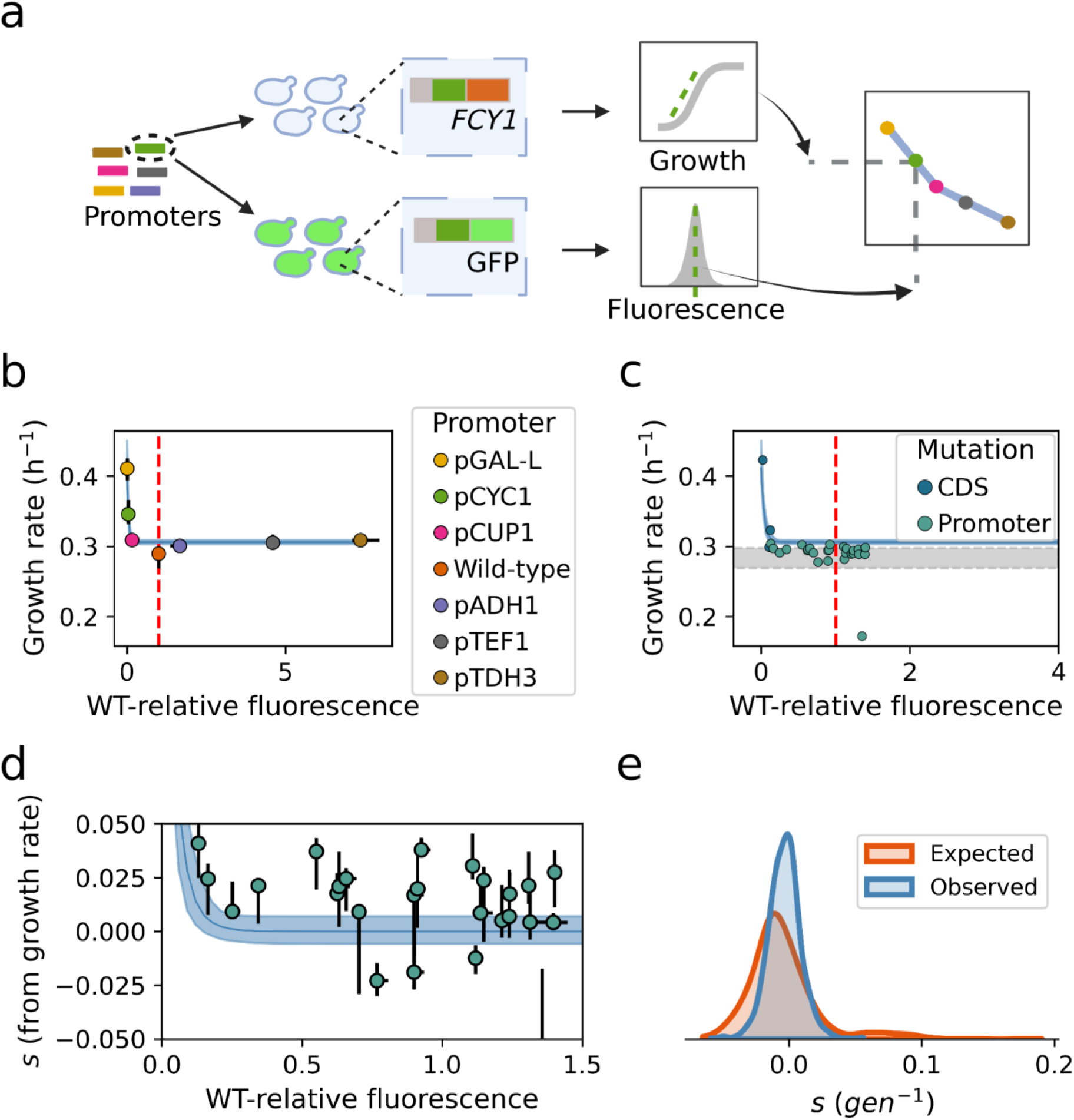
The *FCY1* expression-fitness function is flat around WT expression level, making fitness robust to promoter mutations. **a,** Strains with varying expression levels were constructed to measure the fitness function. A set of yeast promoters covering a wide range of activities were introduced in the genome in place of the endogenous *FCY1* promoter, either upstream of *FCY1* or of a fluorescent reporter. Created in BioRender. Aubé, S. (2025) https://BioRender.com/1p60mp7. **b,** Expression-fitness relationship for *FCY1* in 5-FC conditions. The four promoter constructs expressing *FCY1* to at least ∼18% of its WT level display very similar growth rates. For each promoter, medians of growth rate and WT-relative fluorescence (n=3) are shown along with the corresponding minima and maxima (error bars). The fitness function is displayed as a fitted exponential decay function, with its corresponding 95% confidence interval (*Methods*). The WT strain, which lacks a resistance cassette and is not directly comparable to the other ones (*Methods*), was not used in the fit. **c,** Growth rates of reconstructed mutants are consistent with the fitness function. For validation mutants (Fig. 4c) which could also be reconstructed in the *FCY1* background, medians (n=3) are shown for growth rates in 5-FC [12μM] media and WT-relative fluorescence intensities. This includes single promoter mutations (“Promoter”; n=26) and indels in the first 42 bp of the gene (“CDS”; n=3). The dashed line identifies the WT expression level (1.0), while the shaded area displays the range of growth rates measured for the WT background (panel **a**), into which the mutations have been reconstructed. **d,** Reconstructed promoter mutations are mostly neutral. For the same set of single mutations in the promoter (n=26), medians (n=3) are shown for selection coefficients and relative fluorescence intensities, with error bars displaying the corresponding minimal and maximal replicates. Selection coefficients are computed from growth measurements (*Methods*) shown in the previous panel, relative to the WT strain. The previously fitted fitness function (panels **b** and **c**) is displayed with its corresponding 95% confidence interval, after conversion into selection coefficients. Only the region of the expression-fitness curve where promoter mutants lie is shown. The outlier mutant which is hidden is associated with a median *s* of ∼-0.28. **e,** The shape of the fitness function prevents expression changes from affecting fitness. The DME on fitness observed in the competition experiment is much narrower than the one expected from the relationship between the concentration of 5-FC and growth rate (Fig. 2a, **Extended Data Fig. 1**).

This experiment revealed a very steep fitness function, where growth rate decreases sharply until *FCY1* expression reaches ∼18% of its WT level and then plateaus (**Fig. 5b**). Consequently, this fitness curve is flat around the WT, such that small expression changes have no fitness effect. The shape of this function is thus consistent with the fact that no single-nucleotide promoter mutation conferred resistance to 5-FC in our bulk competition experiment. To further validate this result, we leveraged previously characterized validation mutants. The set of alleles used in the calibration of sort-seq measurements, which also includes a small number of indels in the coding sequence, was reconstructed in the native *FCY1* background and the resulting strains were phenotyped in 5-FC conditions (*Methods*). In combination with the corresponding relative fluorescence intensities, these growth measurements confirm the flatness of the curve around WT expression level. Almost all promoter mutants display WT-like growth rates, with no clear trend along the complete range of effects (**Fig. 5c**). Among three assayed coding sequence mutations, the only one which reached the high fitness region is associated with a ∼98% decrease in fluorescence intensity (**Supplementary Table 11**), caused by a frameshift-inducing indel. This highlights how adaptation is only possible through very large reductions in Fcy1 abundance, which cannot be achieved by single promoter mutations. Zooming in on the promoter mutants further illustrates this point. Among them, the two single mutations with the largest effect barely reach the threshold below which fitness rapidly increases, although they respectively decrease *FCY1* expression to ∼16% and ∼13% of its WT level (**Fig. 5d**). As such, the shape of the fitness function, jointly with the corresponding DME, renders adaptation to 5-FC inaccessible from single changes in the *FCY1* promoter.

The flatness of the fitness function around WT expression contrasts markedly with the steepness of the 5-FC dose-response curve around the 12 μM concentration used (**Fig 2a**, **Extended Data Fig. 1**). If varying the abundance of Fcy1p instead had the same effect as changing the 5-FC concentration, large fitness differences would have been observed among the set of promoter single mutants. The most impactful mutation, decreasing Fcy1p abundance to ∼13% of its WT level, would have been associated with a *s* of 0.19, while seven other mutations would have resulted in 𝑠 > 0.1 (**Fig. 5e**). Adaptation would then have been accessible through regulatory changes, albeit much less readily than from mutations in the coding sequence.

Whereas growth rate varies according to 5-FC concentration, the shape of the fitness function remains mostly constant. Across concentrations resulting in ∼10% to ∼70% inhibition for the WT, growth rate variation is restricted to the same narrow range of relative expression levels and only the height of the corresponding plateau varies (**Extended Data Fig. 9a**). A similar phenomenon is observed in media lacking uracil, where the deamination of cytosine into uracil by Fcy1p is essential for growth. With increasing concentrations of cytosine, the height of the plateau of high fitness increases, while the region of varying fitness remains mostly the same (**Extended Data Fig. 9b**). At 5-FC and cytosine concentrations resulting in similar levels of relative inhibition or growth, the fitness functions obtained in the two conditions are very similar, with a sign inversion (**Extended Data Fig. 10ab**). This suggests that our results also extend to the canonical cytosine deaminase function of the Fcy1 enzyme. Growth measurements for a set of promoter mutants covering the full range of mutational effects confirm that increasing *FCY1* expression, as allowed by the DME of single mutations, is not adaptive under uracil starvation (**Extended Data Fig. 10c**). Overall, these experiments demonstrate that adaptation is not accessible from single-nucleotide changes in the *FCY1* promoter, in LOF (5-FC) as well as in gain-of-function (cytosine) contexts. This is due in part to the small number of large mutational effects on gene expression, but more importantly to the shape of the fitness function, which is flat around WT expression level.

## Discussion

The rapid accessibility of adaptive changes through regulatory mutations depends on their effect sizes on gene expression and on the relationship between fitness and expression level. One powerful approach to investigate these determinants is to focus on a gene where adaptation is known to arise and systematically assay all single regulatory mutations. To this end, we studied the promoter of the gene *FCY1*, a protein-coding region within which single-nucleotide changes have been shown to confer resistance to the drug 5-FC^36,57^. We show that no promoter mutation has a measurable fitness effect in yeast exposed to this molecule, although large changes in expression level (reaching a reduction of 87%) are observed. In this adaptive landscape, beneficial phenotypes are thus inaccessible from single nucleotide variation in the promoter, although they arise readily from mutations in the corresponding coding sequence^19,36^.

Our initial data instead suggested that most promoter mutations were highly deleterious. Further investigations revealed that this result was due to secondary resistance-conferring mutations introduced during library construction, outside of the sequenced region (**Supplementary Note; Supplementary Fig. 1,4-8**). These beneficial mutations largely occurred in the WT promoter background, since this genotype was generated at high frequency within the mutant pool. This comparatively made all true promoter single mutants appear less fit. Interestingly, bias thus arose from the reconstruction of the WT within the CRISPR library, which is widely regarded as a best practice to ensure the accuracy of mutant fitness estimates^58^. Such artifacts may be common in large-scale genome editing experiments, due to the increased frequency of single-nucleotide mutations – and especially indels – in synthetic DNA oligonucleotides^59^. This highlights the importance of designing library construction workflows which minimize the use of genic regions as homology arms to guide recombination. Alternatively, these segments should at least be fully included in the genotyping region, as we did to correct our experiment.

Within our *FCY1* model system, the inaccessibility of adaptation through promoter mutations emerges jointly from the shape of the fitness function and from the DME on expression level. Whereas the fitness function is flat around WT protein abundance, single promoter mutations do not push expression level far enough to escape this fitness plateau. A related phenomenon has been observed for the *TDH3* gene of *S. cerevisiae*, for which the effects of cis-regulatory mutations on expression and fitness have been extensively studied^10–12,49,60^. A vast majority (216/235) of single mutations in the *TDH3* promoter were reported to lie on a plateau of the fitness function and thus have no significant effect on fitness^10^. Yet, because the deletion of *TDH3* only decreases fitness by ∼5%^10,12^, the relevance to adaptation is unclear. For *FCY1* in 5-FC conditions, a 30-40% fitness increase is in contrast possible through LOF (Extended Data Figs 2, 7). Similarly, the characterization of 1400 single-nucleotide substitutions in the *SUL1* promoter of *S. cerevisiae* revealed only 61 mutations significantly affecting fitness, with the largest reported increase approaching 10%^13^. In this case, cis-regulatory mutations were assayed in conditions where *SUL1* coding mutations and copy number amplifications can respectively increase fitness by 23%^61^ and 51%^33^. While this is more directly comparable to *FCY1* in 5-FC, the corresponding DME on expression level was not measured and the fitness function was only partially characterized^14^. As such, it is difficult to assess whether the paucity of fitness effects among mutants of the *FCY1*, *TDH3* and *SUL1* promoters emerges from similar mechanisms. More studies characterizing additional promoters as well as the corresponding fitness functions are needed.

Although data on multiple cis-regulatory regions is currently lacking, dissecting how the absence of adaptive mutations in the *FCY1* promoter arises still offers clues regarding the generalisability of this observation. An important caveat is however that this model may not represent a typical gene. Because the same loss of expression which allows adaptation to 5-FC also impedes the canonical cytosine deaminase activity (**Extended Data Figs 9, 10**), the inaccessibility of LOF-based adaptation through promoter mutations may reflect past selection to improve this latter function. This is suggested by the high abundance of Fcy1p, in the top 10% among yeast proteins^62^, which could have arisen by progressively climbing up the increasing fitness curve observed in cytosine conditions (**Extended Data Fig. 9**). Such an evolutionary trajectory would imply that the apparent plateau of the fitness function actually displays a small curvature, visible to natural selection but not measurable using our experimental approach^63^. As such, the *FCY1* model may be representative only of adaptive pathways which involve the loss of a previously optimized function. It nonetheless provides a useful framework to understand how and under which conditions adaptation may occur through a single cis-regulatory mutation. An additional caveat is that we have only assayed the adaptive potential of *FCY1* mutants under constant 5-FC exposure. It is plausible that, in a fluctuating environment where the concentrations of 5-FC and cytosine would both vary, adaptation could occur through cis-regulatory mutations.

In the current study, the absence of adaptive mutations in the *FCY1* promoter first arises from the rarity of mutations strongly decreasing expression. This is largely consistent with previous work, and thus likely to apply to a number of other genes. The large-scale mutagenesis of promoters and regulatory elements in yeast^3,49^ and mammals^6–9^ indeed showed that only a minority of single mutations strongly affect expression. In the yeast *TDH3* promoter, for instance, only 7.7% (18/235) of mutations were reported to decrease expression by more than 6%^49^, compared with 17.2% (175/1019) in the *FCY1* promoter (**Fig. 4d**). Moreover, genome-scale computational predictions indicate that 10% of cis-regulatory mutations account for 50% of expression variation in yeast^4^. The DME on expression level we report is thus likely representative of many yeast genes, although larger changes could potentially have been observed if indels of variable lengths^64^ had been included. The *FCY1* promoter is in addition a typical yeast promoter. It is TATA-less like the majority of such regions^47,65^, features the most common poly-T/poly-A configuration^54^ and belongs to the most abundant class of yeast promoters (2474/5378)^66^. Whereas it is bidirectional, it drives higher expression of *FCY1* than of the opposite gene and there are hundreds such promoters in the yeast genome^67^.

Second, the saturating shape of the *FCY1* fitness function prevents even the most impactful promoter mutations from affecting fitness. Such abundance-fitness curves featuring a plateau have generally been reported for enzymes across organisms^1,68–72^, including the well-studied *TDH3* of *S. cerevisiae*^10,12^. They are additionally expected from enzyme kinetics and metabolic control theory^63,69^. In the case of *FCY1*, the flatness of the fitness function around WT expression was unexpected from the steep growth response to changes in 5-FC and cytosine concentrations (**Extended Data Fig. 1**). This suggests that permeases, rather than the Fcy1 enzyme, may be exerting control over metabolic flux. A similar phenomenon has been reported in bacteria, with the β-galactosidase enzyme and the corresponding permease. While the former displays a step-like fitness function as *FCY1*, the latter is associated with a much more graded fitness function, such that small activity or abundance changes may have selective consequences^70^. More generally, recent proteomics and metabolomics studies have shown that, in most cases, changing the abundance of a single yeast enzyme has a weak impact on the corresponding metabolic flux^73–75^. Overall, it thus appears likely that the shape of the *FCY1* fitness function emerges from fundamental principles. As such, it is plausible that LOF-based adaptation is similarly inaccessible from single mutations in the cis-regulatory regions of a substantial number of other enzymes, in yeast but potentially also in other organisms.

Contrary to promoter mutations, single-nucleotide substitutions in the *FCY1* coding sequence readily confer adaptation to 5-FC^36^ (**Extended Data Fig. 5**). In accordance with the fitness function we report (**Fig. 5b**), this would mean that 29% (408/1411) of coding mutations decrease the abundance or activity of the Fcy1 enzyme below 5% of its WT level. This mutational accessibility is consistent with numerous other studies showing that a substantial fraction of missense mutations in protein sequences induce LOF^68,76–83^. The contrast with single mutations in the *FCY1* promoter, among which the most impactful decreases expression to 13% of its WT level, is nonetheless striking. If large-effect mutations are generally rare in cis-regulatory regions – as is the case for *FCY1*, *TDH3*^10,12,49^ and *SUL1*^13^ in yeast –, LOF-based adaptation would then largely occur through coding changes. Whether coding mutations contribute predominantly to adaptation, including in gain-of-function contexts, remains a fully open question. Two recent surveys of adaptive mutations, in yeast evolved across different environments^28^ and in pathogenic fungi exposed to diverse antifungal drugs^19^, have collectively identified only tens of adaptive noncoding changes, compared with thousands of coding mutations. Yet, a large fraction of these mutations are LOF-inducing. Single promoter mutations conferring adaptation have moreover been identified across bacteria^84–86^, yeast^87^ and humans^15,16^ and have even been associated with a reduction of expression in some instances^15,16,85,87^. Fitness differences between two yeast strains have in addition previously been attributed largely to noncoding variants^88^. As such, elucidating whether coding or regulatory mutations generally contribute more to adaptation will likely require additional experiments as well as a systematic literature comparison similar to that of Hoekstra & Coyne^89^.

Recent work demonstrates that it is now possible to comprehensively assay the effect of mutations in full-length promoters, even in multicellular organisms^91^. An important complement to such work should be the characterization of the corresponding fitness functions. In *FCY1*, its step-like nature makes promoter mutations much less impactful on fitness than expected (**Fig. 5e**). More generally, an important role for fitness function shape in defining whether cis-regulatory or coding mutations contribute more to adaptation was previously theorized from data on a handful of mutations conferring β-lactam resistance in bacteria^90^. When the WT resides on a fitness plateau, as is the case for *FCY1*, expression changes are unlikely to provide a large fitness benefit (**Fig. 6a**). Coding mutations, on the other hand, may cause complete LOF – like in our *FCY1* model – or bypass the fitness plateau by modifying the curvature of the fitness function (in gain-of-function contexts), for instance by changing the affinity of an enzyme for its substrate^90^. When the WT instead resides in a region of the curve where fitness varies rapidly, expression variation provided by regulatory mutations could be equally beneficial as coding changes (**Fig. 6b**). An additional possibility, suggested by the discrepancy between cis-regulatory and coding effects on Fcy1p abundance, would be that, in case of a peaked fitness function, regulatory mutations could be favored due to large-effect coding changes (e.g. LOF) overshooting the optimum (**Fig. 6c**).

**Fig. 6:**
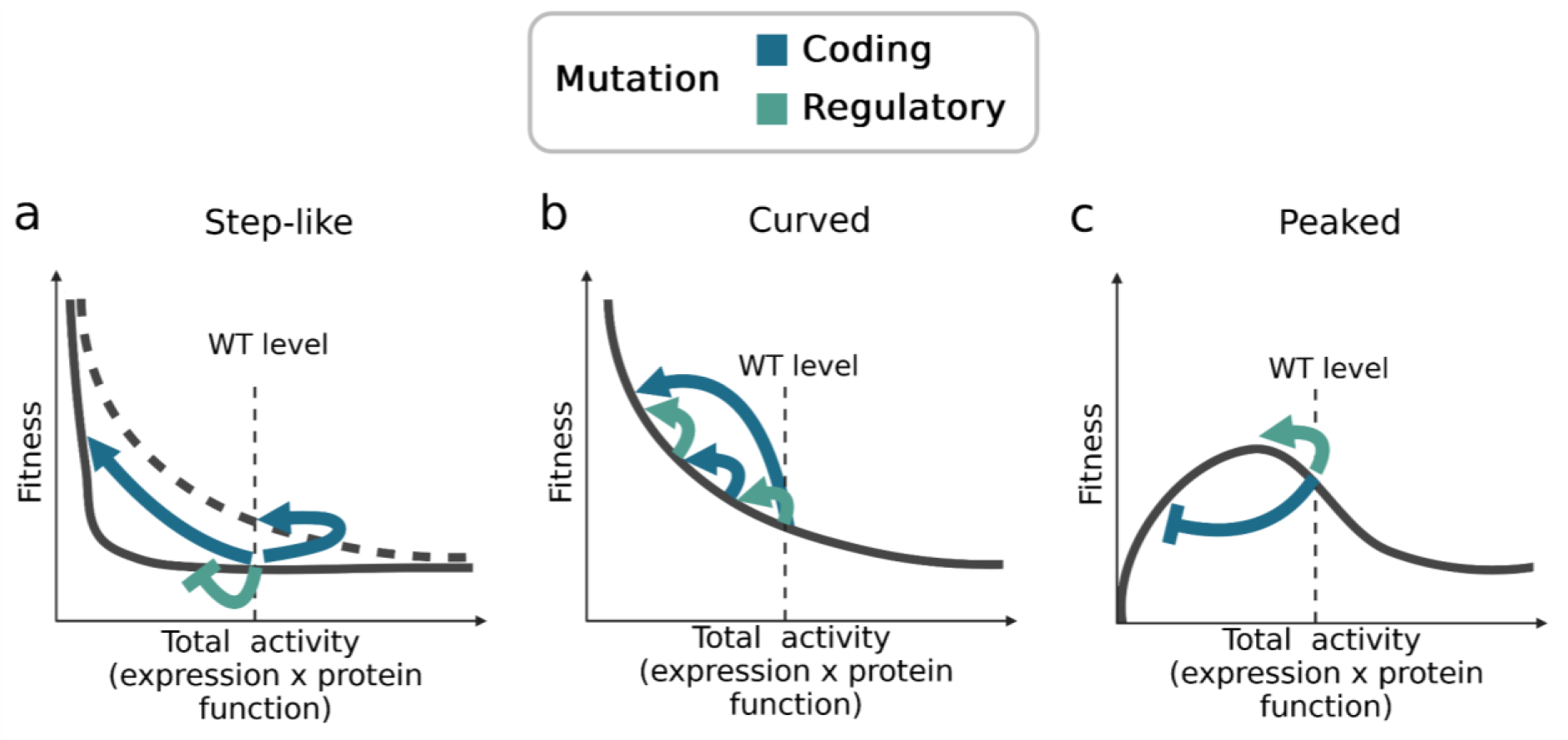
Different shapes of fitness function may favor rapid adaptation through cis-regulatory or coding changes. Although we only measured expression level for *FCY1*, fitness functions are here generalized to total protein activity. For consistency with our experiments, all three are drawn under the assumption that a decrease in activity provides a benefit. **a,** A step-like fitness function where the WT lies on a plateau, far from the fitness cliff, would favor mutations inducing complete LOF or modifying the fitness curve through functional changes in the protein^90^. If LOF cannot occur through a single cis-regulatory mutation, as we report for *FCY1*, this type of mutation will not be selected for (blunt arrow). **b,** A fitness function with high curvature around WT would allow a wide range of mutational effects to provide a benefit. LOF mutations, as well as cis-regulatory or coding changes with more moderate effects on activity, may all be selected for. **c,** A non-monotonic fitness function would favor smaller changes in total activity. Because they overshoot the curve’s optimum, LOF-inducing mutations would be deleterious and accordingly selected against (blunt arrow). If such large effects were only found among coding mutations, cis-regulatory changes could be comparatively favored. Created in BioRender. Aubé, S. (2026) https://BioRender.com/ksrv5jw.

Altogether, previous studies and our results demonstrate the importance of considering the shape and curvature of fitness functions to understand the molecular bases of adaptation. Jointly with the DME on expression level and other molecular traits, fitness functions dictate the availability of adaptive changes. Through widespread characterization of mutational effects and expression-fitness relationships, the breadth of adaptive pathways available in different contexts may be better understood, revealing whether major trends exist among genes or every gene is idiosyncratic. This importance of shape and curvature goes beyond the relationship between fitness and expression levels in microbes, as it appears key for phenotypic traits in general. For instance, it was recently shown that considering the full shape of gene dosage response curves significantly enhances our understanding of genotype-phenotype relationships in humans^92^.

## Methods

### Strains, oligonucleotides, plasmids and culture media

All *S. cerevisiae* strains constructed in this study were generated from a BY4742 (*his3Δ leu2Δ lys2Δ ura3Δ*) background^93^ harboring a hygromycin resistance marker at the *HO* locus (*hoΔ::HPHMX*). This AKD1123 parental background and all other strains are listed in **Supplementary Table 1**, along with the identity of the corresponding genomic inserts and PCR primers as well as the selective media used. The sequences and references for all oligonucleotides used in this study can be found in **Supplementary Table 2**, while references for all plasmids are listed in **Supplementary Table 3**. Recipes for culture media are detailed in **Supplementary Table 4**. When specified, selective media was prepared by adding one or multiple of the following agents: geneticin [200 μg/mL] (G418; Bioshop Canada), nourseothricin [100 μg/mL] (NAT; Jena Bioscience) and hygromycin B [250 μg/mL] (HYG; Bioshop Canada). All Sanger sequencing was performed at the sequencing platform of Centre de recherche du CHU de Québec-Université Laval (Université Laval, Québec, Canada).

### Yeast transformation

Whenever inserts which included a selection marker had to be inserted in the yeast genome, a standard lithium acetate yeast transformation protocol was used^94^. After selection of transformants on the appropriate selective media, four to eight clones were selected for validation PCR, depending on the strain. Unless stated otherwise, colony PCR was performed on each side of the integration site to confirm proper integration, using Taq polymerase (Bioshop Canada). All genomic inserts were amplified using the KAPA HiFi HotStart DNA polymerase (Roche Molecular Systems).

This approach was for instance used to delete each fragment F1 to F4 of the intergenic region between *JID1* (*YPR061C*) and *FCY1* (*YPR062W*), as well as the full intergenic segment and the complete *FCY1* region (including the corresponding gene), by the insertion of a *natNT2* cassette^95^. This resulted in a set of six strains harboring a different deletion of the *FCY1* region (SA0127, SA0128, SA0129, SA0130, SA0131, SA0132). Similar deletions of regions F3 and F4 were also performed in the 42bp-mEGFP construct obtained by CRISPR-Cas9 yeast transformation (see below), generating fluorescent strains SA0246, SA0247 and SA0254

### CRISPR-Cas9 yeast transformation

When sequences needed to be inserted in the yeast genome without any flanking selection marker, a CRISPR-Cas9 approach was used. This required the prior generation of a strain harboring a *natNT2* cassette at the chosen insertion site, which was done through standard yeast transformation as described above. Competent cells from this genetic background could then be cotransformed with the pCAS-NAT plasmid (**Supplementary Table 3**) – encoding a guide RNA engineered to target the Cas9 endonuclease to the *natNT2* cassette^39^ – and the desired repair DNA, amplified with KAPA HiFi HotStart DNA polymerase (Roche Molecular Systems). A previously published CRISPR-Cas9 transformation protocol was followed^96^. Briefly, competent cells were prepared as described, and transformed with unpurified PCR amplification of the insert, DNA carrier [10 mg/ml], pCAS-NAT and PLATE solution (polyethylene glycol 2000 [40%], lithium acetate [0.1 M], Tris-EDTA [10 mM Tris-Cl, 1 mM EDTA] buffer [pH=8.0]). After mixing, cells were incubated for 30 minutes at 30°C, heat shocked for 15 minutes at 42°C, resuspended in YPD media and incubated for 4-6 hours at 30°C. Transformation reactions were plated on YPD media containing G418, to select for uptake of the pCas plasmid, and incubated at 30°C for 48 hours. Eight colonies from each transformation were then grown overnight (30°C with shaking) in liquid YPD and streaked on solid YPD. After 48 hours of incubation at 30°C, one isolated clone from each streaking was grown overnight in YPD (30°C with shaking) and spotted on both YPD and YPD+G418 solid media, to confirm the loss of the pCAS plasmid. Because the AKD1123 parental background is resistant to hygromycin, HYG was added to the YPD throughout. Correct integration was then validated by colony PCR and Sanger sequencing.

Using this protocol, the reinsertion of deleted regions F2, F4 and F1-F4 (full intergenic) was performed, resulting in strains SA0162, SA0166 and SA0170. It was also used to generate the 42bp-mEGFP construct (SA0228) to measure *FCY1* expression level and all validation clones expressing either *FCY1* or mEGFP from the *FCY1* locus.

### Growth assays

Unless stated otherwise, the following protocol was used each time the growth of *FCY1* strains was assessed in media containing 5-FC (Alfa Aesar) or cytosine (Alfa Aesar). Each yeast strain of interest was first inoculated in three replicates in SC [pH=6] liquid media – potentially with HYG, NAT and/or G418, depending on each strain’s resistance marker(s) – and grown overnight at 30°C with shaking. After ∼16 hours of incubation, a sample (100 μl) of each culture was centrifuged and resuspended in the same volume of sterile water, before being aliquoted and diluted again in sterile water to obtain up to 250 μl of cells at 1 optical density (OD) unit/ml. Growth assays were then assembled in Greiner flat-bottom transparent 96-well plates as follows: 160 μl of media (SC complete for 5-FC or SC-Ura for cytosine); 20 μl of diluted cells [OD=1]; 20 μl of 5-FC or cytosine diluted to obtain the desired final concentration in a volume of 200 μl. Each plate was incubated for 48 hours at 30°C without shaking in a Tecan Infinite M Nano (Tecan Life Sciences) and 𝑂𝐷_595_ was measured at 15 minutes intervals.

To obtain exponential growth rates, the exponential part of each growth curve was first identified. The end 𝑇_𝑚𝑎𝑥_ of the exponential phase was defined as the time where the derivative of OD, estimated as the moving median of differences between consecutive OD measurements, reached its maximum. To limit experimental noise, the first two hours of OD measurements were discarded, while any 𝑇_𝑚𝑎𝑥_ above 30 hours or below 3 hours was ignored. In such cases, the first ten hours of the growth curve were instead defined as the exponential phase. Exponential growth rates were then estimated as the slope of the linear regression of 𝑙𝑜𝑔(𝑂𝐷) over time. When converting the growth rate of any strain 𝑖 into selection coefficient 𝑠_𝑖_, the following formula was used^97^:

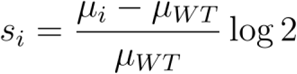

Variables 𝜇_𝑖_ and 𝜇_𝑊𝑇_ are the growth rates for the strain of interest and the corresponding WT reference, respectively. This comparison was always made using WT measurements obtained in the same experiment as the growth rate of strain 𝑖.

### Construction of the mutant libraries

#### Amplification of mutated F3 and F4 promoter fragments

All single-nucleotide substitutions, insertions and deletions in the F3 and F4 regions were ordered as six separate oPools Oligo Pools (Integrated DNA Technologies Inc.) containing sequences of 117 bp, 118 bp and 116 bp, respectively. Within the different pools, each oligonucleotide is a variant of the full-length F3 or F4 region (77 bp) with flanking 20 bp homology regions at both ends. The latter sequences are respectively the 20 bp immediately upstream or downstream of the corresponding fragment in the reference genome of *S. cerevisiae* S288C, as retrieved from the Saccharomyces Genome Database^98^. For substitutions, each position of each fragment was iteratively degenerated into all four possible nucleotides by specifying a N base, while for insertions, the same mixed base was inserted in 3’ of each position in the reference F3 and F4 sequences. When such a N is encountered, an equal ratio of each of the four standard nucleotides (A, T, G, C) is dispensed during synthesis. For the two pools of deletions, single-nucleotide removal was performed iteratively at each position of F3 and F4 and duplicates (occurring whenever two or more consecutive nucleotides are identical in the reference) were removed to obtain non-redundant sets. All sequences ordered as part of the six oPools are listed in **Supplementary Table 5**.

Upon reception, each oPool was spun down and resuspended in fresh PCR-grade water, to obtain a final concentration of 50 μM. Aliquots of 100 μl at 0.05 μM were then obtained by serial dilution from 2 μl of the initial resuspension. From these aliquots, PCR amplification of each oPool was performed in four replicates using KAPA HiFi HotStart DNA polymerase (Roche Molecular Systems) and the following cycle: 98°C for 3 min; {98°C for 30 s; 60°C for 15 s; 72°C for 10 s} x 22; 72°C for 20 s; Hold at 10°C. Each reaction was done in 50 μl:

| Volume (µl) | Reagent |
| --- | --- |
| 33 | PCR-grade water |
| 10 | KAPA HF buffer [5X] |
| 1.5 | dNTPs [10 mM] |
| 1.5 | Forward oligo [10 µM] |
| 1.5 | Reverse oligo [10 µM] |
| 1.0 | KAPA Hi-Fi HotStart DNA polymerase |
| 1.5* | oPool [0.05 µM] (or PCR-grade water) |
| **50** | **Total** |

In each case, the diluted oPool resuspension was not added to the reaction mix and was instead added individually in each reaction tube. In addition, one control without the oPool dilution was performed for each primer combination. Both primer pairs were designed to extend the 20 bp homology regions included in the oPool, such that mutated F3 and F4 featured 60 bp homology arms upstream and downstream of the corresponding genomic region. The two pairs were CLOP299-C3 / CLOP299-C4 and CLOP299-D3 / CLOP299-D4, respectively to amplify F3 and F4 oPools.

#### Genomic insertion of mutated fragments in *S. cerevisiae*

To insert promoter mutations upstream of *FCY1*, competent yeast cells harboring a *natNT2* deletion of the F3 or F4 region (strains SA0129 and SA0130) were transformed with the corresponding PCR-amplified oPools. For each mutation category and each region, four replicate transformations were performed from the corresponding four PCR reactions described above – for a total of 24 transformations. The protocol described by Ryan *et al.* (2016)^96^ was followed (see *CRISPR-Cas9 yeast transformation*, above). After incubation, G418-positive colonies from each agar plate were collected separately. Liquid YPD (5 ml) was added to the plate, cells were resuspended by scraping with a glass rake and the suspension was transferred to a 15 ml Falcon tube. From this suspension, glycerol stocks (687 μl cells mixed with 313 μl sterile [80%] glycerol) were made for each transformation and stored at −80°C. Two 500 μl aliquots from each tube were spun down and stored at −20°C after aspiration of the supernatant. From these samples, mutant libraries were validated by PE150 sequencing on an Illumina MiSeq instrument (Genomics Analysis Platform [PAG] of Institut de Biologie Intégrative et des Systèmes, Université Laval, Québec). This approach was also employed to insert the same set of promoter mutations upstream of the 42bp-mEGFP construct, using strains SA0246 and SA0254. In this case, three replicate transformations were instead performed per amplified oPool.

#### Selection of coding sequence mutations

Fifteen codon positions of the *FCY1* gene, for which all possible states had previously been generated by DMS^36^, were selected to obtain a diversity similar to that of the complete mutational scanning of the F3-F4 region (15 positions x 64 codons = 960 mutants). Among these positions, six were chosen for their wide range of variant effects and nine others were selected randomly, to fully cover the previously reported range of mutational effects. The 15 corresponding pools of mutants were each grown for ∼24 hours in 4 ml SC complete media (30°C with shaking), from 25 μl of glycerol stock. Optical densities were measured and all cultures were mixed in equal amounts to make four replicate 1 ml glycerol stocks (687 μl mixed cultures and 313 μl sterile [80%] glycerol) of this pool of Fcy1p mutants, which were stored at-80°C.

#### Pooling of mutants

The different sets of promoter mutations (substitutions, insertions and deletions, separately for F3 and F4) were grown separately in 4 ml SC complete media, from 50-100 μl of the corresponding glycerol stock. After ∼16 hours of incubation at 30°C with shaking, 𝑂𝐷_595_ was measured for all cultures. Subpools were then mixed to obtain equal proportions of all single mutants, according to their 𝑂𝐷_595_ and how many unique genotypes they each contained.

For mutations in the *FCY1* background, the four replicate pools of coding sequence mutations were also grown from 75 μl of glycerol stock. All subpools were then combined as described, according to their replicate number (i.e. replicate 1 for F3 substitutions was mixed with replicate 1 of all other subpools). Each subpool of promoter mutations was thus included in only two final pools: one with added coding sequence mutations and the other without. For mutations in the 42bp-mEGFP background, all subpools of promoter variants were combined as described without the addition of other mutants, except that there were only three replicates of each.

### Bulk competition experiment in 5-FC media

Each mutant pool was grown in two replicates in 4 ml SC complete media (30°C, 200 rpm) for ∼20 hours, from 111 μl or 65 μl of the corresponding glycerol stock. These inoculum volumes were chosen to have at least 2000x cells for each mutant, respectively for pools with and without added coding sequence mutations. The competition experiment was initiated by diluting each preculture at final 𝑂𝐷_595_ of 0.01 in 25 ml of SC complete media (with and without 5-FC [12 μM]) in 50 ml Falcon tubes. T0 samples were also prepared by centrifuging two microtubes containing 5 𝑂𝐷_595_ units for each preculture (850 g for 5 minutes; 4°C). After aspiration of supernatant, cell pellets were stored at −20°C. Newly inoculated cultures were incubated (30°C with shaking) in the dark until 𝑂𝐷_595_ reached ∼1.0. At this first timepoint T1, each culture was diluted in at final 𝑂𝐷_595_ of 0.01 in 25 ml of fresh media. Additional cell pellets were prepared from the remaining cultures. The full volume was first centrifuged at 4°C (1000 g; 5 minutes). Cell pellets were resuspended in ∼5 ml of supernatants and two samples of ∼5 𝑂𝐷_595_ units were prepared as previously. This process was repeated at timepoint T2 when the new cultures reached 𝑂𝐷_595_ ∼1.0 and final T3 samples were prepared when the newly inoculated T2-T3 cultures reached the same 𝑂𝐷_595_. The timings of each timepoint and the corresponding 𝑂𝐷_595_ values for all mutant pools are listed in **Supplementary Table 6**.

### Sort-seq experiment

One final pool of F3-F4 mutants in the 42bp-mEGFP background was grown overnight from 100 μl glycerol stock in 5 ml of SC complete [pH = 6.0] media, at 30°C with shaking. After measurement of its OD_595_, the culture was diluted to 0.15 OD/ml in 20 ml of fresh SC complete [pH = 6.0] and incubated for 5 hours at 30°C with shaking, to reach exponential phase (∼0.5 OD/mL). From this, 4.5 ml were transferred to a 5 ml polystyrene round-bottom Falcon tube. The remaining 15.5 ml were centrifuged to collect cells, so that genotype frequencies before sorting could also be measured by sequencing.

A FACSMelody cell sorter (BD Biosciences) was used to sort yeast cells based on their fluorescence intensity. Five bins were defined along the distribution of 𝑙𝑜𝑔_10_ fluorescence intensities, numbered 0 to 4 from the lowest to the highest fluorescence levels (**Supplementary Methods**). More cells were collected from the most populated parts of the distribution (50 000 cells each from bins 2 and 3, versus 10 000 cells each from bins 0,1 and 4). Each bin was sorted twice, resulting in two technical replicates, in a 96 well plate. The resulting fractions were each used to inoculate 3 ml of SC complete [pH = 6.0] media in a 24 deep-well plate and grown for 24h at 30°C with shaking (400 rpm). Each culture was spun down at 3000 rpm (1000 g) for 5 min and all pellets were washed with sterile water and transferred to a 1.5 ml microtube. All microtubes were centrifuged at 15 000 rpm (21 000 g) for 1 min, supernatants were removed and pellets were frozen at −80°C. This process was repeated two times for the two other biological replicates, such that six replicate samples were obtained for each bin.

### DNA library preparation and sequencing

The same approach was used for all sequencing libraries, for both the bulk competition and sort-seq experiments. Genomic DNA was first extracted from each cell pellet using a standard phenol-chloroform extraction protocol^99^, including RNAse treatment. For the resequencing of bulk competition samples in PE300, an additional RNAse treatment (10 minutes, 37°C) was performed on previously purified DNA samples, followed by precipitation with sodium acetate and cold ethanol. A first PCR was then performed using Q5 High-Fidelity DNA Polymerase (New England Biolabs) to amplify the genotyping region chosen for each type of samples, which included the F3-F4 promoter regions and flanking sequences of varying lengths. This PCR step added Illumina primer-binding sites at both ends, separated from the fragment of interest by small numbers of N bases (between 2 and 8; 10 total per PCR amplicon). For the initial (PE150) sequencing of bulk competition samples, primers CLOP311-C6 to CLOP314-H9 were used (as ordered in **Supplementary Table 2**). Two reactions were made from each DNA extraction from cultures with added coding sequence mutants. One was specific to the F3-F4 pool while the other targeted only the added *FCY1* variants. This was possible because the pool of coding sequence mutations was generated from a codon-optimized sequence, harboring many nucleotide changes from the WT ^36^. For the PE300 resequencing of the same samples, only F3-F4 PCR were performed, using oligos CLOP328-A6 to CLOP328-H7. Finally, for the sequencing of sort-seq bins, also in PE300, oligos CLOP328-A6 to CLOP341-A10 were employed. In this case, 7 N bases were added per amplicon.

In all cases, amplicons from this first PCR step were purified with AMPure XP magnetic beads (Beckman Coulter Life Sciences) and diluted to 0.01 ng/µl. These dilutions were used as template for a second set of PCR reactions using Q5 polymerase, which added Nextera barcodes and Illumina adapters i5 and i7. A unique combination of i5 and i7 indexes was used for each DNA sample. The final PCR products were also purified with AMPure XP beads, before quantification using a NanoDrop spectrophotometer (Thermo Fisher Scientific). Amplicons from the same sequencing round (PE150 competition, PE300 competition or PE300 sort-seq) were mixed in equal proportions based on the NanoDrop quantification and the resulting pool was sent for sequencing. PE150 sequencing was performed on a NovaSeq Illumina instrument (sequencing platform of Centre de recherche du CHU de Québec-Université Laval, Québec), while PE300 sequencing was performed on an AVITI instrument from Element Biosciences (Genomics Analysis Platform [PAG] of Institut de Biologie Intégrative et des Systèmes, Université Laval, Québec).

### Analysis of sequencing reads

Following each sequencing run, reads were received already demultiplexed, according to the sample id associated with each combination of i5 and i7 adaptors used. Read quality was first assessed using FastQC (v 0.12.1) and MultiQC (v 1.12)^100^. Paired-end reads were next merged using PANDAseq (v 2.11)^101^, with the ‘-k’ parameter set to 4. The N positions which had been added at both ends during the first PCR step of library construction were trimmed using command “fastx_filter” of VSEARCH (v 2.28.1)^102^, according to a table listing the number of N added for each sample. The first and last 20 bp of the resulting sequences, which were the binding sites for the oligonucleotide primers used in the first PCR step, were also trimmed using the same command. Following these trimming steps, all identical sequences were aggregated with the “derep_fulllength” command of VSEARCH. Each unique sequence was aligned to the corresponding reference (depending on the type of sample and the length of the sequenced region) using the needle aligner from the EMBOSS suite (v 6.5.7)^103^. For promoter (F3-F4) libraries, the opening and extension of gaps was equally penalized (4,4), while penalties of 50 and 0.5 were respectively used for opening and extension for *FCY1* coding sequence libraries^36^. Custom Python functions were then used to call variants from each alignment. In addition to WT reads, only sequences harboring single-nucleotide mutations (F3-F4 libraries) or single-codon substitutions (*FCY1* libraries) were kept for downstream analysis. These variants were further filtered by length, to remove those that differed too much from the expected length of the PCR fragment. For bulk competition samples, read counts of each single mutant were converted to WT-relative frequencies. These read counts were instead used as is in the downstream analysis of sort-seq samples.

### Estimation of fitness effects

#### Calculation of selection coefficients

For each bulk competition culture, time series of WT-relative genotype frequencies were reconstituted from variant frequencies obtained at the four timepoints (T0 to T3). The number of generations associated with each timepoint was obtained from the corresponding durations in hours, which varied between cultures, and the doubling time of the WT strain in the same conditions (SC media with or without 5-FC [12 μM]), as estimated from growth curve experiments. For each single-mutant genotype, the selection coefficient 𝑠 was obtained by fitting a linear model using the number of generations and the log-transformed WT-relative frequencies as dependent and independent variables, respectively. This was performed through ordinary least squares regression, using the “ols” implementation of the statsmodels (v 0.12.2)^104^ Python module, with an intercept added to the model. Regressions were performed iteratively over all time intervals which included the T0 (T0-T1, T0-T2, T0-T3; with intermediate data points when applicable). Because selection coefficients were inferred separately within each culture replicate, as many replicate values as there were cultures were obtained for each of the intervals mentioned above.

To identify the selection coefficients which should be used in further analysis, two diagnostics were performed. First, the agreement between the initial (T0) log-scaled WT-relative frequencies and the fitted intercept from each regression model was assessed. Second, the linearity of log-scaled frequencies through time, as expected for cultures growing exponentially, was validated graphically. This revealed that no interval was reliable for the initial 150 paired-end genotyping of promoter (F3-F4) libraries, while T0-T2 selection coefficients should be used for the 300 paired-end resequencing of the promoter fragments and the 150 paired-end sequence of *FCY1* coding sequence (see Supplementary Note). The chosen selection coefficients were then used to identify a T0 abundance threshold for variant filtering, such that inferred 𝑠 would not be correlated with initial read counts. This threshold was set at 85 reads, both for the promoter (300 paired-end) and coding sequence (150 paired-end) sequencing libraries. To ensure that low frequency but beneficial secondary mutations could be assessed, any variant which was below the threshold at T0 but over it at the end of the experiment was kept.

#### Removal of outliers and aggregation of replicates

In 5-FC samples, mutations sometimes appeared beneficial in only one pair of cultures (coming from the same transformation replicate), likely due to secondary resistance-conferring mutations (See Supplementary Note). Outlier removal was thus performed, separately for two types of libraries (promoter or coding sequence). The promoter (F3-F4) libraries were further separated depending on whether coding-sequence mutants had been added in the corresponding cultures. Within each group of samples, the median absolute deviation (MAD) of selection coefficients was first computed for each mutant. This was then used to compute robust z-scores for each replicate measurement within each genotype. The resulting distribution of robust z-scores calculated across all mutant genotypes was compared and its 95th percentile was defined as the threshold to identify outlier replicates. The filtering of outliers was however restricted to a subset of mutants which displayed a higher variance, to ensure that small absolute variations between replicates were not identified as outliers. This was assessed from each genotype’s standard deviation, with the 95th percentile of the overall distribution used as threshold. For F3-F4 sequencing libraries obtained from cultures with added coding sequence mutations, the 85th percentile was instead used.

To obtain final estimates of the fitness effects of promoter mutations, only F3-F4 libraries without added coding sequence mutants were taken into account. Four of the eight replicate cultures, corresponding to two of the four transformation replicates, were additionally ignored. In these four samples, selection coefficients displayed a temporal shift towards negative values which appeared consistent with some undetected resistance-conferring mutations being genotyped as WT (see Supplementary Note). For each of the 1020 assayed promoter mutations, the corresponding fitness effect was estimated as the median of the selection coefficients observed across the four remaining replicates, which were considered unbiased.

The statistical significance of selection coefficients was assessed from the same four replicate measurements for each mutation, by comparing their mean to 0 using the SciPy (v 1.15.2)^105^ implementation of the one-sample t-test, followed by Benjamini-Hochberg correction controlling the false discovery rate at 5%. For the coding sequence mutations added as controls, fitness effects were instead estimated as the median of the eight replicate measurements, without ignoring any of the corresponding samples. Alternatively, significant selection coefficients were also defined according to thresholds of minimal absolute deviations from 0, using the most extreme positive and negative replicates for each genotype. In this case, the number of fitness changes identified was compared with a null expectation derived from the observed experimental noise. To this end, the complete dataset of final selection coefficients (from the four culture replicates mentioned above) was shuffled into 1020 random “genotypes”. From each culture replicate, 1020 selection coefficients were first sampled with replacement. The resulting four sets of values were then combined randomly into the 1020 genotypes, such that each was associated with four replicate measurements. The number of random genotypes exceeding the chosen threshold of absolute deviation from 0 was then compiled. This was repeated 10 000 times.

### Comparison with published effects of amino acid substitutions in Fcy1p

The selection coefficients of added coding sequence mutations were first aggregated into grand medians, according to the corresponding amino acid substitutions. They were then compared with previously published variant effect scores obtained in the same conditions (12 μM 5-FC)^36^. The agreement between the two experiments was further assessed by classifying variants as resistant and susceptible in both cases. To this end, Gaussian mixture models with two components and a “tied” covariance matrix were fitted on each of the two datasets, using the scikit-learn (v 1.6.1)^106^ implementation.

For the comparison of promoter and coding sequence mutations, the full dataset from^36^ was used. All single-nucleotide substitutions in the reference sequence of the *FCY1* gene, obtained from the Saccharomyces Genome Database^98^, were first performed computationally. The corresponding variant effects were retrieved from the Després *et al.* (2022) data^36^, according to the resulting amino acid substitutions. Of 1431 possible single-nucleotide substitutions in *FCY1*, only 20 were not covered, mostly in the first and last codons. The Gaussian mixture model previously fitted on the full set of amino acid changes was then used to classify all single mutants as either resistant or susceptible. From this classification, the fraction of adaptive (resistance-conferring) mutations in the coding sequence was obtained.

### Calculation of sort-seq expression scores

Within each of the six replicate sort-seq experiments, all variants which reached a total abundance of at least 85 reads across the five sampled bins were kept for downstream analysis. Although this included a small number of secondary mutations (outside of the F3-F4 mutagenized region), any variant which occurred in the mEGFP sequence was filtered out. For each of the remaining genotypes *i*, which includes the WT, fluorescence intensity was quantified from a weighted average *B_i_* of bins where the corresponding reads were detected:

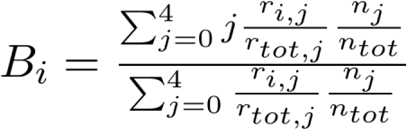

In the previous formula, *j* is the bin number, where 0 and 4 are associated with the lowest and highest fluorescence intensities, respectively. Variables *r_i.j_* and *r_tot.j_* are the read count for genotype *i*, as well as the total number of reads, in bin *j*. Finally, *n_i_* and *n_tot_* are respectively the number of cells collected from bin *j* and the total amount of cells collected across the five bins. The *n_i/_ n_tot_* ratio was introduced because the number of yeast cells collected was not uniform across all bins.

Separately for each replicate experiment, the difference was computed between the *B_m_* weighted average of each mutant *m* and the *B_WT_* weighted average obtained for the WT genotype:

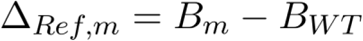

For each mutant, up to six replicate Δ*Ref* scores were obtained (one for each sort-seq experiment). The median of these values was used as an estimate of each mutation’s effect on fluorescence intensity. Positive values denote an increase relative to the WT, while negative values instead indicate a decrease.

### Selection and construction of validation mutants

The sort-seq data was used to select 30 mutant genotypes for validation. These mutations were chosen to cover the whole range of scores, as well as all regions of the promoter where single-nucleotide changes appeared impactful. Substitutions were favored to ensure a high success rate to the downstream reconstruction of genotypes. To ensure that low-confidence mutations were not selected, a one-sample t-test (with Benjamini-Hochberg correction at FDR=5%) was first performed on the replicate sort-seq scores to identify Δ*Ref* values which were significantly different from 0. This was also repeated with Δ*mod* scores computed from Δ*mod* weighted averages obtained from reads of bin 1 to 4, excluding the 0th one displaying the lowest fluorescence intensities. Because the Δ*Ref* scores appeared biased towards positive values compared with Δ*mod*, the latter were used to select significant mutations to be reconstructed, except when annotated *. This ensured a more stringent selection of genotypes displaying altered fluorescence intensity compared with the WT reference. Within each of the mutant groups described below, the specified number of sequences were chosen randomly, unless stated otherwise. Among mutations which significantly decreased Δ*Mod* below zero, there were seven groups: 1) indels before the second methionine of *FCY1* (n=2); 2) indels after the second methionine of *FCY1* (n=2)*; 3) top 3 lowest scores between positions −50 and 0 of the promoter*; 4) substitutions between positions −133 and −127 (n=6); 5) substitutions between position −45 and −11 (n=2); 6) substitutions and insertions near the −75 position (one each) and 7) two chosen substitutions outside of regions with many significant effects. Among mutations which significantly increased Δ*Mod* above zero, a similar approach was employed. The four categories were: 1) substitutions between position −125 and −118 (n=4); 2) substitutions between positions −112 to −109 (n=2); 3) substitutions between positions −63 to −57 (n=2) and 4) variants among the 25 highest scores (n=4).

The corresponding validation mutants were individually reconstructed using the CRISPR-Cas9 yeast transformation protocol described above. For each genotype, this was performed in two genetic backgrounds: fluorescent strain SA0247 (ΔF3-F4 with 42bp-mEGFP construct) and *FCY1* deletion (ΔF3-F4) mutant SA0159 (see **Supplementary Table 1**). A fusion PCR product harboring the single mutation of interest was used as donor DNA. This amplicon was generated by two consecutive rounds of PCR. In the first round, two reactions were performed with WT genomic DNA as template. The first one used a generic forward primer hybridizing upstream of F3 (CLOP326-A11) with a mutant-specific reverse primer harboring the mutation of interest (**Supplementary Table 2**). In the second reaction, a forward primer with the mutation of interest (**Supplementary Table 2**) was used alongside a generic reverse primer binding either in the mEGFP sequence (CLOP326-G10; for background SA0247) or in the *FCY1* coding sequence (CLOP326-E10; for background SA0159). The two mutant-specific mutagenesis primers are complementary to each other, such that there is homology between the two amplicons. The second PCR round used the two PCR products from the first round diluted 1/40 as template and the same two generic primers used in the first pair of PCR (CLOP326-A11 with either CLOP326-G10 or CLOP326-E10, depending on the mutagenized background), which allowed fusion of both PCR parts. This approach thus allows to amplify the full F3-F4 promoter region (with a short segment of *FCY1*) while introducing a single mutation within it and adding homology regions on both ends for subsequent genomic insertion. It was additionally used to perform deletion of short segments of the promoter (−133 to −127 and −125 to −118), similarly in both genetic backgrounds. After transformation of each fusion PCR product, clones from each transformation were sent for Sanger sequencing to validate proper genomic integration of the mutation. All 66 resulting strains, including reconstructions of the WT promoter, are listed in **Supplementary Table 1**.

### Expression measurements by flow cytometry

Measurement of promoter activity using fluorescent strains was performed with a Guava EasyCyte HT cytometer (CytekBio). Each strain of interest was grown overnight in three replicates in SC complete media at 30°C with shaking, along with similar replicate cultures of a reference strain with the WT promoter (SA0247 or AKD1462) and of a non-fluorescent control (AKD1123 or AKD1429). The overnight cultures were diluted at 0.15 𝑂𝐷_595_ in fresh SC-Trp (MSG) media and grown at 30°C with shaking until ∼0.5 𝑂𝐷_595_. Exponentially growing cells were then diluted to 0.05 *OD*_595_ in fresh media and single-cell fluorescence intensity measurements were performed with the Guava EasyCyte HT system. For each replicate culture of each strain, 5000 events were acquired (488 nm excitation; 525 nm detection). The medians of the corresponding distributions of fluorescence intensities were used to estimate relative expression levels.

Within each cytometry experiment, events were first filtered out according to forward scatter and side scatter values, with the 99th percentile of each distribution used as thresholds. All fluorescence intensities were then normalized by the corresponding forward scatter measurement. From these normalized fluorescence intensities, a median was computed for each replicate culture assayed. These medians were then normalized by the grand median of measurements made for the reference strain, such that they were converted into WT-relative fluorescence intensities. The resulting values, and especially their medians, were used as estimates of relative expression levels. Accordingly, three replicate measurements of WT-relative fluorescence intensity were obtained for each tested strain, with a value of 1 being equal to the WT level.

### Calibration of sort-seq scores and bootstrapping of relative expression levels

WT-relative fluorescence intensities were first measured for all validation mutants reconstructed in the SA0247 (42bp-mEGFP) background, as described above. These measurements were then compared with the corresponding sort-seq expression scores and a LOWESS regression was fitted. This approach allowed to simultaneously validate the sort-seq experiments and convert the resulting scores into more interpretable WT-relative fluorescence intensities. The Δ*Ref* scores were used, since some genotypes with very low fluorescence intensities were absent from the Δ*mod* dataset. In addition, because fluorescence intensity measurements from reconstructed mutants were used to fit the relationship, any bias in Δ*Ref* was unlikely to affect the downstream inference of relative expression levels. So that uncertainties could also be obtained for each of the >1000 mutant expression levels, both the LOWESS fitting and the conversion of sort-seq scores into relative fluorescence intensities were performed within a bootstrapping routine.

At the start of each iteration, Δ*Ref* scores were first resampled with replacement. For each genotype which had been included in the sort-seq analysis, as many values were sampled as there were replicate measurements among the six experiments. Per-genotype medians were then computed on the newly sampled data. For each reconstructed mutant for which fluorescence had been measured directly, three median WT-relative fluorescence intensities were sampled with replacement among the corresponding replicate measurements. Grand medians were computed from each trio of observations, which were combined with the newly recomputed median Δ*Ref* scores of the corresponding validation genotypes to obtain 30 pairs. From these data points, LOWESS regression was performed with median Δ*Ref* as independent variable, using the “nonparametric.lowess” implementation from statsmodels (v 0.14.4)^104^ with default parameters. This regression was then evaluated at each of the >1000 median Δ*Ref* values obtained from the first resampling steps. This generated an estimate of WT-relative fluorescence intensity for each of the genotypes which had been assayed in the sort-seq screen. The regression was also evaluated at 100 equally spaced values from the minimal to the maximal median Δ*Ref* observed experimentally. This process was repeated 5000 times. After all iterations had been completed, medians and corresponding 95% prediction intervals were computed from the distribution of WT-relative fluorescence intensities obtained for each mutant genotype. These medians were used as WT-relative expression levels associated with the corresponding F3-F4 mutations. The same calculations were performed for each of the 100 equally spaced values at which the LOWESS had been evaluated, thus allowing to plot the relationship between Δ*Ref* and relative fluorescence intensity.

### Experimental measurements of the expression-fitness function

#### Fitting of dose-response curves

The WT background AKD1123 was phenotyped in liquid media containing various concentrations of 5-FC and cytosine, according to the growth assays protocol described above. From two to three replicates were made for each concentration of 5-FC and cytosine. A five-parameter logistic model was then fitted, using the exponential growth rate measurements and the concentrations of 5-FC / cytosine as dependent and independent variables, respectively. The resulting curves were converted into relative inhibition or relative growth, using measured growth rates in absence of 5-FC or at the highest cytosine concentration as references.

#### Construction of strains with distinct *FCY1* expression levels

Six yeast promoters covering a wide range of expression levels were used to construct two sets of strains, respectively from *FCY1* background AKD1123 and from fluorescent background SA0228. The six promoter cassettes, each including a G418 resistance marker, were PCR-amplified from the corresponding plasmids^95^ (**Supplementary Table 3**) using oligonucleotides CLOP338-G6 and CLOP338-H6 (**Supplementary Table 2**). The two primers provided homology for genomic insertion immediately upstream of *FCY1* (or of its first 42 bp for the SA0228 background), between the gene and its native promoter. Yeast transformations were performed in the two genetic backgrounds as previously described, using each of the corresponding PCR products. Transformants were selected on YPD+HYG+G418 media. Correct integration of the promoter cassette was validated by PCR in four clones from each transformation, using oligonucleotides CLOP310-D1 and CLOP291-A3 (**Supplementary Table 2**). Two to three positive clones were obtained in most cases and validated by Sanger sequencing, such that biological replicates could be used in downstream assays for the majority of genotypes. All 25 corresponding yeast strains are listed in **Supplementary Table 1**.

#### Measurements of growth rates and fluorescence intensities

The set of *FCY1* strains (background AKD1123) was used to measure growth rates in liquid media containing different concentrations of 5-FC or cytosine, following the growth assays protocol described above. The set of fluorescent strains (background SA0228) was instead used in flow cytometry experiments to measure expression levels (see Expression measurements by flow cytometry). In both cases, all available biological replicates were used for each genotype. When less than three strains were available, three replicate cultures were nonetheless inoculated.

The growth assays in 5-FC media were performed in more than one experiment. To combine measurements obtained for concentrations of 12 μM and 3 μM (**Extended Data Fig. 9**), a normalization step was first performed.Growth rates from the first experiment were adjusted to have the same grand medians as those of the second one, by the addition of the difference between the two experiments’ grand medians. This was performed separately for each concentration and allowed to combine the two sets of growth rates, in spite of potential systematic differences between them – for instance if the true concentration of 5-FC in the media varied. The cytometry measurements, which were done in parallel with the growth assays, were also performed in two separate experiments. In this case, potential systematic differences were controlled for by computing WT-relative fluorescence intensities within each experiment and only combining measurements after this step.

#### Fitting of expression-fitness functions

Fluorescence measurements associated with each promoter were combined with the corresponding growth rates. Measurements made from the AKD1123 WT strain were however not used, because this strain lacks the G418 resistance marker included in promoter cassettes and is thus not directly comparable. An exponential function was fitted for each condition, using WT-relative fluorescence as the independent variable and growth rate as the dependent variable. This could model the sharp decline observed in 5-FC as well as the steep increase seen in cytosine conditions. The fitting of all expression-fitness curves was performed within a bootstrapping routine. For each promoter, two WT-relative fluorescence intensity measurements were first sampled with replacement from each cytometry experiment. Then, the four resulting values were resampled with replacement. Corresponding growth rates were sampled similarly. If a given condition had been tested in two experiments, two replicate growth rates were sampled with replacement from each experiment for each promoter, and the four measurements were afterwards resampled with replacement. This approach was employed to avoid favoring one experiment over the other, since the number of replicates sometimes varied between them. When only one experiment had been performed, replicate growth rates from this assay were instead resampled with replacement. Once relative fluorescence intensities and growth rates had been sampled, values obtained for the same promoter were paired randomly into data points. The exponential function was then fitted using the “optimize.curve_fit” implementation of the SciPy module (v 1.15.2)^105^ module and evaluated over 250 equally spaced values covering the full range of WT-relative fluorescence intensities. After 5000 iterations, medians and corresponding 95% prediction intervals were computed for each of these 250 fluorescence levels, thus generating the expression-fitness function. This was repeated for each of the 5-FC and cytosine concentrations tested.

#### Inference of the expected distribution of effects on fitness for promoter mutations

Changes in *FCY1* expression were assumed to have the same effect on fitness as variations in 5-FC concentration. The dose-response curve (**Extended Data Fig. 2**) was converted such that concentrations were expressed as relative changes from 12 μM (used in the bulk competition experiment). From this new curve, corresponding exponential growth rates were inferred for the 1019 relative expression estimates obtained from the sort-seq experiment. All growth rates were then converted into selection coefficients, as described above.

### Validation of *FCY1* promoter variants fitness effects

The previously described validation mutants reconstructed in the SA0159 *FCY1* background (**Supplementary Table 1**) were grown in SC complete (MSG) media with 12 μM 5-FC and without, according to the growth assays protocol described above. For all other mutants, growth rates measured in 12 μM 5-FC were converted into selection coefficients as described in the growth assays section, using growth rates measured in the same experiment for CRISPR reconstructions of the WT genotype in the SA0159 background (generated at the same time as the validation mutants). The expression-fitness function obtained in the same conditions and the bounds of its 95% prediction interval were also converted into selection coefficients using the same approach. In this case, the WT growth rate interpolated from the curve was instead used as comparison.

### Assessment of 5-FC resistance arising from mutant library construction

To test whether mutations conferring 5-FC resistance were introduced during mutant library construction, the process described in *Construction of mutant libraries* was repeated, but using WT sequences of the F3 and F4 regions. Briefly, the same two oligonucleotide pairs (CLOP299-C3/CLOP299-C4 and CLOP299-D3/CLOP299-D4) were first used to amplify F3 and F4 from genomic DNA of the AKD1123 strain, using KAPA HiFi HotStart DNA polymerase (Roche Molecular Systems). Competent cells from strains SA0129 and SA0130 (prepared as described by ^96^) were then transformed with the F3 and F4 inserts, respectively. Two biological replicates of each transformation were made. This was performed following the CRISPR-Cas9 approach described above for the pools of single mutations. After 48 hours of incubation at 30°C, transformants from each Petri dish were collected in 5 ml YPD by scraping, as previously described. From each of the resulting cell suspension, cultures were started in 5 ml of SC complete [pH=6] + HYG, from 100 μl of cells.

After overnight incubation at 30°C with shaking, *OD*_595_ was measured for each culture and dilutions at *OD*_595_ = 1 were made in 200 μl of sterile water. Additional dilutions were afterwards made for *OD*_595_ values from 0.2 to 0.002. From each dilution, 150 μl were plated on SC complete (MSG) solid media with and without 5-FC [100 μg/ml]. Petri dishes were incubated at 30°C for 48 hours and colonies were counted. While colonies were too numerous to be counted on SC complete controls, some dilutions plated on 5-FC media resulted in countable colony numbers (between 20 and 340). For each plating (one per CRISPR-Cas9 transformation), colony counting was performed for two dilution levels, which acted as technical replicates. Since the SA0130 background was initially resistant to 5-FC due to the insertion of a natNT2 in place of F4, all associated platings were replica plated on YPD+NAT media. Nourseothricin-resistant colonies were subtracted from the total number of 5-FC-resistant colonies. The frequency of 5-FC resistance was then estimated as follows, separately for each plating on media containing 5-FC:

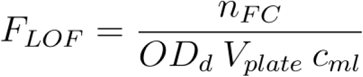

In the equation above, 𝑛_𝐹𝐶_ is the number of colonies on 5-FC media (after subtracting NAT resistant ones, when applicable), while *OD*_𝑑_ is the *OD*_595_ of the plated dilution, 𝑉_𝑝𝑙𝑎𝑡𝑒_ is the volume deposited on each plate (0.15 ml) and 𝑐_𝑚𝑙_ is the number of yeast cells in 1 ml of media at *OD*_595_ = 1, for which a value of 10 million cells/ml was used. The resulting *F_LOF_* was considered as a putative frequency of *FCY1* LOF in the corresponding transformations. To validate whether that was the case, 16 colonies were selected for Sanger sequencing, using primer CLOP326-F10, which covered nearly 70% of the *FCY1* coding sequence **(Supplementary Table 2)**. All colony counts, resistance rates and relevant calculations are listed in **Supplementary Table 7**.

### Simulations with *invisible* resistance-conferring mutations

A simulation framework replicating the bulk competition experiment was designed. It involved a library of neutral variants within which a small number of invisible resistance-conferring mutations was introduced, thus mimicking secondary mutations occurring outside of the genotyping region in the experiment. A set of variants of interest (hereafter named genotype labels) was first generated by sampling 1000 neutral mutations from *s* ∼ *N*(0, 0.004) four times. The standard deviation used was set according to what was observed in sequencing libraries from cultures without 5-FC. Four transformations were then obtained by replicate random samplings of 10 000 clones, among which 17% were defined as WT (*s* = 0) and the remaining were drawn from the set of genotype labels. Within each such transformation replicate, a small fraction of clones was defined as harboring an invisible resistance-conferring mutation (or resistance flag), which increased its 𝑠 (between 0.2 to 0.4, depending on simulations) without changing its genotype label. From each transformation, two replicate cultures were generated by randomly sampling two sets of 2.5 million cells, resulting in eight cultures, as in the bulk competition experiment (5-FC condition without added coding sequence variants). From each culture, 1 million of sequencing reads (genotype labels) were sampled randomly, to simulate sequencing of the initial T0 samples. Simulated cells in each culture were next aggregated according to genotype labels and presence/absence of a resistance flag, prior to calculation of WT-relative frequencies for each group (called true genotypes). Growth over the course of a bulk competition experiment was simulated by modeling exponential changes of WT-relative frequencies through time for each true genotype 𝑖, as follows:

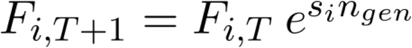

The WT-relative frequency 𝐹 of true genotype 𝑖 at timepoint 𝑇 + 1 is obtained from the corresponding WT-relative frequency at previous timepoint 𝑇 according to the genotype’s true selection coefficient (increased by a resistance-conferring mutation when applicable) and the time interval 𝑛_𝑔𝑒𝑛_ between the two timepoints, in WT generations. Three such timepoints, from T1 to T3, were simulated and their respective durations were set in accordance with the bulk competition experiment. At each timepoint, 1 million reads as well as a new initial population of 2.5 million cells were sampled from each culture, as described above. Selection coefficients for all variants were then computed from time-series of simulated read counts, as described for experimental variant frequencies. This was performed from genotype labels only as well as with knowledge of true genotypes (resistance flags). Such simulations of one full experiment were repeated while varying both the initial frequency and the fitness effect of resistance-conferring mutations. For each combination, multiple replicate simulated experiments (each with eight simulated cultures) were performed.

## Supporting information

Description of Supplementary Tables

## Data availability

All sequencing data from bulk competition and sort-seq experiments has been deposited on the NCBI SRA (https://www.ncbi.nlm.nih.gov/sra/PRJNA1321317). All other experimental data and files needed to replicate the analysis have been deposited on Zenodo: https://doi.org/10.5281/zenodo.17088129.

## Code availability

All analyses were performed using custom Python scripts and notebooks, which are available at: https://github.com/Landrylab/Aube_et_al_2025_FCY1promoter. A custom Python pipeline was deployed on a computing cluster (Institut de Biologie Intégrative et des Systèmes, Université Laval) to compute variant counts and selection coefficients from raw sequencing data. As described above, this used the following software: FastQC (v 0.12.1), MultiQC (v 1.12)^100^, PANDAseq (v 2.11)^101^, VSEARCH (v 2.28.1)^102^ and EMBOSS suite (v 6.5.7)^103^. The versions of all Python modules used are included as requirements files, separately for the analysis pipeline and for the notebooks, which were executed locally.

## Acknowledgements

We thank Philippe Després, Isabelle Hatin, Olivier Namy and all Landry lab members, especially Soham Dibyachintan, David Jordan, Pascale Lemieux and Camille Bédard, for helpful discussions. We are grateful to Pavithra Venkataraman, Angel Cisneros, Soham Dibyachintan and Philippe Després, who provided comments on this manuscript. We additionally thank Angel Cisneros for helpful suggestions on the simulation framework. This work was supported by grants from the Natural Sciences and Engineering Research Council of Canada (NSERC RGPIN-2020-04844) and from the Human Frontier Science Program (HFSP RGP011/2025) to CRL. CRL holds the Canada Research Chair in Cellular Systems and Synthetic Biology. SA was supported by doctoral research fellowships from Natural Sciences and Engineering Research Council of Canada and Fonds de recherche du Québec - Nature et technologies. The F65V-34a-mEGFP plasmid used in this study was a kind gift from Dr Michael Springer.

## Author contributions

C.R.L., A.K.D and S.A. designed the study. S.A. and A.K.D. performed the experiments as well as the data analysis. S.A., A.K.D. and C.R.L. interpreted the results. S.A., A.K.D and C.R.L. wrote the manuscript.

## Extended Data Figures

**Extended Data Fig. 1:**
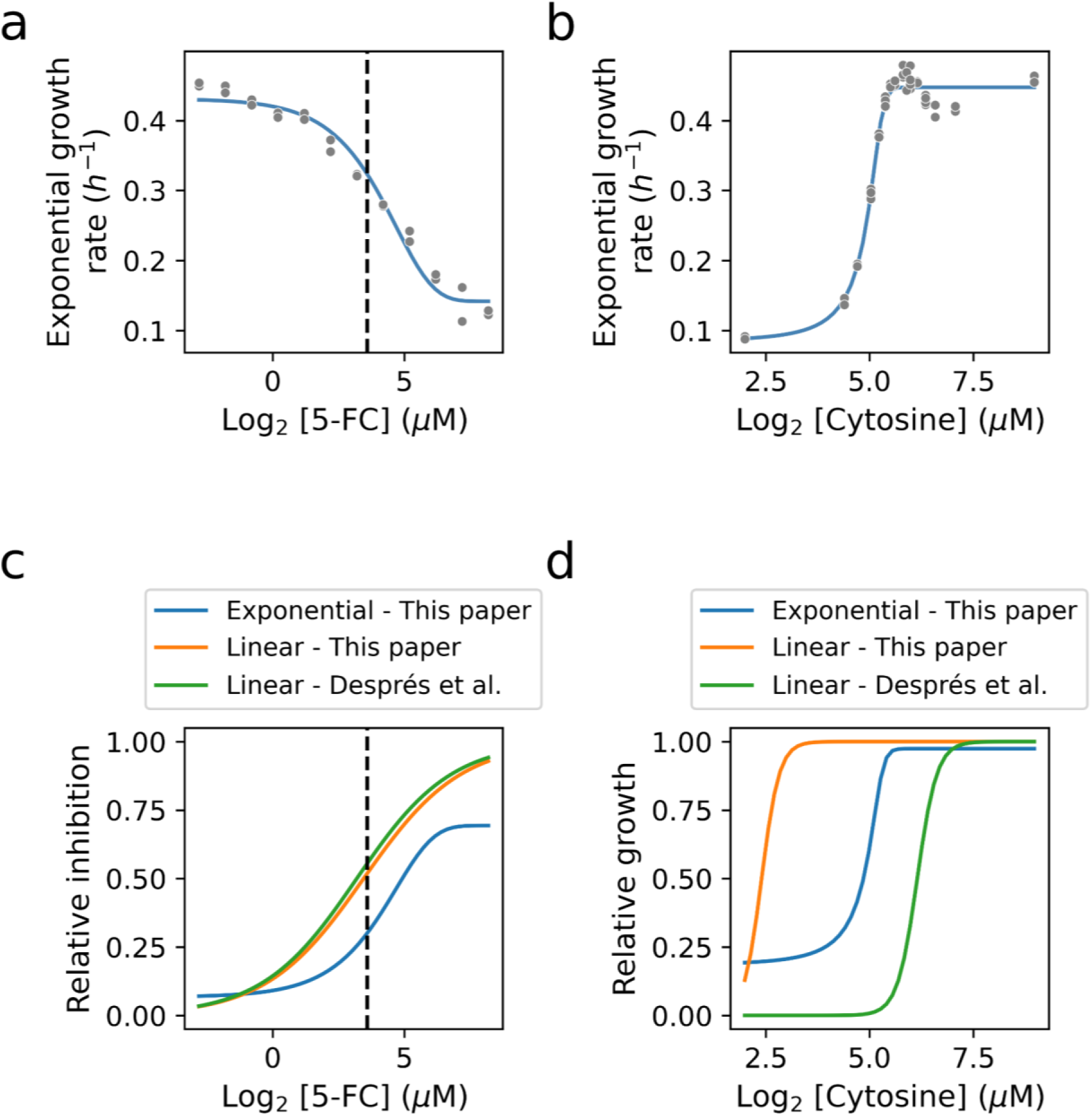
The growth rate of wild-type yeast varies sharply across a range of 5-FC and cytosine concentrations. **a,** Exponential growth rates for the wild-type reference strain AKD1123 across 5-FC concentrations. Two replicate cultures were assayed for each tested concentration (dots). The line displays a fitted five-parameter logistic function (Methods), while the vertical dashed line identifies the 12 μM concentration used in the bulk competition experiment. b, Same as panel a, but for a range of cytosine concentrations in media lacking uracil. c, The dose-response curve obtained for 5-FC is consistent with a previously published relationship. The curve from a is shown after conversion into relative inhibition levels, using growth rates measured in absence of 5-FC within the same experiment as comparison (0 relative inhibition). This curve, labeled “Exponential - This paper”, was used in downstream analyses to obtain the relative inhibition level associated with a given 5-FC concentration. When the same growth curve data is used to compute maximum linear growth rates (rather than exponential growth rates) and then fit a dose-response curve as in Després et al.^36^, the published relationship is replicated. d, Same as panel c, but using the cytosine growth data shown in panel b. Here, the published relationship cannot be replicated by using the same analysis approach. The curve, labeled “Exponential - This paper”, was nonetheless used in downstream analyses to obtain relative growth levels.

**Extended Data Fig. 2:**
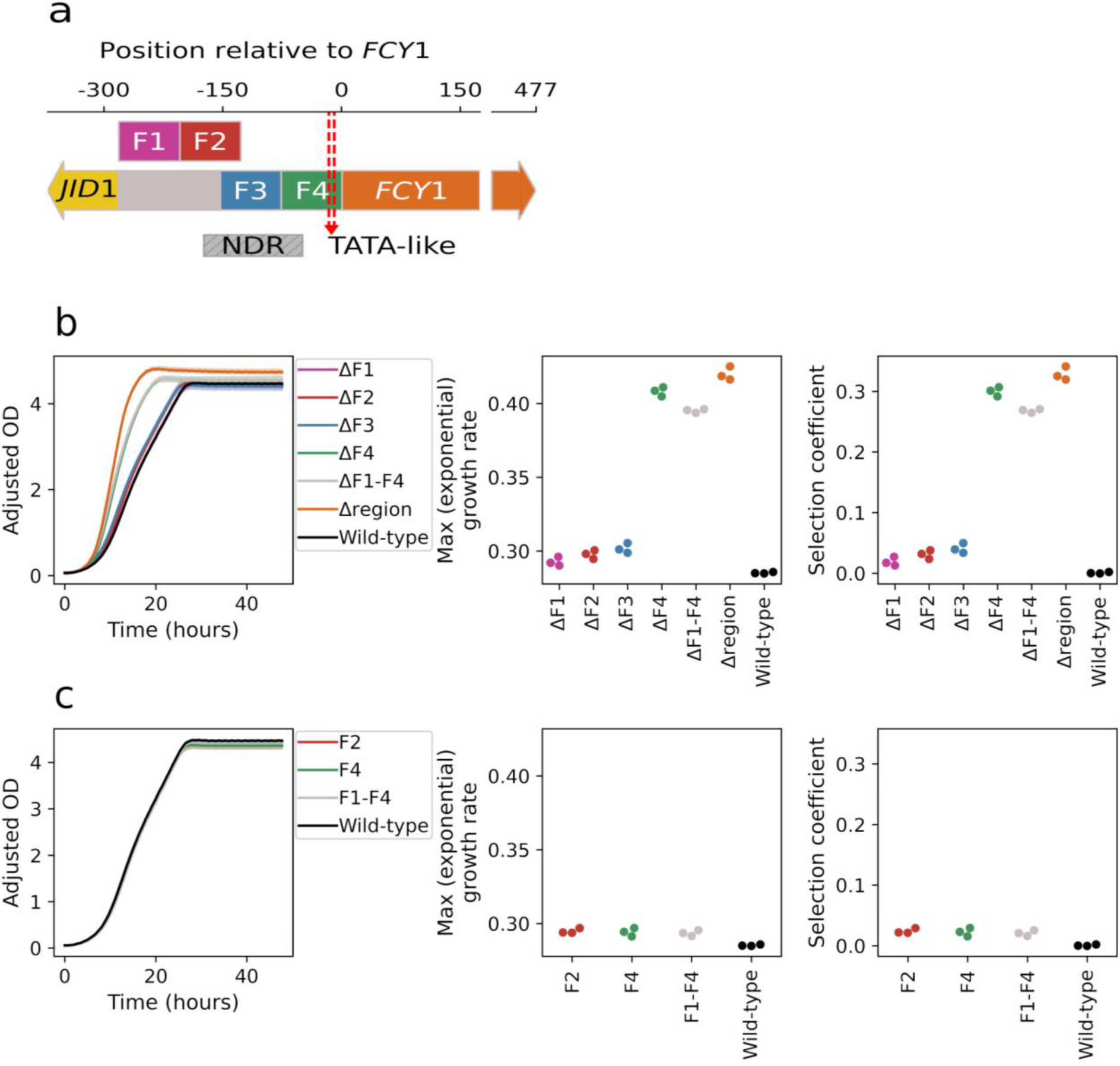
The deletion of the F4 promoter fragment is clearly adaptive in 5-FC conditions. **a,** Schematics of the four 77 bp fragments into which the intergenic region between *JID1* (YPR061C) and *FCY1* (YPR062W) was divided. There is a 26 bp overlap between F2 and F3. The nucleosome-depleted region (NDR) of the corresponding bidirectional promoter^67,107^ as well as the TATA-like element, involved in transcription initiation at *FCY1*^47,48^, are shown. **b,** Fitness of deletion mutants in [12 μM] 5-FC. The ΔF1-F4 mutant refers to deletion of the complete intergenic region, while Δregion instead represents the simultaneous deletion of the intergenic region and the *FCY1* gene. Exponential growth rates and the corresponding selection coefficients (*Methods*) show that F4 is the only promoter fragment for which deletion confers a clear growth advantage (and thus 5-FC resistance). **c,** Fitness of reconstructed WT strains in [12 μM] 5-FC. Here, F2, F4 and F1-F4 refer to CRISPR reconstruction of the WT intergenic region from the corresponding deletion mutant (*Methods*). In all three cases, the WT 5-FC sensitivity is recovered. All growth measurements were made from three replicate cultures of each strain.

**Extended Data Fig. 3:**
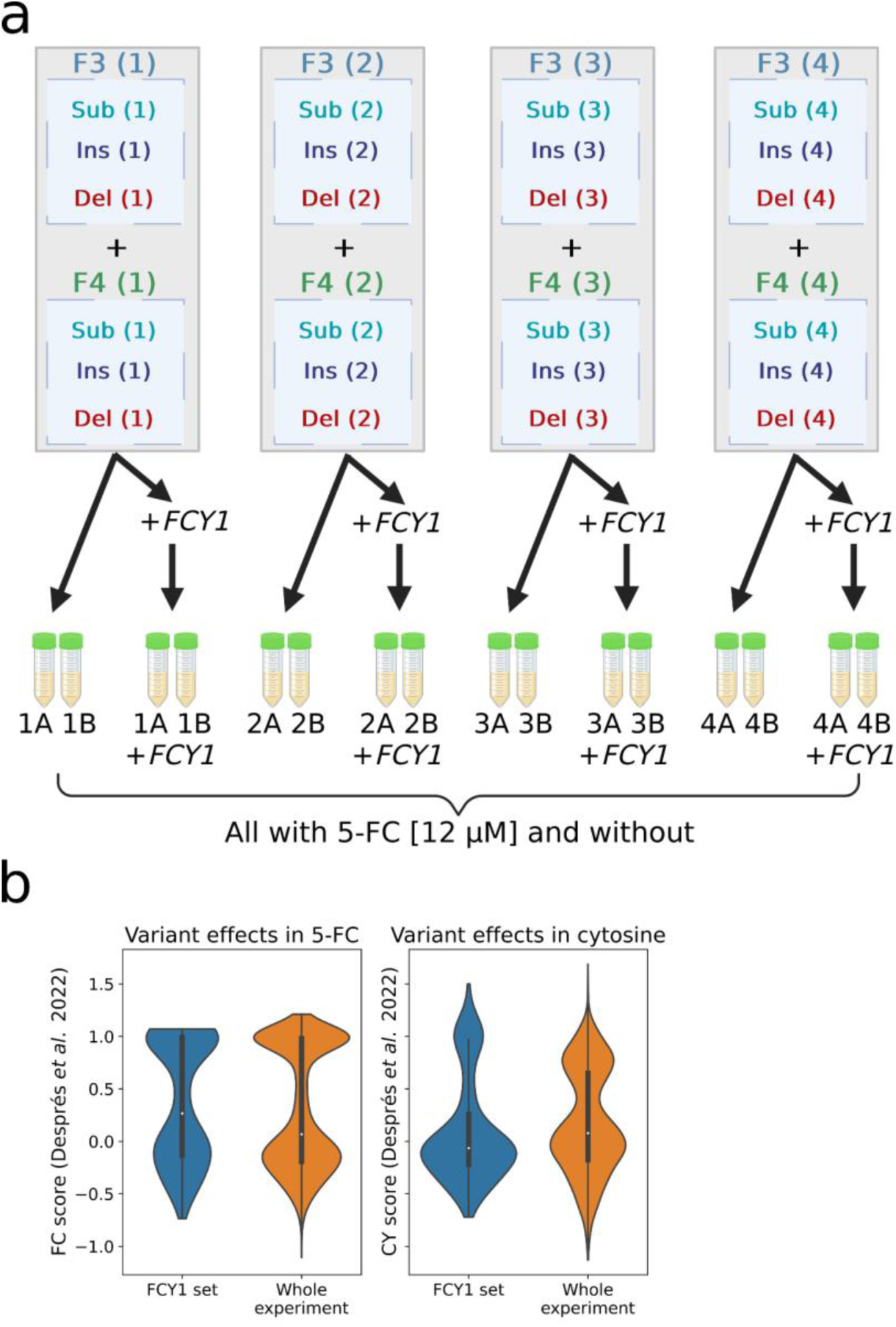
Complete overview of the bulk competition experiment. **a,** Full experimental design showing all 32 cultures and their relationships as transformation and/or culture replicates. Each pool of substitutions (Sub), insertions (Ins) or deletions (Del) was constructed four times and corresponding replicates were mixed to obtain each of the four *transformation replicates* (grey rectangles). Each transformation replicate was used to inoculate two culture replicates, with or without the addition of a set of previously characterized coding sequence mutations (*FCY1*; n=960)^36^. Created in BioRender. Aubé, S. (2025) https://BioRender.com/acpk3ks. **b,** The selected coding sequence variants cover the full range of variant effects reported in^36^, both in 5-FC and cytosine (without uracil) conditions, where Fcy1 activity is respectively toxic and essential.

**Extended Data Fig. 4:**
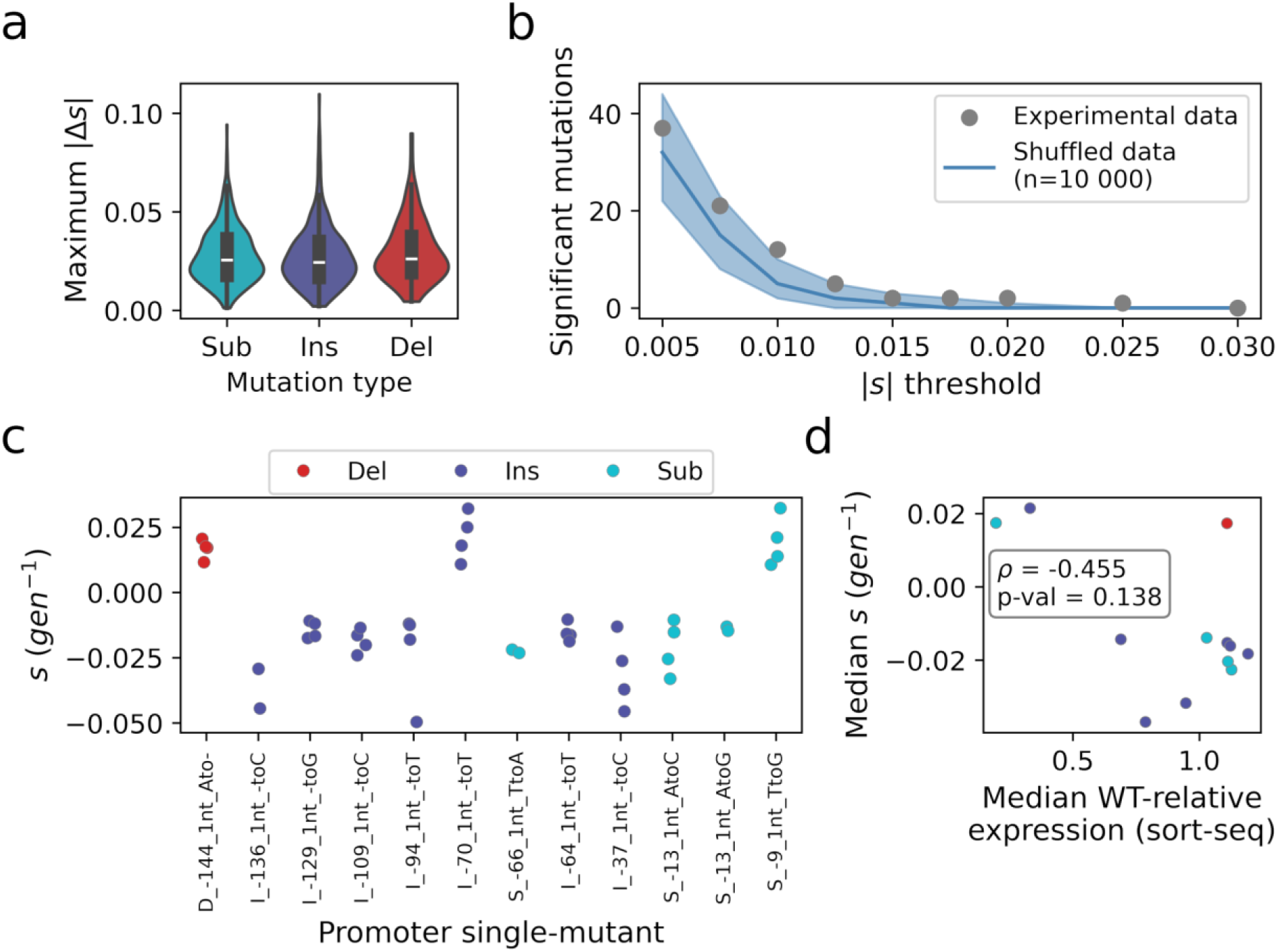
The number and magnitude of small fitness effects observed among promoter single-mutants are consistent with the level of noise in the bulk competition experiment. **a,** Selection coefficient measurements for the same genotype typically cover an interval of ∼0.028 𝑔𝑒𝑛^−1^. The distributions of maximal |𝛥𝑠|, the difference between the two most extreme selection coefficient estimates obtained across replicate cultures (n=4), are shown for all assayed single-nucleotide substitutions (Sub; n=462), insertions (Ins; n=462) and deletions (Del; n=96) in the *FCY1* promoter. b, The number of mutations exceeding different thresholds of minimal fitness effect is consistent with the expectation derived from random shuffling of the data. Across a range of minimal |𝑠|, all genotypes for which the lowest fitness estimate is above +𝑠 (significant beneficial effects) or the highest estimate is below −𝑠 (significant deleterious effects) are shown (dots). The solid line and shaded area display the median and associated 95% confidence interval for the number of significant effects (using the same definition) obtained from random permutations of the data (n=10 000). Within each iteration, 1020 mock single mutants were assembled by randomly combining selection coefficients from each of the four replicate cultures (Methods). c, Twelve promoter single-mutants display |𝑠| > 0.01 in the bulk competition experiment. For each genotype, all replicate 𝑠 estimates are shown (dots). The identifiers of the mutations (x axis) inform on their type (deletion, insertion or substitution), position relative to the *FCY1* coding sequence (where 0 is the base immediately upstream of the gene), and nucleotide change. d, The small fitness effects measured are not clearly related to corresponding changes in *FCY1* expression level. For mutants included in panel c, the median selection coefficient estimate is compared with Wt-relative expression level estimated from sort-seq (see Fig. 4e). Whereas the correlation is negative as expected, the second highest fitness effect is observed for a mutation increasing expression (>1) and the most negative selection coefficient is associated with a moderate decrease of *FCY1* expression.

**Extended Data Fig. 5:**
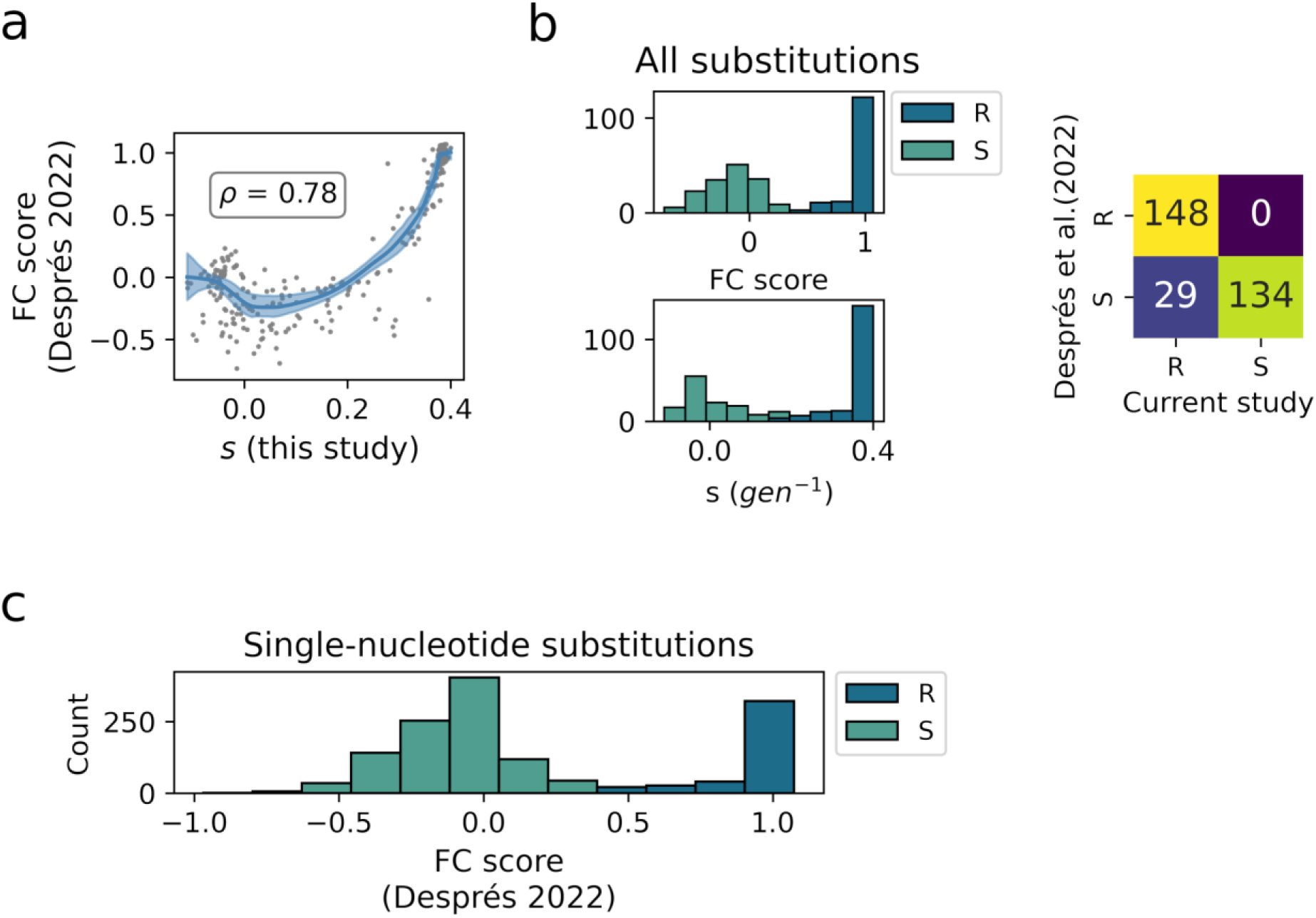
Single-nucleotide substitutions in the *FCY1* coding sequence readily confer resistance to 5-FC, across experiments. **a,** Comparison of median selection coefficients for coding sequence mutants (n=960) added to the bulk competition experiment with previously published variant effect scores measured in the same conditions^36^. Values have been aggregated into medians for all genotypes which result in the same amino acid sequence, such that n=311 points are shown. A LOWESS fit highlights a mostly monotonically increasing trend, as shown by a high Spearman correlation (𝑝 < 10^−64^). **b,** The set of coding variants which confer resistance to 5-FC is consistent across the two experiments. All mutants which were added to the “+*FCY1*” samples (**Extended Data Fig. 3a**) were classified into resistant (R), and sensitive (S) variants according to their effect in 5-FC conditions, using Gaussian mixture models (*Methods*). This was done using either the FC score reported by^36^ (top) or the selection coefficients inferred in the current study (left). The confusion matrix (right) shows that, among all variants which were classified as resistant using data from^36^, none appear sensitive to 5-FC in the current study. As in the previous panel, mutated sequences were collapsed into amino acid substitutions resulting in 311 variants. **c,** Almost a third of single-nucleotide substitutions in *FCY1* confer resistance to 5-FC. The distribution of variant effect scores from^36^ for all single-nucleotide substitutions in *the coding sequence*. For each genotype (n=1411), the score reported for the corresponding amino acid variant was used. Only 20 of the possible nucleotide substitutions are missing, mostly in the first and last codons. Variants are identified as resistant (R) or susceptible (S) according to the classification shown in panel **b**. The “R” class includes ∼29% (408/1411) of the single substitutions shown.

**Extended Data Fig. 6:**
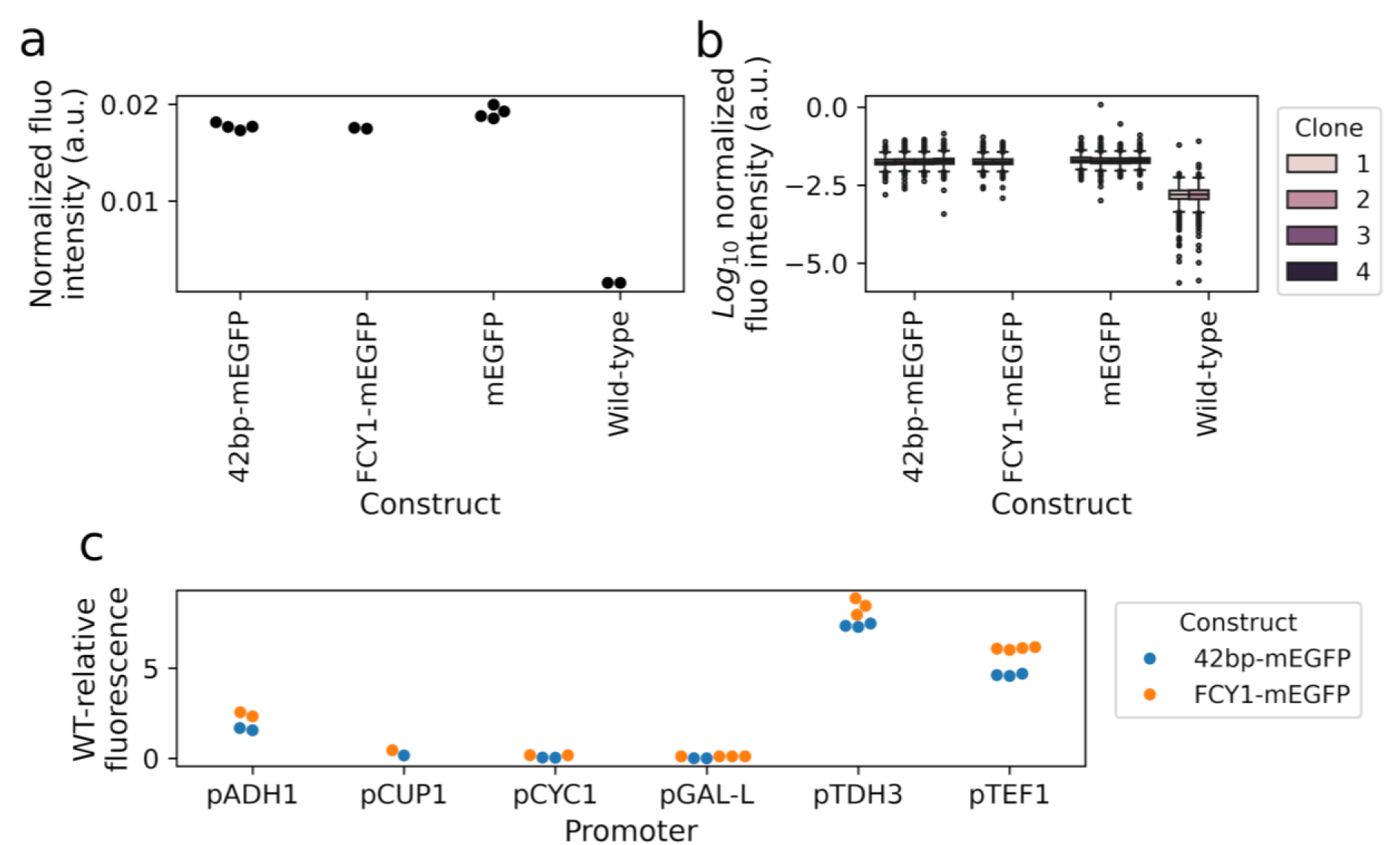
Validation of the 42bp-mEGFP construction. **a,** Medians of normalized fluorescence intensity, as measured by flow cytometry (*Methods*), for three mEGFP insertions at the genomic *FCY1* locus. Each point is a distinct clone of the corresponding strain, except for the WT. In the 42bp-mEGFP construct, which was used in the current study to measure promoter activity, the mEGFP green fluorescent protein is fused in C-terminal of the first 42 bp of the *FCY1* coding sequence. In contrast, FCY1-mEGFP designates a C-terminal fusion with full-length *FCY1,* while mEGFP stands for the insertion of the fluorescent protein in place of the *FCY1* coding sequence. **b,** As on the left, but showing the full distribution of log-scaled normalized fluorescence intensities for each clone (5000 events). **c,** Comparisons of median WT-relative fluorescence intensities for the different promoter constructs, in the 42bp-mEGFP or FCY1-mEGFP backgrounds. In each case, the WT-relative value has been computed relative to the corresponding reference (WT *FCY1* promoter sequence, upstream of either the 42bp-mEGFP of FCY1-mEGFP construct). The ranks of the promoters are unchanged between the two backgrounds, although WT-relative fluorescence intensities are consistently higher when the fluorescent protein is fused to full-length *FCY1*. Values shown for the 42bp-mEGFP are the same ones which were used throughout the paper to draw expression-fitness curves, although replicate measurements obtained from the same clone were aggregated into grand medians. The FCY1-mEGFP strains had not been validated prior to cytometry. All clones which displayed a fluorescence intensity clearly different from the corresponding WT are shown. Because the promoter cassettes were directly inserted in place of the endogenous promoter in the WT background, the WT expression level (and thus fluorescence intensity) is expected in cases where integration failed (*Methods*).

**Extended Data Fig. 7:**
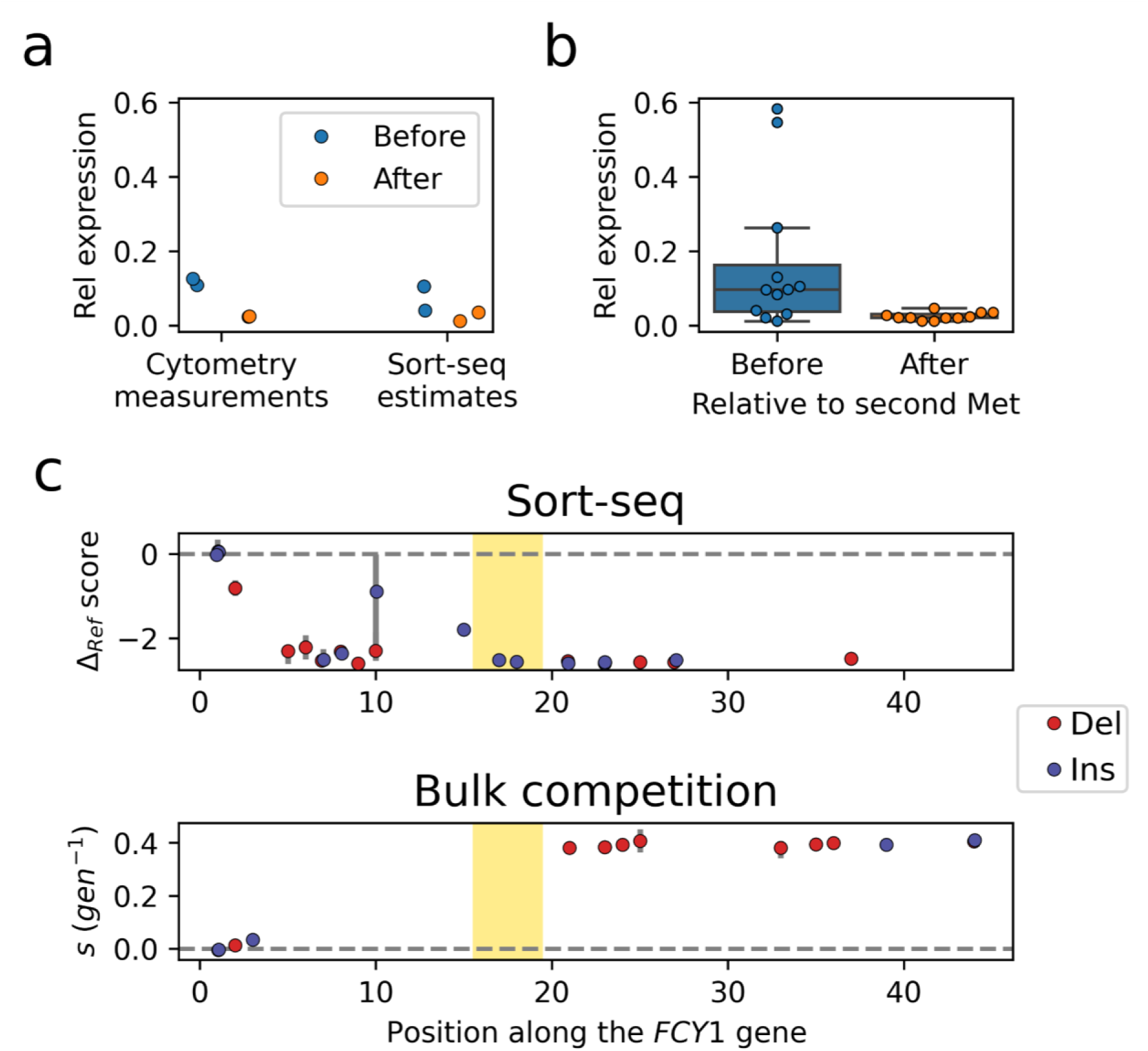
Indels before and after the second methionine codon of *FCY1* have contrasting effects on expression level. **a,** Comparison of relative expression levels for the four indel mutations which were included among the set of validation mutations, classified according to whether they occur before or after the second methionine codon of *FCY1*. Cytometry measurements from reconstructed mutants as well as estimates obtained from the large-scale sort-seq experiment are shown. **b,** Relative expression levels, as estimated from the sort-seq experiment, for all detected indel mutations in the *FCY1* coding sequence. Those occurring after the second methionine are clearly more impactful. **c,** Median effect of insertion and deletion variants detected in the sort-seq (top) and bulk competition (bottom) experiments. The shaded area encompasses the three bases of the second methionine codon. Mutations occurring before this codon were recovered at a much lower frequency from the bulk competition, which is consistent with the observation that they do not completely abolish *FCY1* expression and thus do not confer resistance.

**Extended Data Fig. 8:**
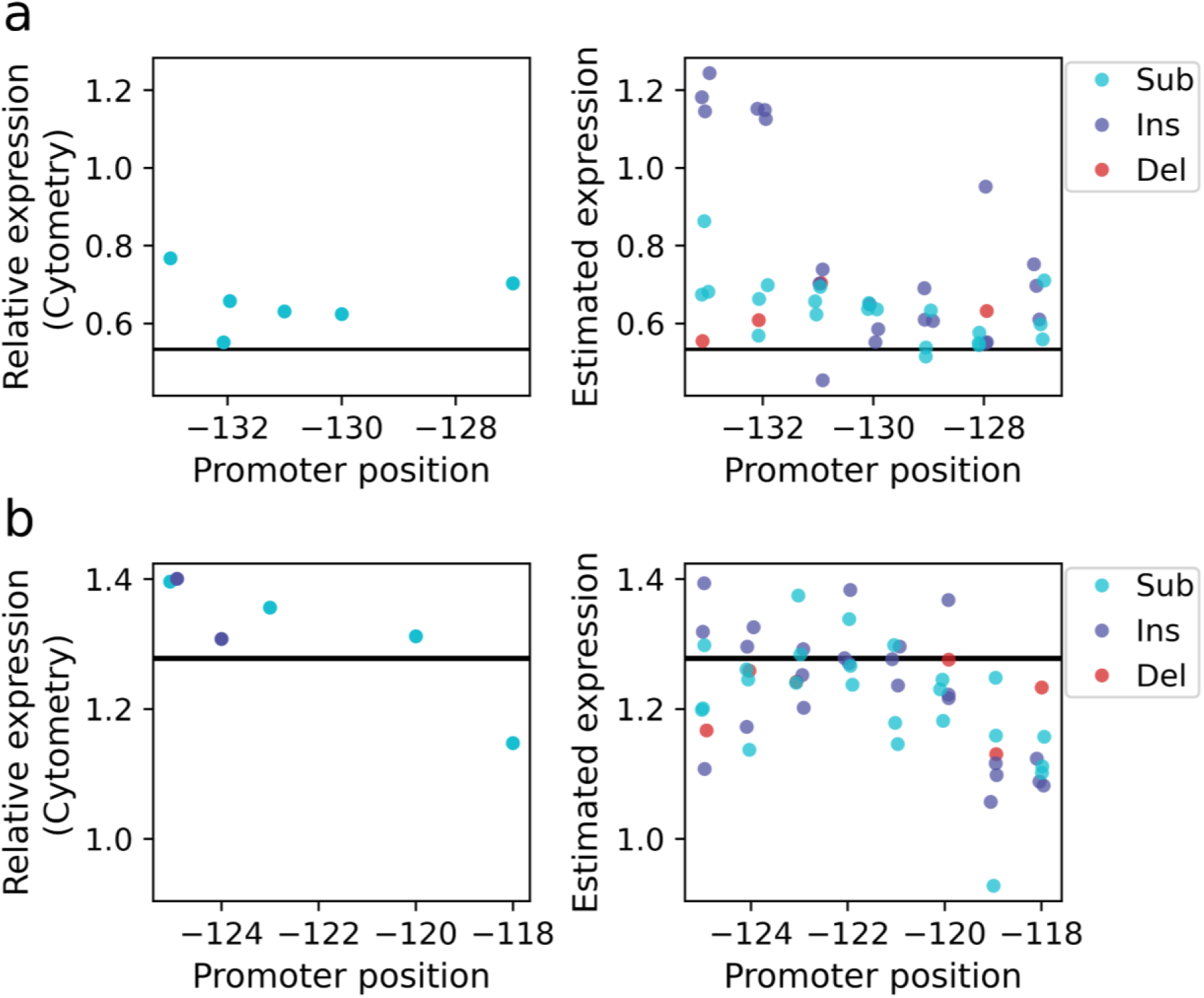
Deletion of putative TF binding sites is not more impactful than single-nucleotide mutations in the same regions. **a,** Deletion of positions −133 to −127 of the *FCY1* promoter, corresponding to site 1 (positions −132 to −127) of Fig. 4e. The WT-relative fluorescence of the deletion mutant, from three replicate measurements in flow cytometry, is shown as an horizontal line whose width displays the minimum and maximum values among the three corresponding medians. Among reconstructed validation mutants (left), one single-nucleotide substitution decreases *FCY1* expression as much as the deletion, as shown by its median WT-relative fluorescence intensities (dots; grand median of three replicate cytometry measurements). Among all mutations assayed by sort-seq (right), the most impactful single-nucleotide changes decrease expression as much as the deletion, as shown by the corresponding relative expression level estimates (dots; median of the 95% prediction interval obtained from sort-seq calibration). b, Deletion of positions −125 to −118, corresponding to site 2 (−125 to −120) of Fig. 4e. The plots are the same as in a, but this time for a deletion which increases *FCY1* expression. Cytometry measurements of validation mutants (left) and sort-seq estimates (right) both show that many single-nucleotide changes have an effect that is comparable to that of the deletion – or even larger.

**Extended Data Fig. 9:**
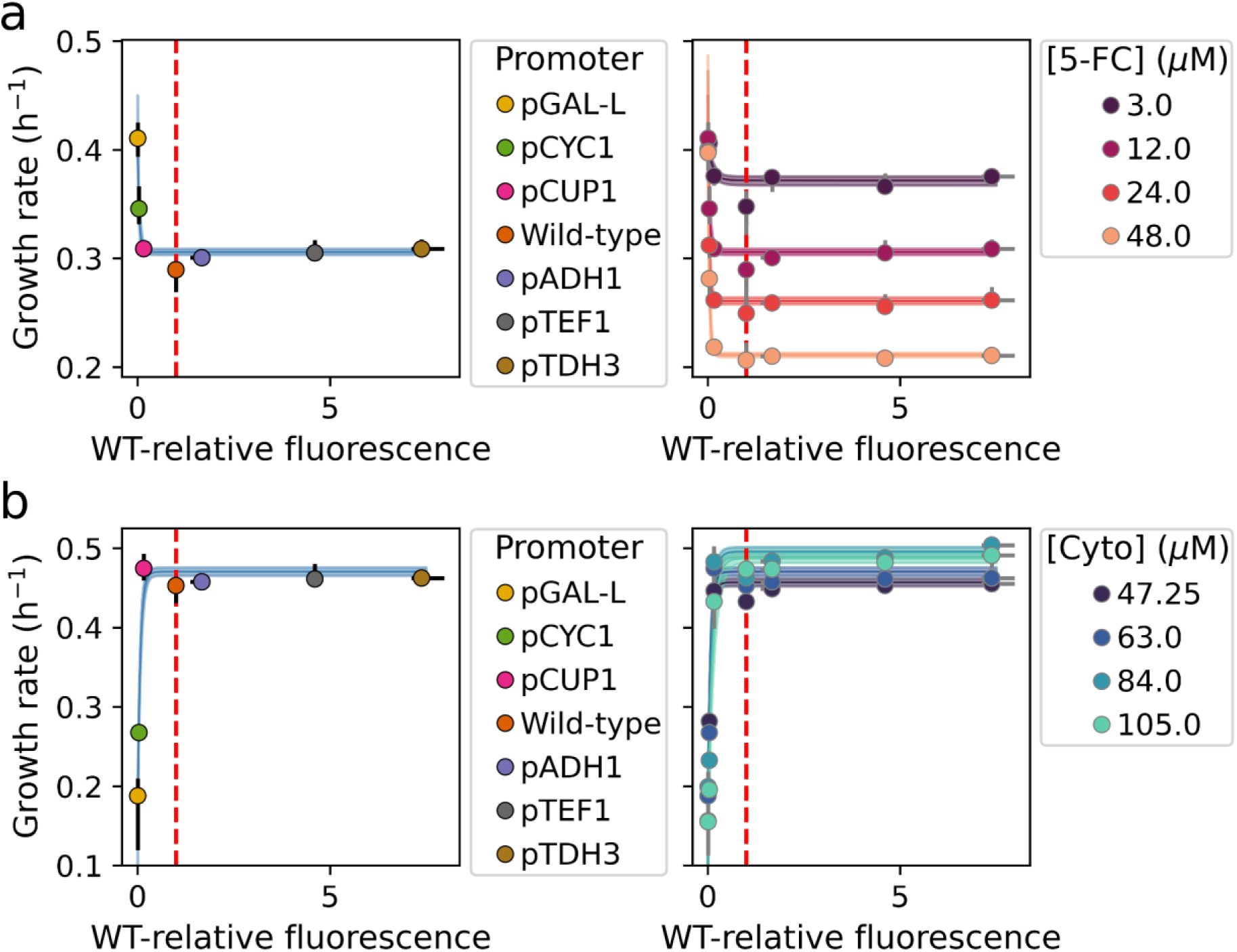
The insensitivity of fitness to changes in *FCY1* expression level is constant across many concentrations of 5-FC and cytosine. **a,** Fitness functions in 5-FC. In [12 μM] 5-FC (left), a constant plateau of low fitness is reached starting from ∼0.18 WT-relative expression (same as Fig. 5b). Across three additional concentrations of 5-FC (right), the boundary of the plateau does not change. Its height however varies, such that lower growth rates are observed for increasing 5-FC concentrations. b, Fitness functions in media lacking uracil, where deamination of cytosine by Fcy1p is essential. When cytosine is added at [63 μM] (left), a similar plateau is observed from ∼0.18 WT-relative expression, such that any further increase in *FCY1* expression does not increase growth rate. When three additional concentrations of cytosine are tested (right), the boundary of the fitness plateau also remains constant. Error bars around each dot display minima and maxima of three replicate measurements for growth rate and relative fluorescence intensity for each combination of promoter and 5-FC/cytosine concentration. In addition, dashed vertical lines identify the WT expression level (relative fluorescence intensity of 1.0).

**Extended Data Fig. 10:**
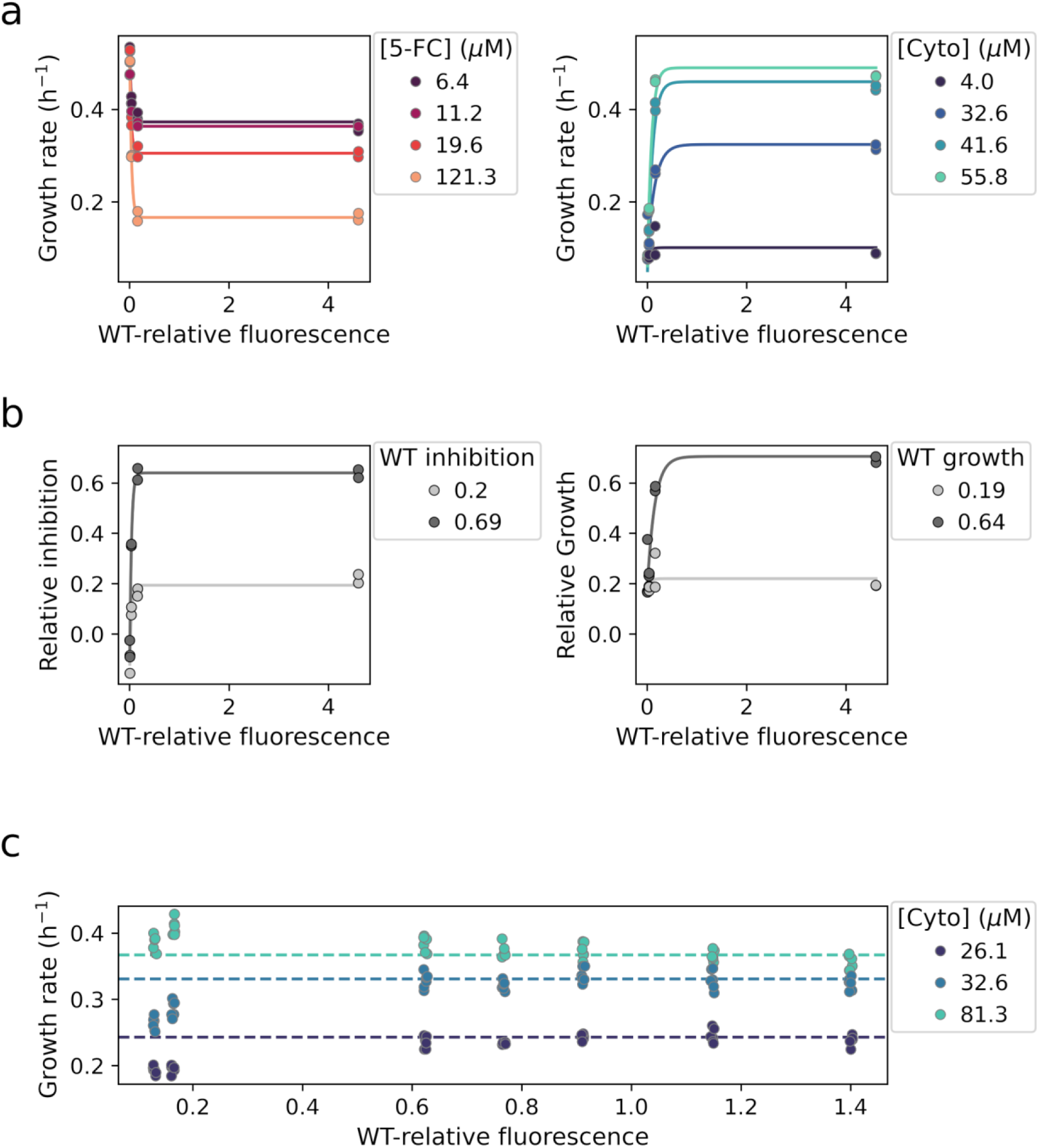
The expression-fitness function of *FCY1* displays a sign inversion between 5-FC and cytosine conditions. **a,** Fitness functions across 5-FC and cytosine concentrations covering the full ranges of the corresponding dose-response curves (**Extended Data Fig. 2**). Exponential growth rate measurements were performed in duplicates for each of the promoter constructs which define the elbow of the fitness function: pGAL-L, pCYC1, pCUP1 and pTEF1. In 5-FC (left), the conditions shown, from the lowest to the highest concentration, respectively result in relative inhibition levels of 0.20, 0.29, 0.40 and 0.69. In cytosine (right), the concentrations shown are respectively associated with the following relative growth levels: 0.19, 0.64, 0.93 and 0.97. **b,** Comparison of fitness functions obtained in 5-FC (left) and cytosine (right) concentrations resulting in matching levels of relative inhibition or growth for the WT reference strain (**Extended Data Fig. 2**). Although growth does not reach 0 in cytosine conditions when *FCY1* expression is decreased to very low levels, the two pairs of curves display very similar shapes. Same data as in panel **a**, but converted into relative inhibition or growth values using the relevant dose-response curve. **c,** Increasing *FCY1* expression through single mutations in the promoter does not provide a benefit in cytosine conditions, where Fcy1 activity is essential for growth. Exponential growth rate measurements were performed in triplicates for eight reconstructed promoter mutants covering the full range of expression changes available from single-nucleotide changes (Fig. 5d). From the lowest to the highest, the three cytosine concentrations used respectively result in relative growth levels of 0.42, 0.64 and 0.97. The dashed horizontal lines display the medians of growth rates obtained for the two mutants closest to the WT expression level. Across all three concentrations, the mutant associated with the highest expression level is not clearly fitter than this baseline. This confirms that, just like they are not deleterious in 5-FC conditions, promoter mutations increasing *FCY1* expression are not beneficial in cytosine conditions.

