## Supplementary Material for "The shape of fitness functions and the distribution of mutational effect sizes jointly limit adaptation by regulatory mutations"

#### Supplementary Note:
#### Secondary mutations can bias selection coefficient estimates

The initial genotyping of mutant pools from the bulk competition experiment, using a short sequencing fragment of 174 bp (red primer pair; **Fig. 2a**), revealed replicability issues as well as biases in estimated $s$. Selection coefficients obtained for the same mutants only displayed a high correlation between culture replicates of the same transformation replicate (**Supplementary Fig. 1a**). Moreover, the distributions of selection coefficients were clearly biased towards deleterious effects and appeared to move towards negative values over the course of the experiment (**Supplementary Fig. 1b**). The time dependency of this shift suggested that the reference (WT promoter) was getting fitter over time.

To explain these patterns, we hypothesized that secondary resistance-conferring (thus strongly beneficial) mutations had been introduced during the construction of the mutant library. If such mutations occurred outside of the sequenced region, they would remain invisible to our genotyping and the corresponding reads would be assigned to any F3-F4 single mutant with which they co-occurred. Moreover, double mutant combinations would be shared within a transformation replicate. This would result in strong correlations among corresponding culture replicates, as in **Supplementary Fig. 1a**. In addition, all beneficial "hidden" mutations happening in an otherwise unmutated background would be combined with true WT reads to estimate reference fitness. Because such mutants would grow faster, their frequency within the WT group would increase with every generation, therefore gradually inflating the apparent fitness of the reference allele. This could potentially lead to almost all genotypes appearing less fit than the WT ($s < 0$) at the end of the experiment, as observed in **Supplementary Fig. 1b**.

We tested this hypothesis by reinserting the unmutated F3 and F4 sequences, employing the same approach used to construct the corresponding mutant libraries. The frequency of resistant clones was assayed by plating the resulting pools of reconstructed WT on media containing 5-FC. This experiment showed that F4 mutant pools likely contained $\sim$ 0.22 % of cells harboring secondary mutations conferring resistance to 5-FC, while this frequency was at least ten-fold lower for F3 pools (**Supplementary Fig. 3**). Accordingly, $\sim$ 0.12 % of cells would be resistant to 5-FC within each transformation replicate combining F3 and F4 mutants. Using simulations, we confirmed that introducing such a low frequency of "hidden" (occurring outside the sequenced region) resistance-conferring mutations in a pool of mostly neutral mutants is sufficient to replicate the patterns shown in **Supplementary Fig. 1 a,b** (**Supplementary Fig. 1c**). We however note that, in these simulations, the gradual shift of inferred selection coefficients towards negative values occurred in only three of four simulated transformation replicates. In addition, selection coefficient estimates display much less variance in simulations than in the bulk competition experiment. We nonetheless performed additional simulation to investigate how the frequency of secondary mutations and their fitness effect impact both the general bias of estimated selection coefficients as well as the correlation between transformation replicates (**Supplementary Fig. 4**). This showed that a higher frequency of mildly beneficial secondary mutations may be sufficient for neutral variants to consistently appear slightly deleterious (**Supplementary Fig. 4a**). The magnitude of the bias and the fraction of experiments where it is observed also likely depends on the duration of the competition experiment (in generations of the reference strain) and on the initial frequency of the WT reference.

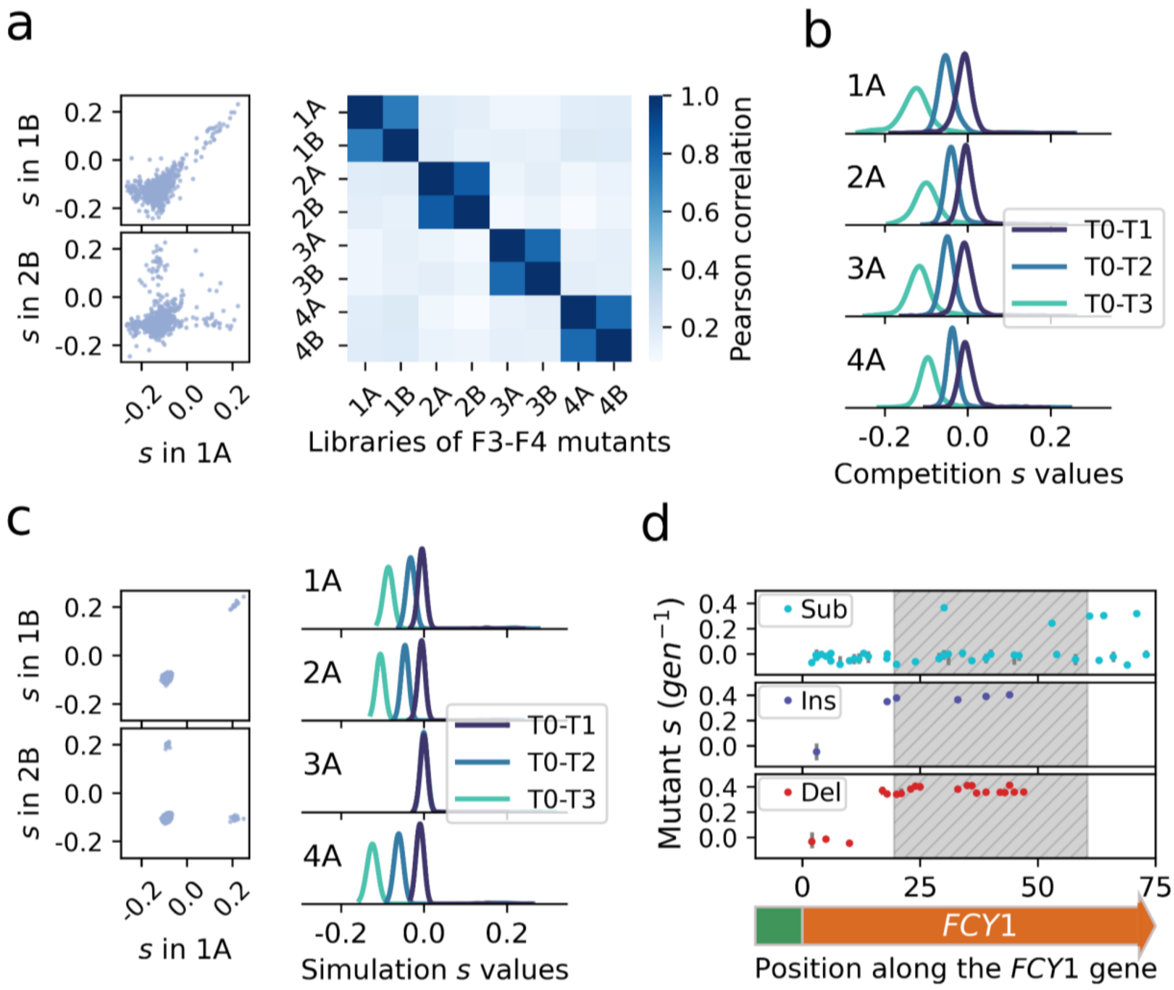


**Supplementary Fig. 1: Secondary resistance-conferring mutations in *FCY1* biased the estimated fitness effects of promoter mutations. a,** Selection coefficients inferred from the initial sequencing of bulk competition samples were only correlated within replicate transformations. Two representative scatterplots displaying all detected variants are shown (left), within the same transformation replicate (1A vs 1B) as well as between two transformation replicates (1A vs 2B). Pearson correlation coefficients from all pairwise comparisons (right) show that this phenomenon occurs across all samples (only displaying cultures without added *FCY1* mutations; **Extended Data Fig. 3a**). **b,** Following the initial genotyping of competition samples, the resulting distributions of selection coefficients display a time-dependent shift towards negative values. One of two culture replicates is shown for each transformation replicate, again using cultures without added *FCY1* mutations. Longer time intervals include intermediate timepoints, when applicable. For T0-T3, the T1 and T2 frequencies are for instance also used in the inference of selection coefficients. **c,** Simulations with hidden resistance-conferring mutations replicate experimental patterns. Equivalent plots to **a** and **b** are shown for a simulated bulk competition assay with 0.12% of resistance-conferring mutations (s=0.4) which are not detected during genotyping, using the same experimental design (8 cultures from 4 transformations). **d,** Resequencing of bulk competition libraries using a longer fragment (**Fig. 2a**) reveals many strongly beneficial single indels in the *FCY1* gene, outside of the original genotyping fragment. The dashed area indicates the part of the CRISPR homology region which was not initially sequenced, where many resistance-conferring mutations are found. More single mutants are shown than in Extended Data Fig. 6, because biased samples, which were discarded from the final fitness estimates used in the main text, are included.

Secondary mutations may be especially likely to occur in the first 60 bp of the *FCY1* gene, which were used as a homology arm for the insertion of the mutated F4 fragments. Notably, single-nucleotide indels in this region could result in a LOF of *FCY1* and grant resistance (**Extended Data Fig. 1**). Such rare resistance-conferring mutations were observed in the competition experiment, even though the sequenced region only covered a third of the homology arm (**Supplementary Fig. 5a**). To assess whether all 5-FC resistance arose similarly, we randomly selected 16 5-FC resistant colonies (2 F3 and 14 F4) among reconstructed WT for Sanger sequencing of *FCY1*. For 12 of the 14 F4 clones, a single indel in the coding sequence was identified. In half of those cases, the mutation occurred outside of the short sequencing region initially chosen for the competition experiment (**Supplementary Fig. 5 b,c**). No putative resistance-conferring mutation was detected for the F3 clones. Yet, since most unwanted resistance seemed to arise from LOFs in the *FCY1* gene, we resequenced the samples from pooled competitions, amplifying a longer fragment (333 bp) including the first 43 codons of *FCY1* (black primer pair; **Fig 2A**). This resequencing revealed several more indels granting resistance, most of which were within the last 40 bp of the homology region (**Supplementary Fig. 1d**).

Correctly genotyping *FCY1* variants by sequencing a larger region alleviated the issues shown in **Supplementary Fig. 1 a,b**, but some artifacts still remained. In half of the libraries, the estimated selection coefficients still displayed a clear bias towards negative values at final timepoint T3, with a median well below 0 (**Supplementary Fig. 6a**). The examination of log-transformed relative frequencies through time reveals that this is associated with a sudden decrease of most F3-F4 mutant genotypes relative to the WT between T2 and T3 (**Supplementary Fig. 6b**). This would be consistent with an initially small number of resistant mutants being included within the WT and progressively increasing in frequency, as described above. Additional simulations show that the low frequency (~ 0.03% or less) of 5-FC resistance which could not be attributed to *FCY1* mutations in WT reconstruction experiments is consistent with the persisting bias in selection coefficients (**Supplementary Fig. 9**). Four biased samples (among eight, like in the experiment) were observed in two thirds of replicate simulations using an initial frequency of 0.03% for 5-FC resistance. As such, the bias which persists in the bulk competition data even after resequencing is plausible under this proposed mechanism. We accordingly restricted all subsequent analyses to the T0-T2 data of the four unbiased samples (see **Fig. 3**).

Overall, experiments show that secondary beneficial mutations were introduced at a low but substantial frequency during the construction of our mutant libraries, while simulations confirm that this mechanism is sufficient to bias estimated selection coefficients. Such issues may affect other large-scale genome editing studies, owing to the high error rate of synthetic oligonucleotides[1]. In the current work, the effect of secondary mutations was particularly pronounced – and thus easier to detect –, since most single-nucleotide indels in the *FCY1* gene are strongly beneficial ($s \approx 0.4$). In a system where the most impactful secondary mutations would instead have been mildly beneficial, the effect might have been more subtle and thus less obvious, while nonetheless biasing all estimated selection coefficients towards negative values (see **Supplementary Fig. 4a**).

#### Supplementary Methods: Fluorescence-activated cell sorting (FACS)

As described in the *Methods* section, a FACSMelody (BD Biosciences) cell sorter was used to sort three mutant populations (transformation replicates), each in two (technical) replicates. Yeast cells were sorted according to their mEGFP fluorescence intensity, following a stringent gating procedure to select only single cells. The same three gates were used in each replicate experiment. The first one was a hexagonal gate defined over the FSC-A and SSC-A dimensions, using the following vertices: {(67811.63608618447, 30234.74780125552), (8834.822727312474, 8971.122270922499), (3879.279140855805, 1870.6636705422352), (8834.822727312474, 1039.145983024778), (100349.91030713769, 4260.322939281529), (167029.65512181344, 25847.660692198682)}. Across three pre-sorting samples of 20 000 events for which fluorescence data was saved, between 98.32% and 99.16% of all events were kept at this step. The second gate was rectangular over the SSC-H and SSC-W dimensions, defined by vertices {(1614.020866364354, 68046.53452600344), (13626.118022071332, 68046.53452600344), (13626.118022071332, 61205.63172555296), (1614.020866364354, 61205.63172555296)}. Across the three pre-sorting samples, between 22.11% and 33.99% of the initial events remained after this selection step. The final gate was also rectangular, but over dimensions FSC-H and FSC-W, and defined by vertices {(7364.164637190676, 73659.80731834323), (39866.54914769986, 73659.80731834323), (39866.54914769986, 60549.9472347268), (7364.164637190676, 60549.9472347268)}. Between 21.92% and 33.84% of events across the three pre-sorting samples passed all three gates. This process is shown graphically for one of the three samples (**Supplementary Fig. 11a**).

From yeast cells which passed the three gating steps, samples corresponding to five bins of fluorescence intensity were collected. Between 10 000 and 50 000 cells were collected per sample, as described in the *Methods* section. The five bins were defined graphically using the FACSChorus software of the FACSMelody and reused with the exact same fluorescence intensity thresholds across all sorting experiments. The corresponding thresholds for the five bins were: {0: (min=116.14564007554117, max=692.3990240619734); 1:(min=1093.6867105032413, max=2101.4995368500568); 2:(min=2091.0116239206377, max=3398.1987171748783); 3:(min=3647.1553305311645, max=5973.265614400292); 4:(min=6209.887318981933, 10973.78499702543)}. These five bins fully covered the distribution of fluorescence intensities observed after gating, as shown using one of the pre-sorting samples (**Supplementary Fig. 11b**). Within each resulting sample, the frequencies of single mutants was assessed by sequencing, as described in the *Methods* section.

### Supplementary Figures

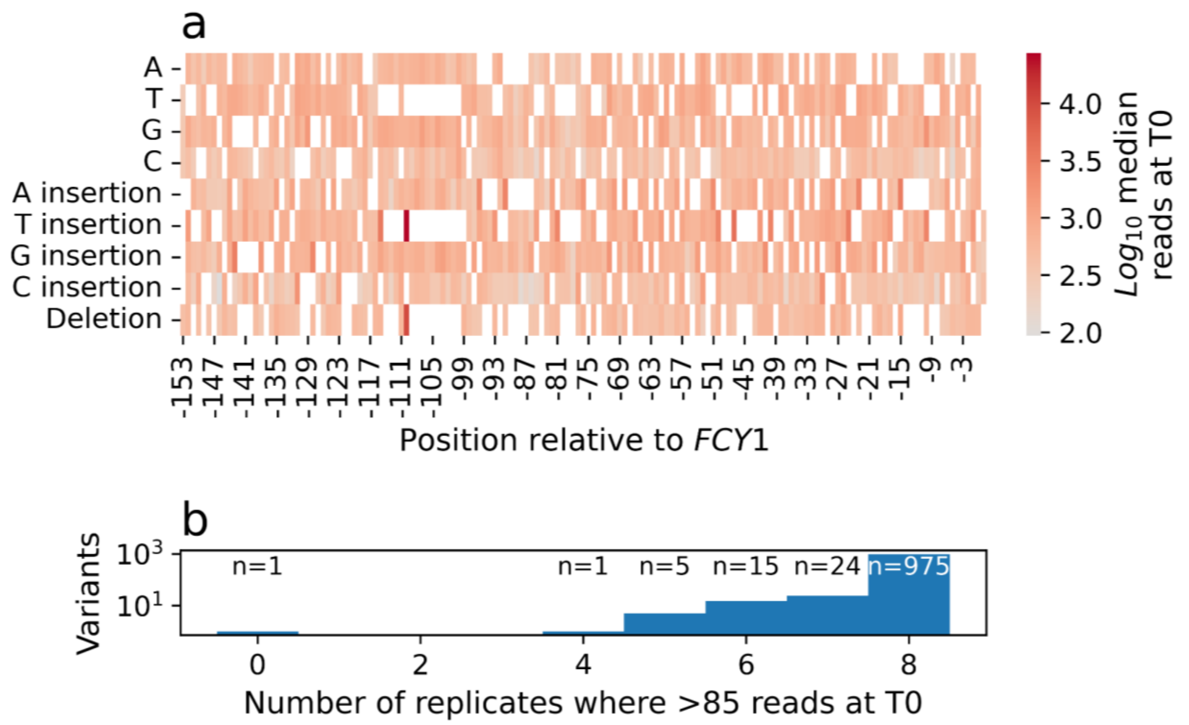


**Supplementary Fig. 2: More than 95% of single mutations in the *FCY1* promoter are detected in all replicate mutant libraries. a,** Coverage of all possible single-nucleotide changes across the 154 bp of the F3-F4 promoter region. The median number of reads from T0 samples of eight replicate bulk competition experiments (F3-F4 pools without added *FCY1* variants) is shown. For substitutions (A, T, G, C; top), white tiles denote WT nucleotides, and thus impossible mutations. For insertions and deletions, almost all white tiles appear due to repeated bases. From TTTT, any single deletion for instance results in the same TTT sequence, while any insertion of a T nucleotide generates TTTTT. As such, only one variant exists in both cases. Among the many white squares, only one indicates a missing single mutant: the insertion of A at position -152. The +1 shift in the position of insertions (-152 to 1, instead of -153 to 0 for other types) is due to how the insertion mutant pools were ordered. The position of a single insertion was initially defined as the base immediately upstream of it, rather than its index in the alignment. **b,** Number of replicate T0 samples (n=8; same as **a**) where each F3-F4 variant exceeded the abundance threshold. More than 95% (975/1021) had sufficient read depth across all bulk competition experiments.

.

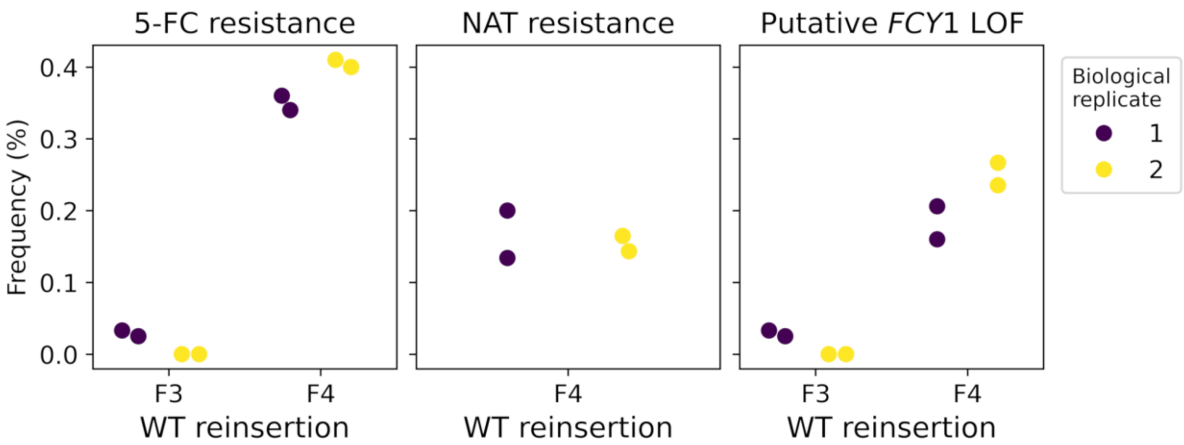


**Supplementary Fig. 3: Reintegrating wild-type F3 and F4 sequences in the *FCY1* promoter induces a low frequency of 5-FC resistance.** The native promoter was reconstructed from either F3 or F4, amplified from WT genomic DNA. The same approach previously employed to insert pools of mutated fragments was used (*Methods*). After recovery of transformants by the scraping of Petri dishes, the resulting cell suspension was diluted and plated on solid media containing [100 µg/ml] 5-FC. Resistant colonies were then counted and corresponding frequencies were calculated from cell concentrations estimated from optical density measurements. The CRISPR-Cas9-based insertion of F3 and F4 fragments (mutated or not) involves the preliminary replacement of the region with a NAT cassette (*Methods*). In the case of F4, this confers resistance to 5-FC. All colonies from F4 reinsertions were thus replicated on media containing nourseothricin (NAT) and the frequency of NAT resistance was subtracted from 5-FC resistance to obtain the corresponding frequency of putative *FCY1* LOF. Across all three plots, the points represent the same set of platings. Biological replicates denote two transformation replicates, while the two points within each indicate two dilution levels that resulted in countable colonies (technical replicates).

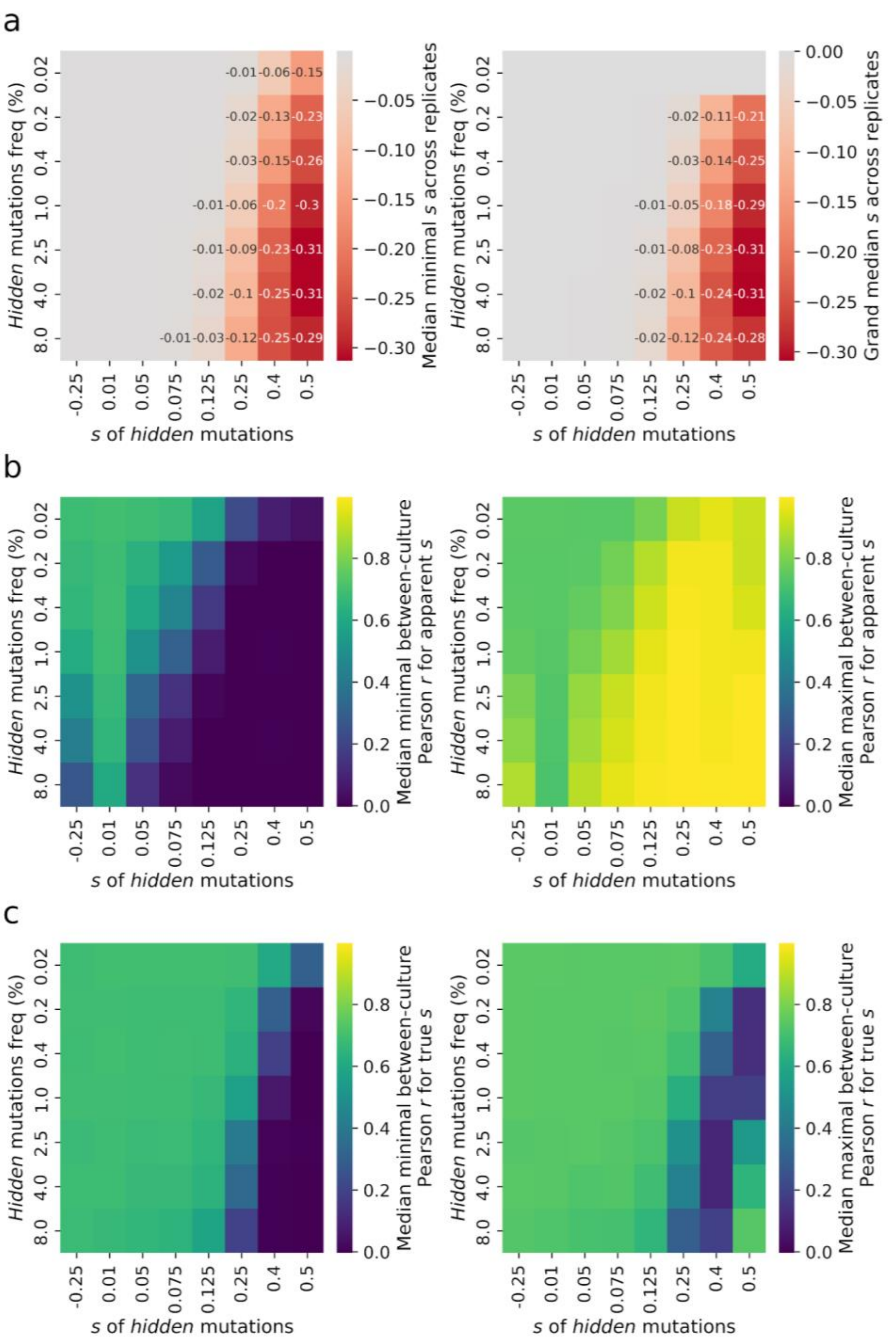


**Supplementary Fig. 4: Simulated bulk competition assays show that mildly beneficial mutations are sufficient to bias selection coefficient estimates.** For each combination of parameters, the specified frequency of invisible (not genotyped through sequencing) beneficial mutations, with the indicated *s*, is introduced within a pool of neutral variants (*Methods*). The experimental design from our experiment is replicated (eight cultures from four replicate transformations, all grown for approximately 20 population doublings). Across all heatmaps, each tile displays the median (or grand median) of six replicate simulations. Different summary statistics are shown for the same set of simulations. **a,** The median inferred *s* decreases from the expected value of 0 with increasing initial frequency (y axis) and *s* of invisible mutations (x axis). On the left, grand medians are computed from the most biased (lowest median *s*) culture within each replicate simulation. On the right, grand medians are instead obtained from all cultures within each replicate. All inferred selection coefficients have been calculated over the full duration of simulated experiments (T0-T3). This shows that a higher frequency (1.0% to 2.5%) of mildly beneficial secondary mutations (s=0.125) may be sufficient to reliably shift by -0.01 the selection coefficients estimated for mutants of interest. These simulations also reveal that deleterious secondary mutations are not sufficient to bias estimated selection coefficients. **b,** Between-culture correlations more strikingly vary across transformation and culture replicates when the frequency and selection coefficient of secondary mutations increase. On the left, the median Pearson correlation across the worst (least correlated) culture pair of each replicate simulation is shown. On the right, the same median is instead computed using the best (most correlated) culture pair in each simulated experiment. Large differences between the lowest (different transformation) and highest (same transformation) correlation coefficients coincide with substantial biases in the previous panel. **c,** Even if all beneficial secondary mutations are correctly genotyped, their presence introduces noise in selection coefficient estimates. Here, all reads associated with secondary mutations are removed prior to the computation of selection coefficients. Because only true reads of each variant of interest are now used, these values are referred to as "true *s*". Correlations between the worst (left) and best (right) culture pair of each simulation reveal that secondary mutations with s=0.4, as in our experiment, substantially decrease replicability across all assayed initial frequencies.

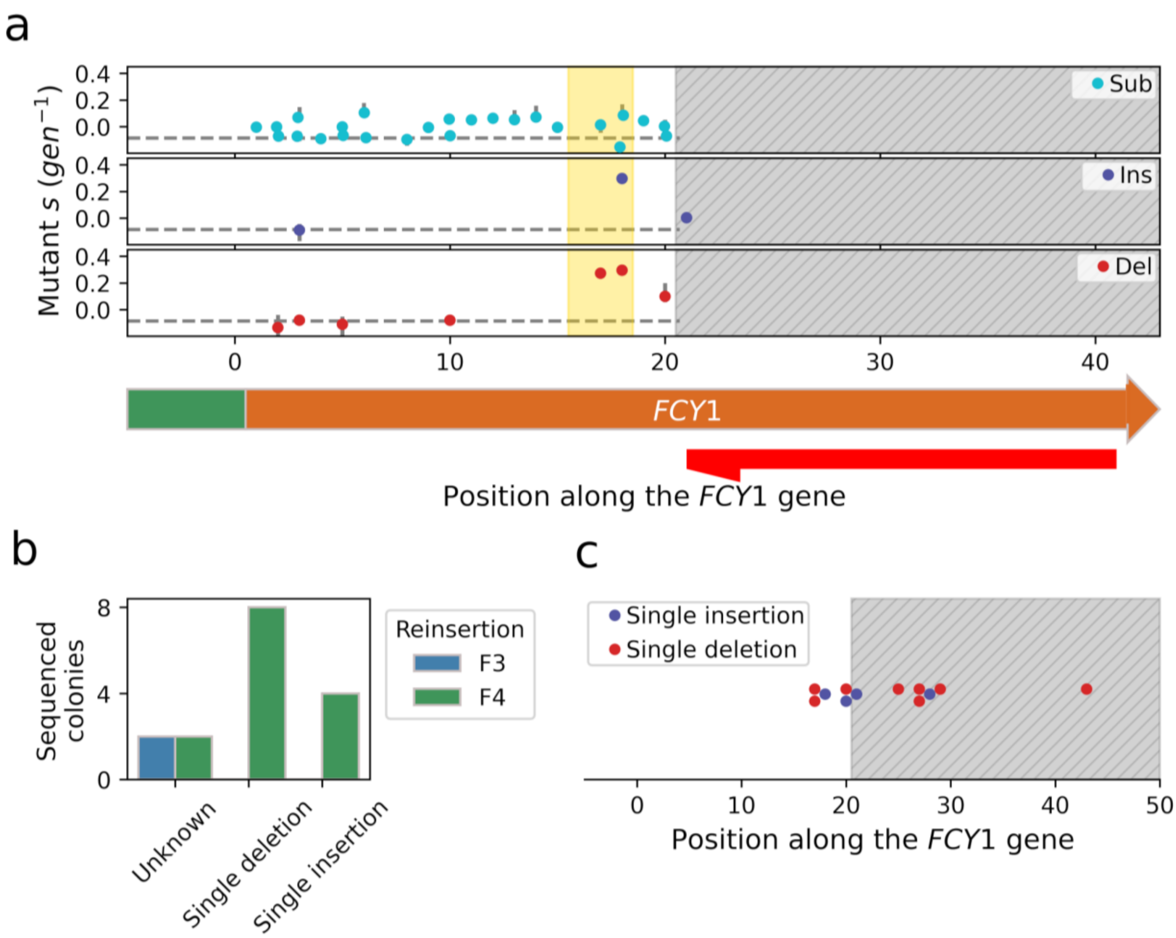


**Supplementary Fig. 5: Single indels were introduced in *FCY1* during mutant library construction.** **a,** A small number of beneficial indels were detected in *FCY1* following the initial genotyping of bulk competition samples (shorter sequencing fragment). All secondary single mutations identified in the first 20 bp of the *FCY1* coding sequence are shown. Points indicate the median selection coefficient across replicates where each mutant was found, while error bars display the corresponding minimum and maximum. The median selection coefficient across all variants in this assay is shown as horizontal dashed lines. Three resistance-conferring indels, with *s* much higher than this median, are visible within the second methionine codon of the gene (golden shaded area). The dashed area to the right indicates the region which could not be genotyped by sequencing, since it is covered by the oligonucleotide used in the first PCR step of sequencing library preparation (red). **b,** Most cases of 5-FC resistance in

CRISPR reconstructions of the wild-type F3-F4 promoter region are associated with a single indel in the *FCY1* gene. Among 16 resistant clones selected for Sanger sequencing (2 F3 and 14 F4), 12 harbor a single indel in *FCY1*. **c,** At least half of resistance-conferring indels occur outside of the genotyping region (initial sequencing, shorter fragment). All 12 indels from **b** are shown along the FCY1 coding sequence. Only six of them, including the insertion at position 21, could have been genotyped using the PCR primer shown in **a**, as indicated by the dashed area.

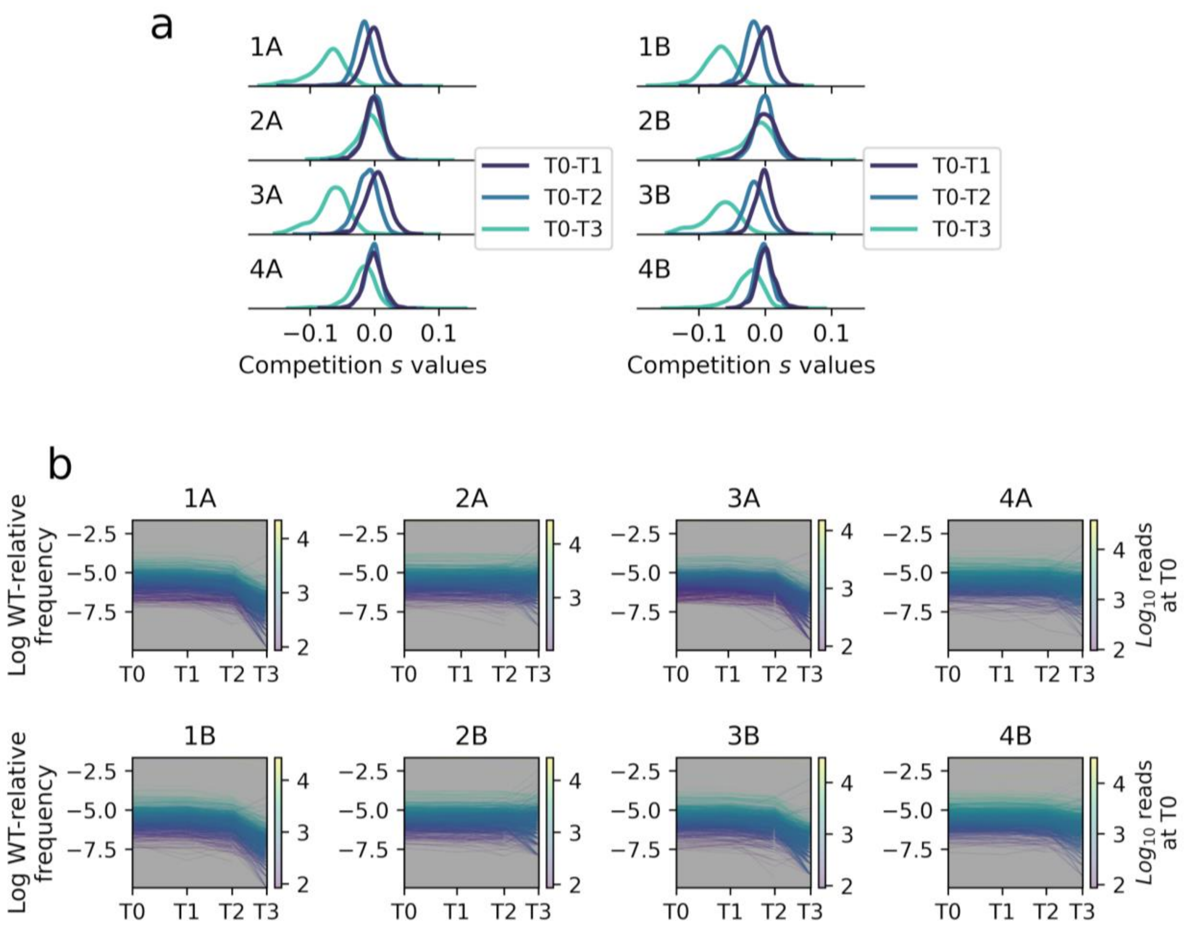


**Supplementary Fig. 6: Genotyping bulk competition samples using a longer sequencing fragment partly eliminates bias in selection coefficient estimates. a,** Half of the cultures display minimal bias when the longer sequencing fragment (**Fig 2a**, black) is used for genotyping. Across all eight cultures without added *FCY1* mutants, the distributions of *s* for F3-F4 variants are shown over each time interval, as estimated after resequencing. For two transformation replicates (culture pairs 1A/1B and 3A/3B), a substantial bias towards negative values persists. **b,** In cultures where selection coefficients are still biased after resequencing, the WT-relative frequencies of all F3-F4 variants suddenly decrease between sampling points T2 and T3. All time series of log-transformed WT-relative frequencies used to compute the *s* displayed in **a** are shown. Because all cultures are growing exponentially, each trajectory should be a line, with its slope set by the corresponding variant's selection coefficient. For cultures 2A/2B and 4A/4B, most trajectories remain linear for the whole duration of the experiment. For 1A/1B and 3A/3B, where *s* estimates remain biased, most slopes instead suddenly change between T2 and T3.

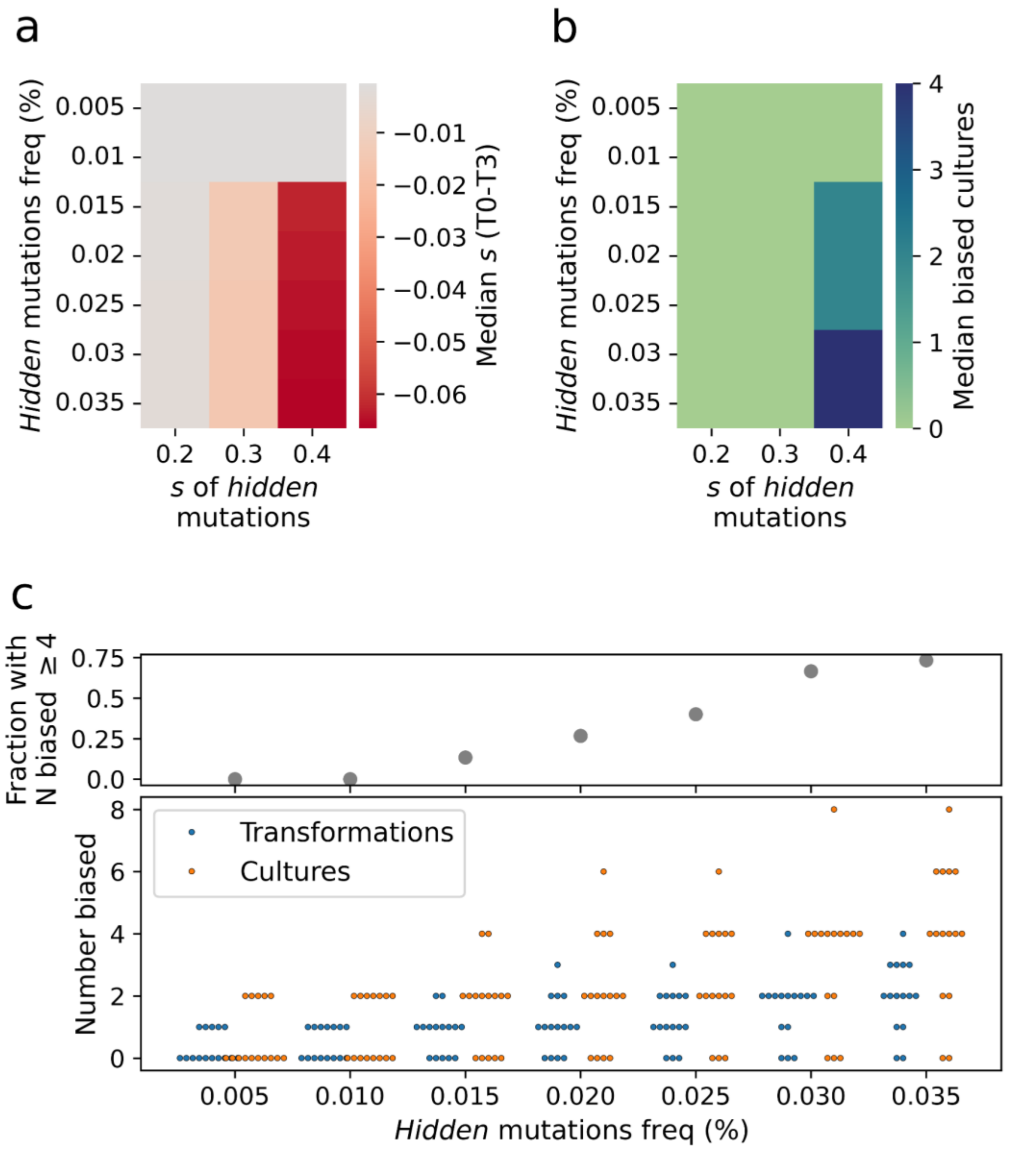


**Supplementary Fig. 7**: **The lower frequency of resistance-conferring secondary mutations could be consistent with the bias which remains after resequencing.** Additional simulations were performed using a low initial frequency of invisible resistance-conferring mutations, based on the frequency of 5-FC resistant colonies for reinsertions of region F3 (between 0 and 0.030%). For each combination of parameters, 15 replicate simulations were done. **a,** Lower initial frequencies of secondary mutants are sufficient to bias the median culture in a typical experiment, even if a lower *s* of 0.3 is used. Across all tiles, the grand median selection coefficient over the corresponding replicates – each made up of eight simulated cultures – is shown. **b,** Only fully resistant secondary mutants (s=0.4) are sufficient for at least one culture to display biased selection coefficients (median s below -0.05) in the median simulated experiment. **c,** When secondary mutants are fully resistant to 5-FC (s=0.4), even low initial frequencies are associated with a substantial probability of observing four biased cultures. The numbers of biased cultures (same definition as in **b**) and transformations (pair of cultures) for all replicate simulations of the s=0.4 column from the two previous heatmaps are shown. For an initial frequency of 0.030%, the probability that four cultures within an experiment display biased selection coefficients is of ~ 0.67 (10/15). While this starting frequency is the upper bound on resistance arising by non-*FCY1* mutations (as estimated from F3 reinsertions; **Supplementary Fig. 4**), this still shows that the level of bias observed in the experiment even after resequencing is plausible. Even lower initial frequencies of resistant mutants are associated with a substantial probability of observing at least four biased samples.

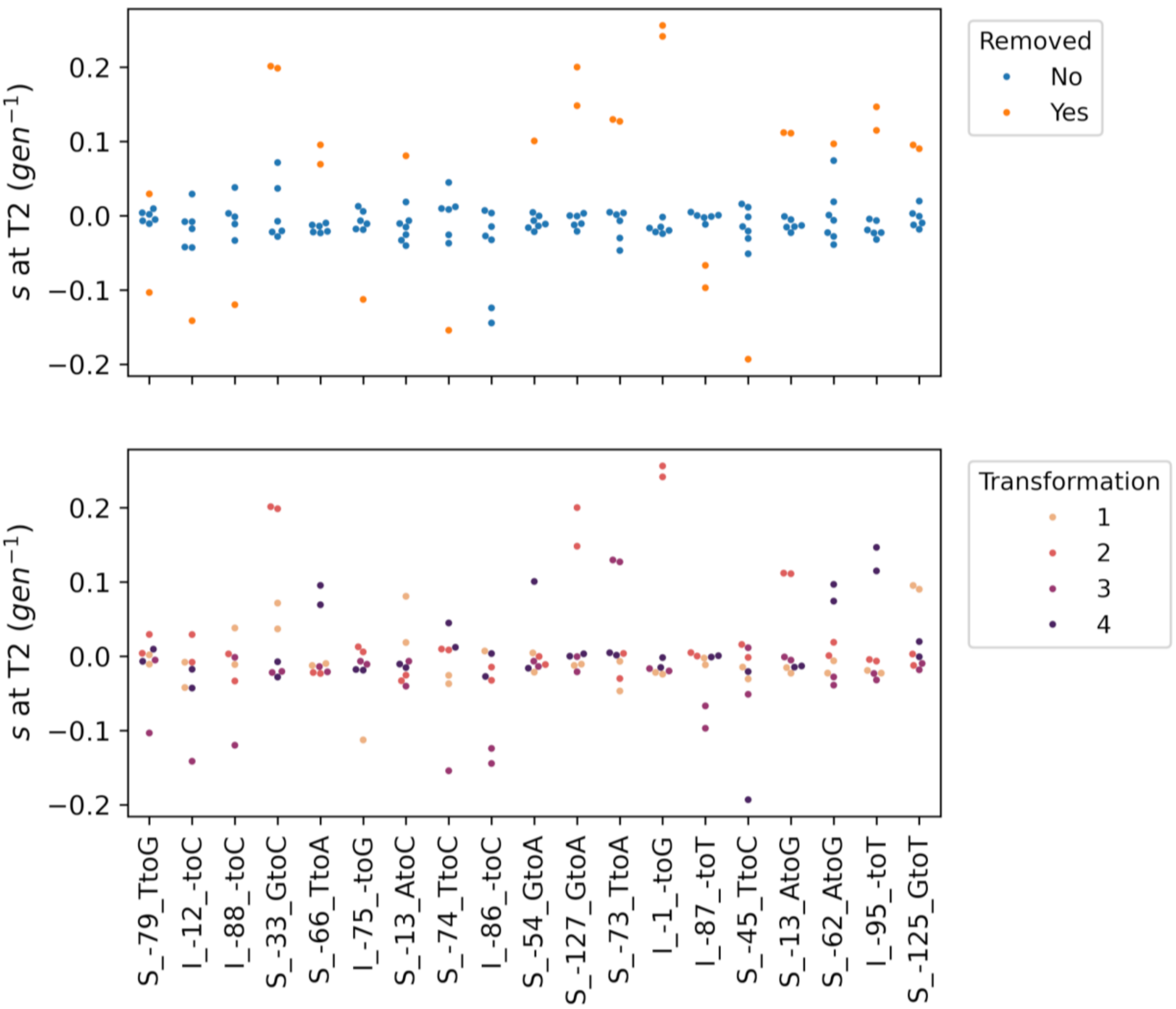


**Supplementary Fig. 8**: **The outlier removal approach mostly discards pairs of culture replicates displaying unusually high selection coefficients**. All high-variance genotypes selected for outlier filtering (*Methods*) during the analysis of the resequencing of bulk competition samples (cultures without added *FCY1* mutants) are shown. Corresponding identifiers (x axis) inform on the type (S=Substitution; I=Insertion) and position of each mutation. Replicate selection coefficients obtained from the T0-T2 of each replicate culture are shown (points). At the top, they are colored according to whether they are classified as outliers or not. While at the bottom, the corresponding transformation replicates are instead shown. Most identified outliers display higher (more beneficial) selection coefficients and these beneficial outliers are most often removed as pairs of cultures replicates derived from the same transformation. This procedure thus largely filters out anomalous *s* values which are consistent with undetected resistance-conferring mutations having been introduced in the corresponding genotype. Outlier filtering was performed before the selection of four unbiased cultures for further analysis, such that all eight replicates are shown here.

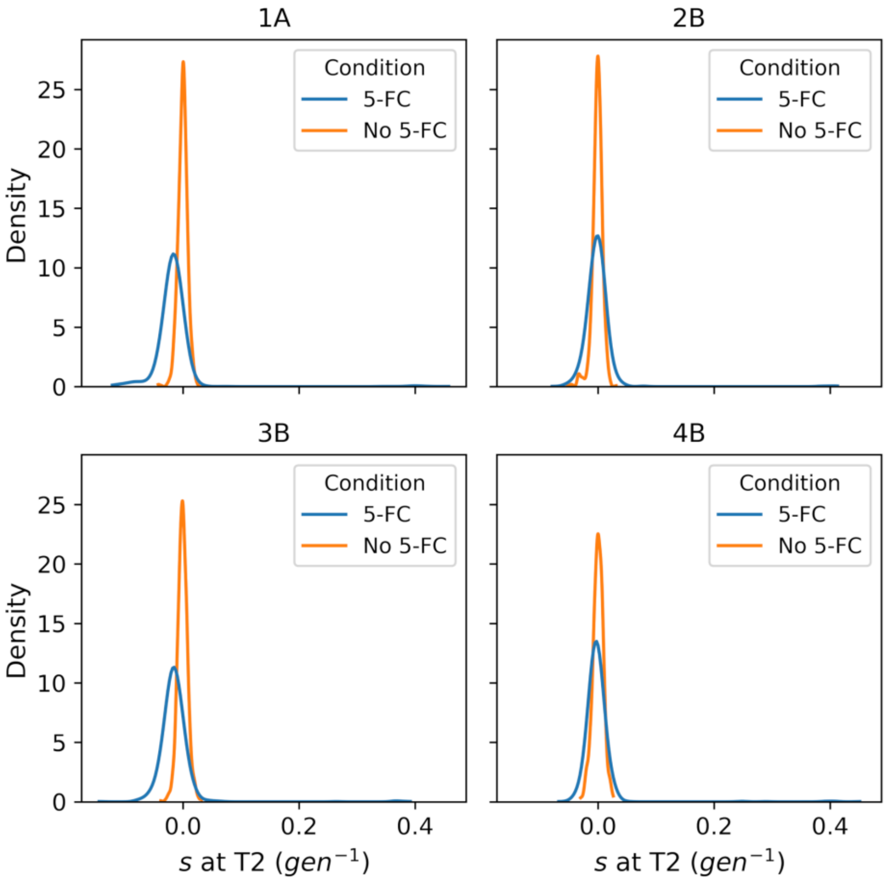


**Supplementary Fig. 9**: **The distributions of inferred selection coefficients display less variance for culture without 5-FC.** The four controls without 5-FC which were sequenced are compared with the corresponding cultures with 5-FC (resequencing; only cultures without added *FCY1* mutants). All identified single mutants, including secondary mutations introduced during mutant library construction, are shown. Tails of strongly beneficial effects, reaching s=0.4, are only seen in cultures with 5-FC. The dispersion around the distribution's mode is additionally higher with 5-FC than without. This confirms that the addition of 5-FC applied a selective pressure on the pool of mutants.

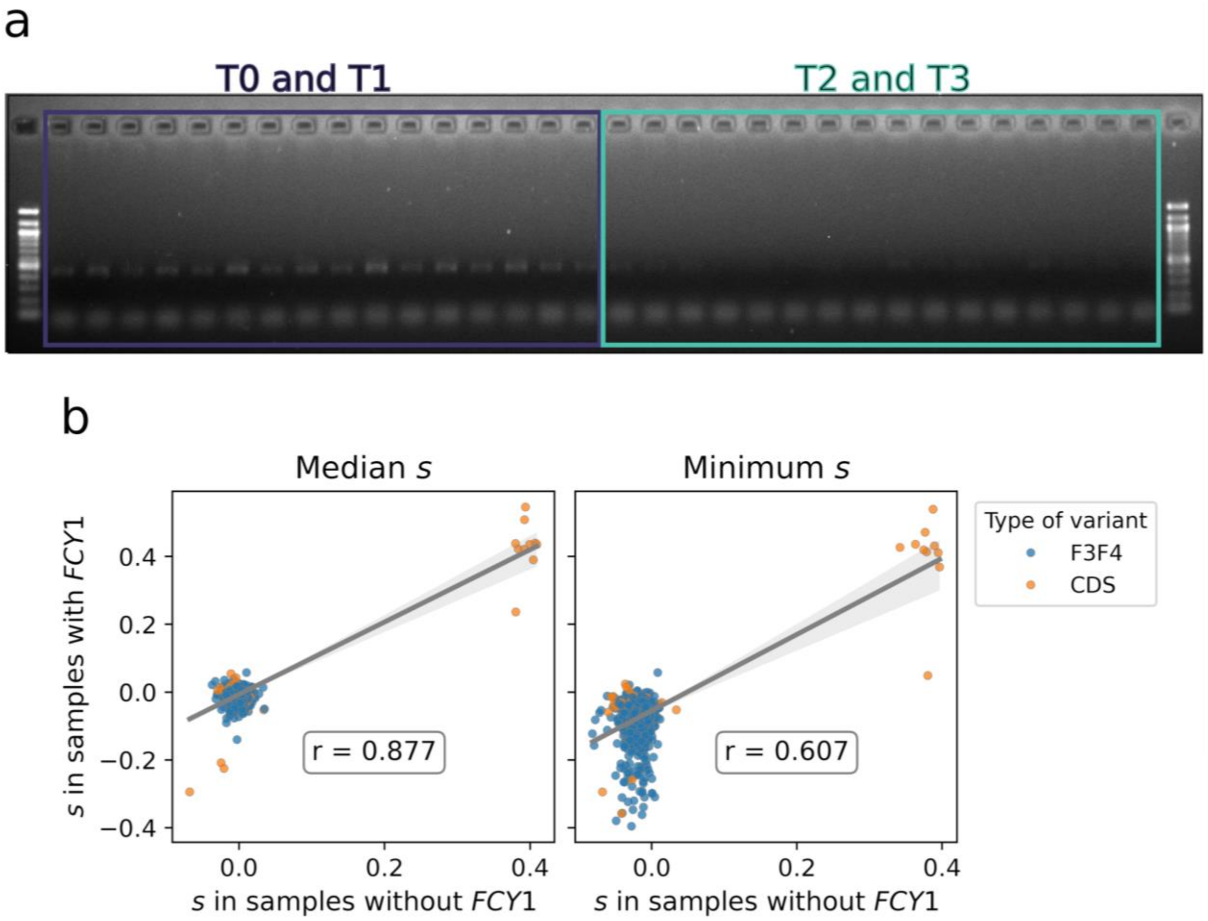


**Supplementary Fig. 10**: **Mixed cultures of promoter and coding sequence variants were quickly taken over by the *FCY1* mutants. a,** Gel electrophoresis image of the first PCR amplification step to prepare sequencing libraries of the F3-F4 promoter region (*Methods*), for the eight +*FCY1* cultures. Primers were designed to be specific to the pool of promoter mutants, to avoid amplifying the WT promoter sequence of the added *FCY1* variants. On the left, T0 and T1 PCR reactions have been loaded in alternance (from sample 1A to sample 4B), while the same was done on the right for T2 and T3 PCR reactions. Almost no band is visible for the T2 and T3 samples, although the same volume of PCR reaction was loaded in all cases. This suggests a much lower quantity of template DNA, and thus a smaller population size for promoter mutants. **b,** Comparisons of inferred selection coefficients in [12 μM] 5-FC for variants in the pool of promoter mutations, across samples with and without added coding sequence mutants. Variants which are identified as "CDS" are secondary mutations in the coding sequence which occurred during construction of the F3-F4 library (see **Supplementary Fig. 1**). Median selection coefficients display high agreement between the two types of samples (left). Using the minimal $s$ observed among replicate cultures (right) however reveals outlier negative values for samples with added *FCY1* mutants. This is consistent with the stochastic disappearance of some genotypes due to the decreased population size of F3-F4 promoter mutants.

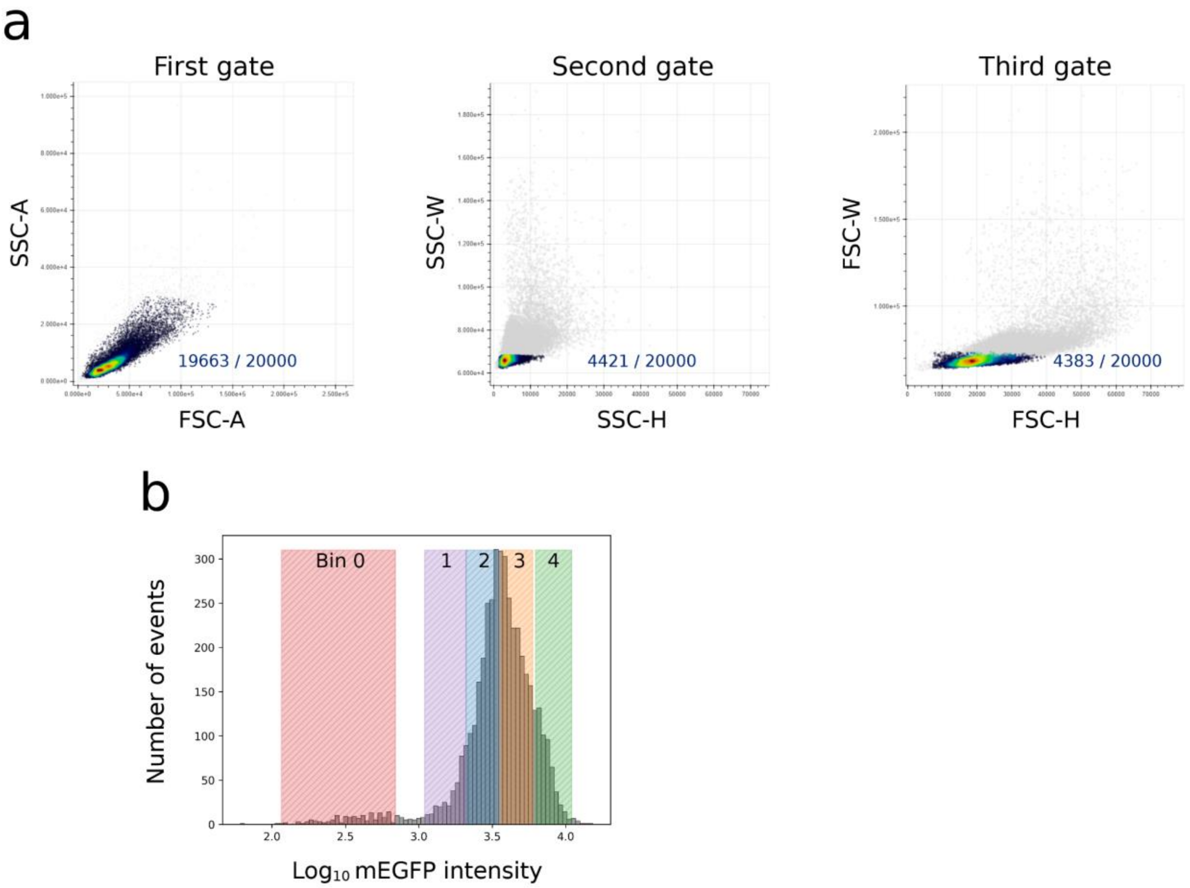


**Supplementary Fig. 11**: **Single yeast cells were sorted by fluorescence intensities in sort-seq experiments.** **a,** Gating strategy. Three successive gates were applied to select single cells of similar sizes. The first one was defined by six points over the FSC-A and SSC-A dimensions, while the two others were each defined by four points. For the second gate, this was over the SSC-H and SSC-W dimensions, while the third gate was instead defined over the FSC-H and FSC-W dimensions. The gating process is shown graphically for one of three samples of 20 000 events which were measured prior to cell sorting. Through each successive gate, the number of events selected for downstream sorting is gradually reduced. Events which are filtered out are shown as grey dots, while those which pass the gating steps are colored according to kernel density. Most of the filtering occurs at the second gate. **b,** Cell sorting strategy. Five bins were defined over the distribution of fluorescence intensities obtained after gating, to cover the full range of effects. Bin boundaries were defined in a preliminary experiment and kept constant for all replicate sort-seq experiments. The five bins, numbered 0 to 4, are shown over the histogram of fluorescence intensities obtained for the 4383 events which passed all three gates in **a**.

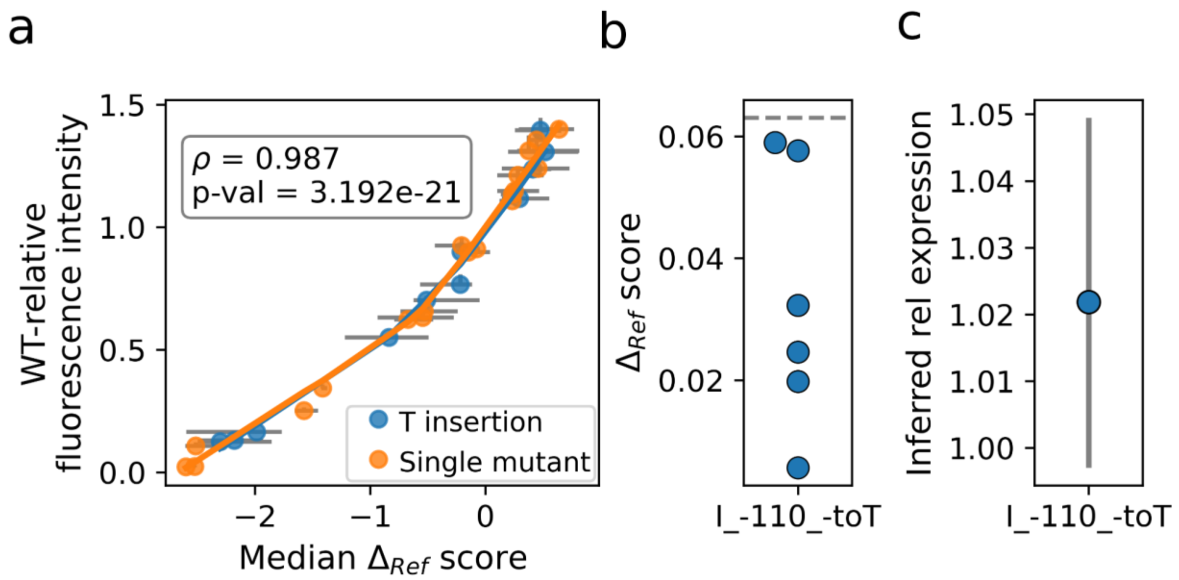


**Supplementary Fig. 12: More detailed look at the calibration curve of sort-seq scores. a,** LOWESS fit as in **Fig. 4c**, but performed separately for true single mutants and for the minority of validation mutants (11/30) which featured an additional T insertion within a poly-T region (positions -110 to -99) in the F3 promoter fragment. The two fits result in almost the same curve, thus showing that these unwanted double mutants do not have a systematic effect on the conversion of $\Delta_{Ref}$ scores into relative expression levels. **b,** Sort-seq scores observed across all six replicates for the corresponding insertion (I) of a single T at position -110. All values are below the median $\Delta_{Ref}$ observed across the six sort-seq experiments (dashed line). **c,** Relative expression level inferred from the sort-seq scores shown in **b** (**Fig. 4c**), with error bars displaying the corresponding 95% prediction interval. Panels **b** and **c** show that the insertion alone has a weak to nonexistent impact on *FCY1* expression, thus confirming the robustness of relative expression estimates to the (accidental) inclusion of double mutants among the validation set.

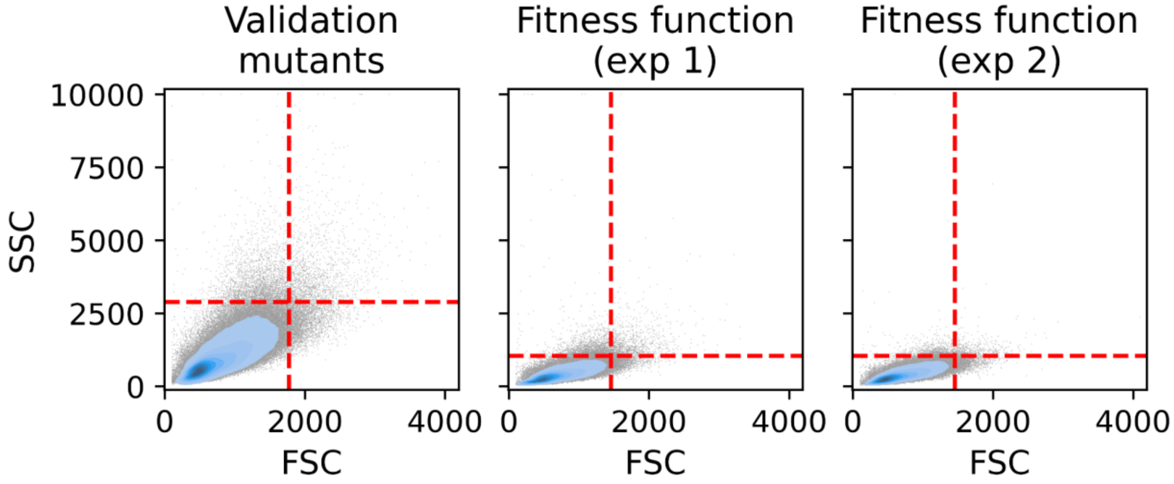


**Supplementary Fig. 13**: **Gating in flow cytometry experiments.** Each time flow cytometry was performed to measure mEGFP fluorescence in cultures of a single yeast strain, events were gated using the 99th percentiles of the forward scatter (FSC) and side scatter (SSC) distributions (dashed lines on each plot). This gating was performed separately within each experiment, as well as all comparisons with WT fluorescence intensity (to obtain WT-relative values). All fluorescence intensities were in addition normalized by the corresponding FSC value (*Methods*). Fluorescence data from the leftmost set of events is shown in **Fig. 4c** and **Fig. 5cd**. Data from the sets displayed in the middle and rightmost columns is instead shown in **Fig. 5bc** of the main text.
