## Supplementary material for "The shape of fitness functions and the distribution of mutational effect sizes jointly limit adaptation by regulatory mutations": Description of Supplementary Tables

Description of Supplementary Tables -- Aubé et al. (2025)

Table 1: Yeast strains used in this study.

Table 2: Oligonucleotides used in this study.
		 *Note* Oligo pair CLOP299-D3 / CLOP299-D4, used to amplify F4 opools for genomic insertion, confers a 60 bp homology with the FCY1 gene.
		 While this may seem incompatible with the insertion of F4 mutations in the 42bp-mEGFP background, the corresponding mutant libraries were
		 nonetheless successfully constructed. This was validated by a very small frequency of reads harboring the full 60 bp in the sequencing of
		 the sort-seq experiments (none of those problematic reads were used to estimate the expression level or promoter variants).

Table 3: Plasmids used in this study.

Table 4: Recipes for culture media used in this study.

Table 5: List of sequences (mutated F3 or F4) which were ordered as IDT oligo pools (oPools).
		 All sequences sharing the same "Pool name" were ordered as one oPool (one tube).

Table 6: Durations of passages during the bulk competition experiment.
		 For cultures without 5-FC, only the samples which were included in the perparation of sequencing libraries
		 are included (some were not sequenced). Culture with added FCY1 mutants are identified "+CDS".
		 All durations are in hours. 
		 Tn_duration columns refer to time interval (hours) between the Tn-1 and Tn samples (T1 is T0-T1, for instance).
		 Tn_OD columns refer to the OD of the corresponding culture when the Tn sample was collected and dilution was performed.
		 Tn_time columns contain the cumulative duration of competition experiment from T0 to Tn.

Table 7: Colony counts for the CRISPR reinsertions of WT F3 and F4 promoter fragments.
		 Data for the reinsertion of the complete F3-F4 region (in background SA0159), which was not shows in the corresponding Supp Fig., is also included.
		 Calculations from colony counts to frequencies of 5-FC resistance and putative FCY1 LOF (both in %) are indicated within the Excel spreadsheet.

** Important note on FCY1 mutations **
In the reads analysis pipeline, all genotypes were assigned a unique identifier. For single mutants, these had the following structure:
{single-letter code for mutation type}-{mutated position}_1nt_{WT nucleotide}to{new nucleotide}. The single letter codes S, I and D were
respectively used for subtitutions, insertions and deletions. For I, the WT nuleotide was written is "-", while for D, the new nucleotide was
represented as "-". In these original codes, positions were defined according to the reference sequence to which all sequencing reads were aligned,
with 1 being the first nucleotide position in this reference. These position were later adjusted to be relative to FCY1 (column FCY1_position), by 
subtracting the position of the first nucleotide of the FCY1 gene. In the case of insertions, the position in the original mutation code was the one
which was immeditely upstream of the inserted nucleotide, such that I-79_1nt_-toT meant that a T had been inserted immediately downstream ("to the
right") of position 79. This was later adjusted in the analysis, for position of S, I and D mutations to be more consistent. A value of 1 was added to
the position of each I mutation, so that it would represent the position of the corresponding mismatch in the alignment, as for S and D. 
* The mutation codes were however not updated *. In intermediate data files, if the adjustement of insertions has been performed, any S and D mutations
should have a higher position (+1) in their mutation code than corresponding I mutations at the same FCY1_position. 

Table 8: Selection coefficients for all F3-F4 single mutants, after removal of outliers (only data from cultures without added FCY1 mutants).
		 The first sheet (Medians_Fig3A) contains the medians (points) shown in Fig 3A and the min/max values per genotype, for s over T0-T2.
		 Adjusted p-values for one-sample t-tests comparing each selection coefficient to 0 are provided. When "_ignored" is added to the label
		 of a column of selection coefficients, the corresponding values have been computed after removal of the four "biased" samples.
		 The min/max values can be used to recompute the mean width of selection coefficient distributions across mutant genotypes, which is
		 informative about the level of noise in the assay (Fig3 and accompanying text).
		 
		 The second sheet (All_reps_no_outs_F3F4) contains all replicate selection coefficients for the same genotypes, after removal of outliers.
		 The "To_ignore" column identifies cultures which were considered biased and thus removed before the computation of the "_ignored" medians.
		 
Table 9: Estimated WT-relative expression levels for all single mutants observed (at least 85 reads) in two or more sort-seq experiments.
		 This includes secondary mutations in the FCY1 gene. Relative expression levels obtained from the calibration of sort-seq scores are shown.
		 As their are based on WT-relative fluorescence intensities, a value of 1.0 by definition corresponds to the WT expression level.
		 The 2.5th, 5th, 50th (median), 95th and 97.5th percentiles are compiled for all genotypes.
		 
Table 10: Transcription factor binding sites (TFBS) identified by YeTFaSCo (http://yetfasco.ccbr.utoronto.ca/) in the F3-F4 region of the FCY1 promoter.
		  The start, midpoint and end of each predicted TFBS is shown relative to the first base of FCY1 (such that the nucleotide immediately upstream
		  of FCY1 is position 0). The "Hit Seq" column contains, for each predicted TFBS, the sequence in the FCY1 promoter which matches the corresponding motif.
		  Predictions were made on the 205 bp of the FCY1 promoter which are included in "ref_F3F4_300_trimmed.fa" (first 205 bp, followed by the first 128 bp of FCY1).
		  Only predicted binding sites which occurred within the F3-F4 region of the promoter were then kept.

Table 11: Expression level (WT-relative fluorescence intensity, measured by flow cytometry) and fitness (from growth curves in [12 uM] 5-FC) for individually reconstructed
		  validation mutants. Relative expression levels and relative fitness (selection coefficients) were obtained from comparisons with a corresponding WT strain constructed
		  in parallel. One of the validation mutants (D1; I-225_1nt_-toA) could not be reconstructed in the FCY1 background.
		  WT-relative fluorescence intensity was measured in three replicates for each variant (42bp-mEGFP background) and the corresponding median, min and max values are
		  reported (Relative_median_exp, Min_rel_exp and Max_rel_exp). Similarly, growth rates was measured for each variant (FCY1 background) from three replicate growth
		  curves, with median, min and max values reported for exponential growth rate (exp_rate, Min_exp_rate, Max_exp_rate), in hours^-1, and for selection
		  coefficients (s_exp, Min_s_exp, Max_s_exp). Columns T_insertion_FCY1 and T_insertion_42-GFP denote whether the corresponding reconstructed mutant featured a single T
		  insertion in the poly-T region of the FCY1 promoter, respectively for reconstructions in the FCY1 and 42bp-mEGFP backgrounds.
